# A distributed neocortical action map associated with reach-to-grasp

**DOI:** 10.1101/2020.01.20.911412

**Authors:** Eros Quarta, Alessandro Scaglione, Jessica Lucchesi, Leonardo Sacconi, Anna Letizia Allegra Mascaro, Francesco Saverio Pavone

## Abstract

Reach-to-Grasp (RtG) is known to be dependent upon neocortical circuits and extensive research has provided insights into how selected neocortical areas contribute to control dexterous movements. Surprisingly, little infor-mation is available on the global neocortical computations underlying RtG in the mouse. Here, we characterized, employing fluorescence wide-field cal-cium imaging, the neocortex-wide dynamics from mice engaging in a RtG task. We demonstrate that, beyond canonical motor regions, several areas, such as the visual and the retrosplenial cortices, also increase their activ-ity levels during successful RtGs. Intriguingly, homologous regions across the ipsilateral hemisphere are also involved. Functional connectivity among areas increases transiently from rest to planning, and decreases during move-ment. Two anti-correlated neocortical networks emerged during movement. At variance, neural activity levels scale linearly with kinematics measures of successful RtGs in secondary motor areas. Our findings establish the coex-istence of distributed and localized neocortical dynamics for efficient control of complex movements.

**SIGNIFICANCE STATEMENT:** In mammals, including humans, the cerebral cortex is known to be critical for the correct execution of dexterous movements. Despite the importance of the mouse for elucidating the neural circuitry for motor control, its neocortex-wide dynamics during RtG are largely unexplored. We used in-vivo fluores-cence microscopy to characterize the neural activity across the neocortex as mice performed a reach-to-grasp task. We show that for such complex movements, a large network of neocortical areas gets involved, while movement kinematics correlates with neural activity in secondary motor areas. These findings indicate the coexistence, at the mesoscale level, of distributed and localized neocortical dynamics for the execution of fine movements. This study offers a novel view on the neocortical correlates of motor control, with potential implications for neural repair.

## INTRODUCTION

Deciphering how the CNS produces goal-directed movements, such as the reach-to-grasp (RtG), is crucial for expanding our understanding on the neural computations underlying motor ability and more generally, for animal cognition (Bayne et al., 2019). RtG is known to be dependent upon neocortical circuits and intensive research has provided detailed insight into how neural activity from motor areas of the neocortex contribute to the control of movements, (Fritsch and Hitzig, 1870; Evarts, 1968; Georgopoulos et al., 1986; Graziano et al., 2002; Churchland et al., 2012). In the mouse, optical methods, which provide mean for minimal invasive neural recordings, are gradually replacing electrophysiological approaches to investigate the neural bases of motor control, however most investigations to date have focused on segregated neocortical areas (Guo et al., 2015; Estebanez et al., 2017). Nevertheless, how neural systems give rise to behavior cannot be fully comprehended from segregated analysis of its components. Indeed, parallel to the patterns of neural activity at the local level, information processing occurring at the cortico-cortical level is necessary for complex behavior (Peters et al., 2014; Allen et al., 2017, Battaglia-Mayer and Caminiti, 2019). However, despite the growing adoption of this organism for elucidating the neural underpinnings of motor control (Ölveczky, 2011), the neocortex-wide dynamics during RtG remain largely unexplored. Here, we inquired about the topographic organization of neural activity during movement, that is, if during RtG specific, large-scale communication schemes emerge across the neocorti-cal mantle. Movement kinematics are associated with neural activity in the contralateral motor areas, and information about such activity can also be employed to predict several features of the executed movement (Prsa et al., 2017, Li et al., 2019). While growing knowledge reported ipsilateral activity during motor behavior, information on the relationship between RtG kinematics and neural activity across the neocortex is still scanty. Filling this gap could provide an important step forward for the understanding of neural mechanics underlying RtG behavior. To tackle these questions, we developed a novel setup and performed a combined analysis of RtG and mesoscale cortical activity from head-fixed mice. The present characterization of the neural activity patterns provide, to our best knowledge, the first evidences indicating that neocortex-wide activity is a prominent neural signature of RtG movements.

## MATERIALS AND METHODS

### Transgenic mice

All experimental procedures were authorized by the Italian Ministry of Health (Authorization Nr. 127-2018-PR). A total of 6 transgenic mice (C57BL/6J-Tg(Thy1-GCaMP6f)GP5.17Dkim/J, stock nr. 025393, herein referred to as GCaMP6f, Chen et al., 2013) aged between 3-9 months were used in this study. Animals were housed on a 12-hr light/dark cycle with ad libitum water. Mice undergoing behavioral training were food restricted to 80-90% of original body weight by limiting food intake to 2-3 g/day. Animals were monitored and weighted daily and food ratios was increased if necessary. Veterinary staff monitored animals twice a week.

### Surgery

The day before surgery, mice were give a subcutaneous injection of en-rofloxacina (10 mg/kg) and each day thereafter for 3 days, to prevent in-fections. The day of surgery, animals were anesthetized with isofluoruane (3% induction, 1.5% maintenance) and placed on a stereotaxic apparatus (Kopf Instruments; Tujunga, CA, USA). Absence of tail reflex and toe-pinch reflex was tested to confirm that the mouse was adequately anesthetized. Throughout the surgery, temperature was kept constant at 37 C by means of an electric heating pad controlled by a rectal temperature sensor (Stoelting; Wood Dale, IL, USA). To prevent cornea dehydration ophthalmic gel was applied over the eyes. The scalp was first scrubbed with ethanol and betadine, then a depilatory cream was applied to remove hairs, then a few drops of topic agent (Lidocaine Hydrochloride 2%) were applied as analgesic measure. A circular piece of scalp was removed and the anatomical points of reference (bregma and lambda) were stained with a permanent marker. Two semicircular coverslip glasses, one per hemisphere, were glued on the skull above the cortex using a transparent dental cement. A custom-build head post was placed ca. 0.5 mm posterior to lambda. At the end of the surgery, subcutaneous injections of analgesic and anti-inflammatory (carpro-fen, 5mg/kg) drugs were given to facilitate recovery. To this end, 200 ml lactated ringer of the 0.9% saline was given to each mouse at the end of surgery. Mice were allowed to recover at least one week following surgery. During this period animals were given ad libitum food and water.

### Behavioral apparatus

The experimental apparatus (Fig. 1A) comprised a stage (size 300 x 300 x 20 mm, Thorlabs, Inc.) and walls to form an enclosure and secured above a lab jack (MLJ050/M, Thorlabs, Inc.) and was inspired by the work of Guo and collaborators (Guo et al., 2015). The enclosure is equipped with a base for the animal, a perch, a speaker to deliver the auditory cue, and a turn-table attached to a servo-motor equipped with a contactless magnetic rotary en-coder to ensure consistent pellet positioning (MX-28AT) and controlled by a controller board (ArbotiX-M Robocontroller, Trousser Robotics). The servo is controlled via an ad-hoc program. Chocolate pellets (10 mg, 5TUL Puri-fied 10mg pellets catalog id. 1811529, TestDiet) were employed to induce the reach-to-grasp movements, serving thus as stimulus. Interstimulus interval (ISI) was 6 to 20 seconds. The enclosure was placed within a custom-made a sensory-isolated enclosed imaging recording chamber that is kept dark and has been insulated with acoustic foam to further reduce ambient sounds (RS components and woods). The behavioral data were collected at 200 frames/s via a high-speed camera (CM3-U3-13YC-CS, Chameleon3, FLIR) with 2.8-8mm varifocal objective lenses (LENS-30F2-V80CS, Fujinon), placed to the left of the head-fixed animal. A 470 nm LED illuminating the stage was used as light source to create a high-contrast image for processing and quantification. Movies were recorded using the FlyCapture software (FLIR).

**Figure 1:**
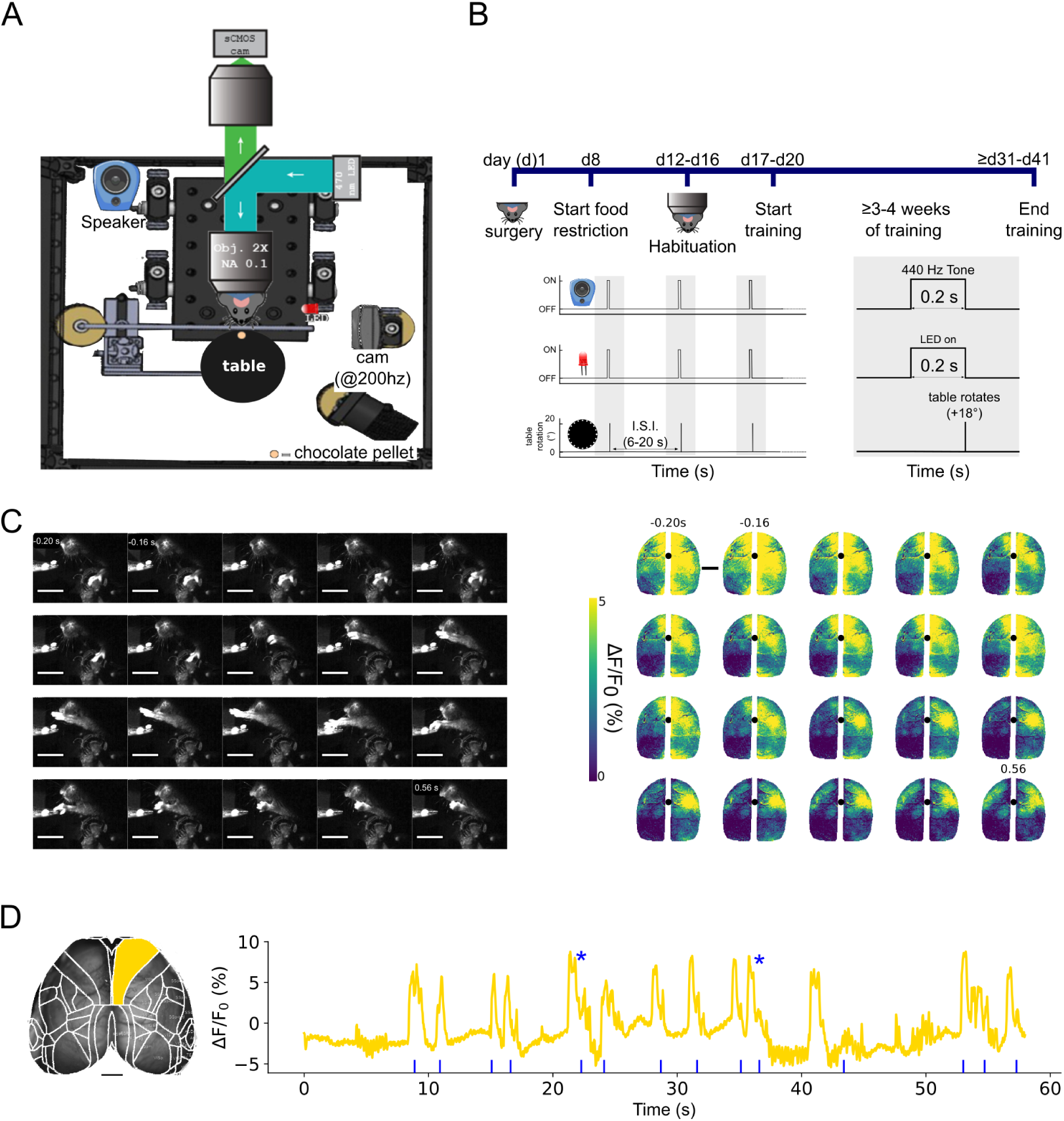
Mesoscale imaging of neocortical dynamics during RtGs. A. Experimental setup. Schematics of the experimental setup, which comprises a custom-made widefield microscope to acquire images, 512 512 pixels, from the intact skull of GCaMP6f transgenic mice with a sampling rate of 25 Hz. Behavioral data were obtained via a high-speed (200 fps) camera placed on the left side of the head-fixed animal. Chocolate pellets were employed to induce voluntary RtG movements. A turn-table attached to a servo-motor was used for pellet delivery. All components of the experimental setup were synchronized by using a common TTL trigger. B. Experimental timeline. Mice underwent a surgery to expose the skull and head-bar implantation. Following recovery from surgery (1 week), mice were food-restricted to 80-90% of their baseline body weight and habituated to the experimental setup for at least 4 days (30 min a day/mouse). After habituation, mice were gently placed under the microscope and head-fixed below the objective. Training consisted of daily sessions (30-45 min/session), and lasted at least 3-4 weeks (5 sessions/week). Each session consisted of several (at least 3) blocks, during which tones (square wave at 440 Hz) preceded by 200ms the rotation of the turntable. Interstimulus interval (ISI) was between 6 to 20 seconds. C. Mesoscale cortical activity during a single RtG movement. Left, image sequence of a mouse performing a successful RtG. Right, image sequence of cortical activity during the same peri-movement epoch (between −0.2 sec and +0.56 sec relative to the movement onset, sampling time 40 msec). Color bar (viridis) represents the range of ΔF/F_0_ (%). In this case, the movement lasts ca. 560 ms. Black dot = bregma. Activity is higher in the medial part of the cortex during early phases of movement. Following grasping phase of the movement, the activity is more localized on the contralateral motor areas. D. Neural activity in right primary motor cortex is associated to repeated RtGs. Left, image showing the typical field of view obtained during recordings. The Allen Mouse Brain Atlas mask was applied on the fluorescence image stacks to parcellate the signals according to the cortical areas of interest. Right, graph depicting the fluorescent and movement signals. F/F_0_ (%) extracted from the right primary motor cortex (gold-colored trace). Onsets of RtGs movements are illustrated by the blue rasterplot. Increase activity in the selected cortical area is observed for each RtG attempt. The asterisks denote successful RtGs.

### Wide-field microscopy

Imaging was performed through the intact skull using a custom-made microscope. The microscope consisted of back-to-back 50 mm f/1.2 camera lenses (Nikon), separated by a FF495-Di03-50.8-D dichroic mirror (Semrock), mounted in a 60 mm cube (Thorlabs). To excite the GCaMP6f indicator, a 470 nm light source (LED, M470L3 Thorlabs, NJ, USA) was deflected by a dichroic filter (DC FF 495-DI02 Semrock, NY, USA) on the objective (TL2X-SAP 2X Super Apochromatic Microscope Objective, 0.1 NA, 56.3 mm WD). The fluorescence signal was selected by a band pass filter (525/50 Semrock, NY, USA) and collected by a CMOS camera (ORCA-Flash 4.0 V2, Hamamatsu). Images were acquired at 25 Hz, with a resolution of 512 *×* 512 pixels with the field of view about 18 *×* 18 mm (depth 16-bit) via the HCImage Live software (Hamamatsu).The microscope thus allows a FOV embracing the entire dorsal prospect of the mouse neocortex. To reduce unwanted light scattering on the mouse eyes (which could serve as visual stimuli, inducing unwanted neural activity), an iris (ID15/M - Mounted Standard Iris, 15 mm Max Aperture, TR75/M Post, Thorlabs Inc.) was placed 1-2 mm above the mouse skull. Imaging sessions were performed for each behavioral training session. The synchronization of the components of the experimental setup occurred via a common hardware trigger signal at the start of each block (see Experimental design for the definition).

### Experimental design

Following recovery from surgery for at least one week, mice were habituated to the experimental setup for 4 or more days (30 min a day/mouse). Then, mice were gently placed under the microscope objective and fixed (Fig. 1B). Each session contained at least 5 blocks, and each block lasted between 65 seconds and 6 minutes, 50 seconds . Here, we analyzed the first 5 days starting from when the mouse performed the first successful RtG. To en-sure consistency, only trajectories from successful RtG, defined as forelimb movements that ended with the pellet eaten, were analyzed.

### Image processing and data analysis

For the movement data, image sequences were loaded onto FIJI (ImageJ) and the metacarpus position was tracked using the manual tracking plugin, in order to obtain the xy coordinates of the successful RtGs. Considering the central aim of the goal-directed task, our investigation was centered on successful RTGs, i.e., those forelimb movements that ended with the pellet being eaten by the mouse. The timing of the movement onset (MoveON) was defined as the frame for which the operator detected a variation in the metacarpus position. The metacarpi coordinates were then employed to reconstruct the movement trajectories and compute the movement kinematics. Here we present the results on movement distance (i.e., arc length), movement duration (in seconds) and the number of movement speed peaks, the latter used as a measure of movement smoothness (Lai et al., 2015). For the neural data, image stacks for each animal collected from different sessions were registered manually using custom made software, by taking into account the bregma and lambda position. To dissect the contribution of each cortical area for RtG, the processed stacks we also registered the cortex to the surface of the Allen Institute Mouse Brain Atlas (www.brain-map.org) projected to our plane of imaging. We further applied a mask to exclude the medial-most areas because they lay over the superior sagittal sinus and therefore the neuronal signal in this region could be more easily affected by hemodynamic artifacts. Areas laying on the most lateral parts of the mouse cortex were also excluded. Also, areas smaller than 100px^2^ were also excluded. This parcellation of the neocortex created 17 areas for each hemisphere, plus three formed by the left, the right and both hemispheres, for a total of 37 areas, whose abbreviations and extended names are: CTX, neocortex; MOs, Secondary motor area; RFA, Rostral Forelimb Area; MOp, Primary motor area; CFA, Caudal Forelimb Area; SSp-bfd, Primary somatosensory area-barrel field; SSp-ll, Primary somatosensory area-lower limb; SSp-n, Primary somatosensory area-nose; SSp-tr, Primary somatosen-sory area-trunk; SSp-ul, Primary somatosensory area-upper limb; SSp-un, Primary somatosensory area-unassigned; RSPagl, Retrosplenial area-lateral agranular part; RSPd, Retrosplenial area-dorsal part; VISrl, Rostrolateral visual area; VISa, Anterior area; VISam, Anteromedial visual area; VISp, Primary visual area; VISpm, posteromedial visual area. Throughout the text and figures, we added the suffix L and R to term cortical areas of the left (ipsilateral with respect to the paw executing the movement) or right (contralateral with respect to the paw executing the movement), respectively (e.g., MOs L, MOs R).

For each block, the image stacks were processed to obtain the estimates of ΔF/F_0_. Briefly, the image stacks were processed considering the equation ΔF/F_0_ = (F-F_0_)/F_0_, where F defines the value of the fluorescence signal in a given moment, F_0_ defines mean fluorescence over each block. Cortical areas were considered associated with movement if their activity during any mo-ment of the time window crossed a threshold defined as the mean+1.5 times the standard deviation of the neural signal occuring during the baseline (-2 to -1 sec with respect to MoveON). The highest value reached in any moment of the 4 second time window was defined as the PeakMax (Fig. 2B). Also, in order to statistically determine if the transaction between the behavioral phases is associated to different neural activity levels, we employed a two-way ANOVA followed, if appropriate, by Tukeys HSD Post-Hoc tests. To evaluate the functional connectivity (FC) between the areas during RtGs, cross-correlations between pairs of areas for each behavioral phase were computed. The fluorescent time series where chunked according to the behavioral phase of interest and are presented as cross-correlation matrices (CCM). We investigated the network membership of these regions by using a hierarchical clustering algorithm (HCA) that was based on the Wards linkage method (Ward, 1963), which minimizes the variant between the clusters. The prox-imity was then interpreted as an indirect measure of FC and represented graphically by means of dendrograms. All data are reported as *mean ± SD* if not stated otherwise. Sample size and appropriate statistical analyses are specified in each figure legend. Statistical significance was defined as *α <*0.05 if not stated otherwise. Multiple comparisons were corrected for by Tukey HSD or Bonferroni correction. No statistical methods were employed to pre-determine sample size. All statistical analyses and related figures were created using either Python or R, and were assembled in Inkscape.

**Figure 2:**
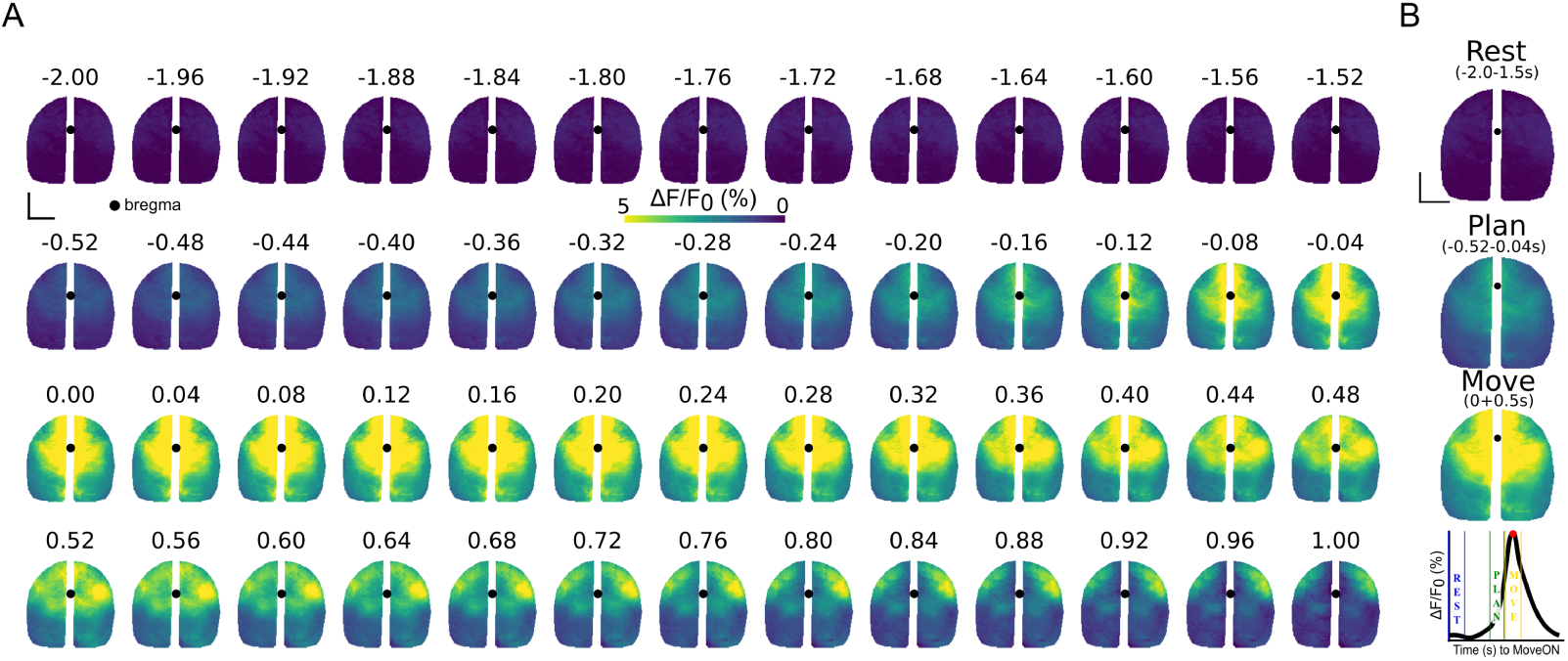
Single-pixel analysis of neocortex-wide activity during RtGs. A. Perievent (-2+2 seconds with respect to MoveON) image sequence. Images display the average signal obtained from aligned stacks of all subjects, for all peri-movement epochs (n=113 movements, 6 mice). On average, activity increases before movement onset in medial regions of the cortex and spreads medio-laterally thereafter. This increase involves areas of both hemispheres. During a latter phase of movement execution we observed a partial bias toward the contralateral hemisphere, possibly associated with the grasping subcomponent of the movement. Black dot = bregma. Scale bar = 2 mm. Color bar (viridis) represents the range of ΔF/F_0_ (%). B. Top, Averaged activity across the behavioral phases rest (-2,-1.5s relative to MoveON), plan (-0.5,-0.04s), and move (0,0.5s). Scale bar = 2 mm. Bottom, cartoon of calcium transient. The vertical lines in the graph depict the division of the investigated behavioral phases. Red dot = PeakMax

## RESULTS

### Wide-field calcium imaging during RtG

We conceived that for a mouse to perform complex, goal-directed movements such as RtG a specific neural network within the neocortex could be recruited. To gain information about such dynamics, our setup integrates a large field of view microscope to simultaneously image the entirety of the neocortical surface and a behavioral apparatus for inducing, delivering and videotaping the RtGs (Fig. 1A). Mice were trained to perform the task while head-fixed under the microscope (Fig. 1B). Once trained, animals executed RtG movements to eat chocolate pellets that were placed on a rotating table placed in front of them (Fig. 1C, left). The time-aligned image stacks obtained from the camera and from the microscope allowed the monitoring of neural activity across functional areas of the neocortex, throughout the behavioral task (Fig. 1C and D).

### Neural activity increases among several areas spanning both hemispheres during reach-to-grasp behavior

Single-pixel analysis of cortex-wide activity indicate that on average, for all movements and for all subjects, the neural activity increases throughout the whole cortical mantle between -2 to -1.5 and between -0.5 to -0.04 seconds with respect to MoveON (Fig. 2A). According to this observation, we analyzed the different contributions of cortical activation during three temporal lapses, namely rest (2 to 1.5 sec before MoveON), plan (0.5 to 0.04 sec before MoveON), and move (0 to 0.5 sec after MoveON, Fig. 2B-C), collectively termed behavioral states or behavioral phases (Scherberger et al., 2005; Hwang and Andersen, 2009; Schieman et al,. 2015). During movement execution, bilateral activation in more rostral areas becomes apparent, while during the latter phase of movement execution a more pronounced increase in neural activity levels emerges in the the contralateral hemisphere (Fig. 2A).

To characterize the contribution of functional areas across the neocortex during RtG, we parcelated the acquired image stacks according to the Allen Institute Mouse Brain Atlas (Fig. 3A, left). The resulting parcellation allowed to extract the calcium signal for each neocortical area. The increase in the levels of neural activity encompassing whole neocortical mantle is observed also on each hemisphere separately (Fig. 3B). Several areas displayed a vigorous increase in neural activity levels with movement execution, including the motor areas and the retrosplenial cortices of both hemispheres, while other areas, such as the barrel field (SSp-bfd), displayed a more modest increase in activity (Fig. 3C-D). Overall, the activity levels across homologous areas of both hemispheres, including motor, somatosensory, retrosplenial, and visual areas appeared comparable.

**Figure 3:**
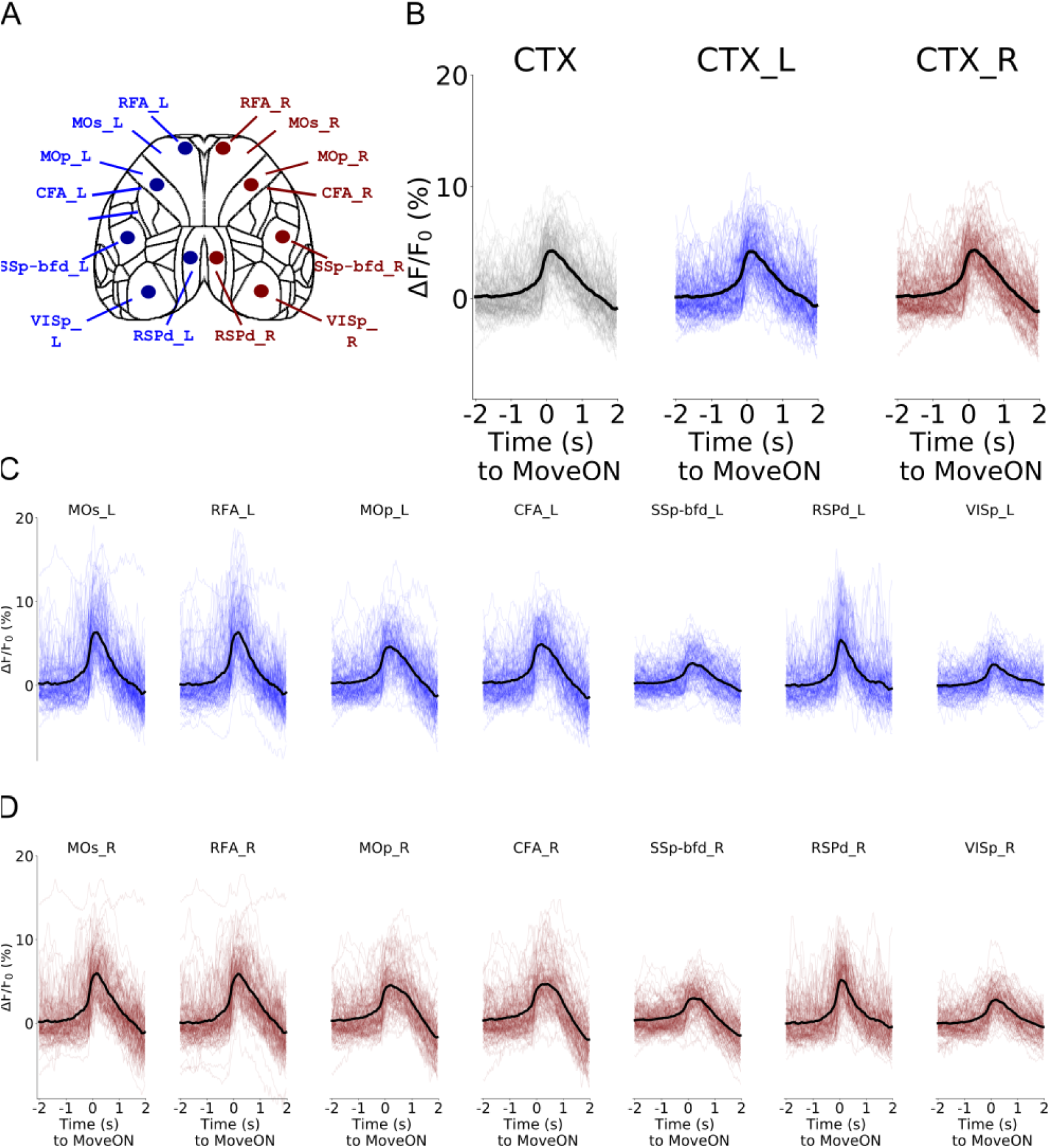
Widespread increase of neural activity during RtGs. A. Illustration of the areas from which the traces displayed in panel B were obtained, which include also the bilateral Rostral Forelimb Area and the Caudal Forelimb Area. Areas and subsequent traces are color-coded accordingly to the left (blue), and right (red) hemisphere. Neural activity traces refer to the peri-movements epochs (n=113 movements, 6 mice). B. ΔF/F_0_(%) activity across blocks (shaded lines) and averaged value (thicker, black line) for the whole cortex, the left and the right hemisphere, for a time window of −2,+2 sec around the movement onset (MoveON). C. Single and mean (thicker, black lines) traces of neural activity for a selection of 8 areas across the two neocortical hemispheres. The MOs, RFA, MOp, CFA and RSPd, and to a lesser extend the SSp-bfd and the VISp, display increased activity levels during RtGs.

### Calcium signals from neocortical areas increases before movement onset

To quantify our observations, we compared the temporal profile of the parcellated fluorescent calcium signal across the perievent time window. All areas emerged as associated to movement, with the rostral-most areas displaying the highest PeakMax and other areas spread across both hemispheres, forming a distributed cortical network associated with reach-to-grasp movements (Fig. 4A). To determine the temporal involvement of cortical areas during the task we computed the latency necessary to reach the 50% and the 100% of the PeakMax (latency to T-halfMax and latency to PeakMax, respectively) in each of the responsive areas (namely the ones that showed activity levels higher than threshold, as described in the materials and methods, 37 cortical areas in total).

**Figure 4:**
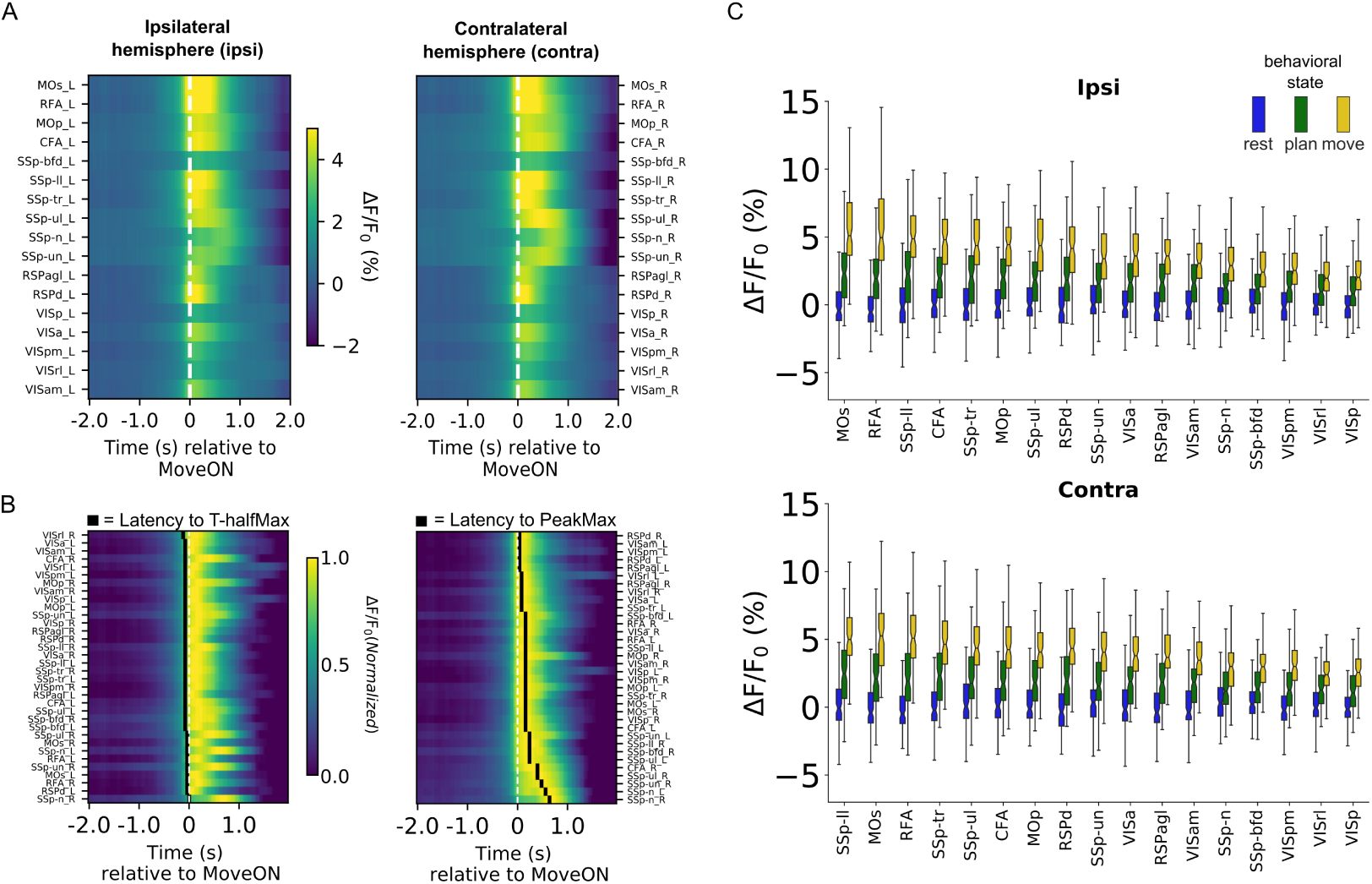
Bilateral and state-associated neural dynamics emerge during RtG. A. Color plots showing the mean activity for all neocortical areas, separated for each hemisphere (ipsi = left, ipsilateral hemisphere, on the left; contra = right, contralateral hemisphere, on the right side). Several homologous areas across the hemispheres exhibit increased activity levels. Colorbar reflects the levels of F/F0 (%) activity. B. Color raster plots showing the mean, normalized activity for the movementassociated areas. Left, the areas in the plot are ranked by latency to reach T-half max%, in ascending order (areas requiring less time to reach the T-half max% are placed higher in the plot). The black squares indicate the latency to T-half max% for each area. Right, the areas in the plot are ranked by latency to reach the PeakMax, in ascending order (areas requiring less time to reach their PeakMax are placed higher in the plot). The black squares indicate the latency to T-half max% for each area. D. Emergence of a state-associated increase of activity levels across cortical areas. Notched boxplots (upper and lower whiskers refer to maximal and minimal value, respectively, excluding outliers) indicate the activity levels for all movement-responsive areas, ranked based on PeakMax, in a descending order from left to right. Left, movement-responsive areas within ipsilateral hemisphere. Bottom, movement-responsive areas within contralateral hemisphere.

We observed that, across the responsive areas, the T-halfMax was reached before movement onset, which indicates that the increases in the amplitude levels of the fluorescent signal anticipate the movement onset (Fig. 4B, left), at variance with the latency to reach PeakMax, occuring mostly after the movement onset (Fig. 4C, right). While the T-halfMax was comparable across areas, the latency to reach PeakMax indicated a higher variability, with so-matosensory areas and the contralateral CFA displaying a delay to reach the PeakMax compared to other areas. ANOVA revealed that the levels of neural activity were significantly affected by the cortical area [F(33,11424)=19.214, p*≤*0.0001], by the behavioral [F(2,11424)=2920.965, p*≤*0.0001)], and by the interaction of these two factors [F(66,11424)]=6.382, p*≤*0.0001)].

Tukey’s HSD Post-Hoc results indicate statistically significant increases from phase rest to movement for several distributed movement-associated areas, including the MOs, the RFA, the MOp, the CFA, the RSPd, the somatosen-sory primary cortex upper limb (SSp-ul), and the visual areas. Mean activity levels were found to be already different when comparing the rest vs the planning phase, e.g., for the MOs, the RFA, the MOp, the CFA, the RSPd and the SSp-ul, but were not found to be different for the comparisons involving the primary visual areas (for all pairwise comparisons, see supplementary table 1). These findings suggest that a global network of areas is activated during movement execution, with the majority of them increasing their levels of activity before movement onset.

### Motor planning is associated with a global increase in FC

Considering the increase in neural activity levels across the areas from rest to movement, we next asked whether such variation was associated with modulation of functional connectivity (FC) parameters across areas, and whether FC was modulated throughout behavioural phases. The HCA revealed that, in the context of the planning phase, all functional areas tend to form a single functional aggregate highlighted by the fact that they meet lower on the dendrogram compared to the rest phase (Fig. 5A-B, top row). This tendency is illustrated by the CCM (Fig. 5B, bottom row), indicating an increase in the correlation coefficients in the transition from the rest phase to the plan phase (0.5-0.04s with respect to MoveON, Fig. 5B, left and center, respectively).

**Figure 5:**
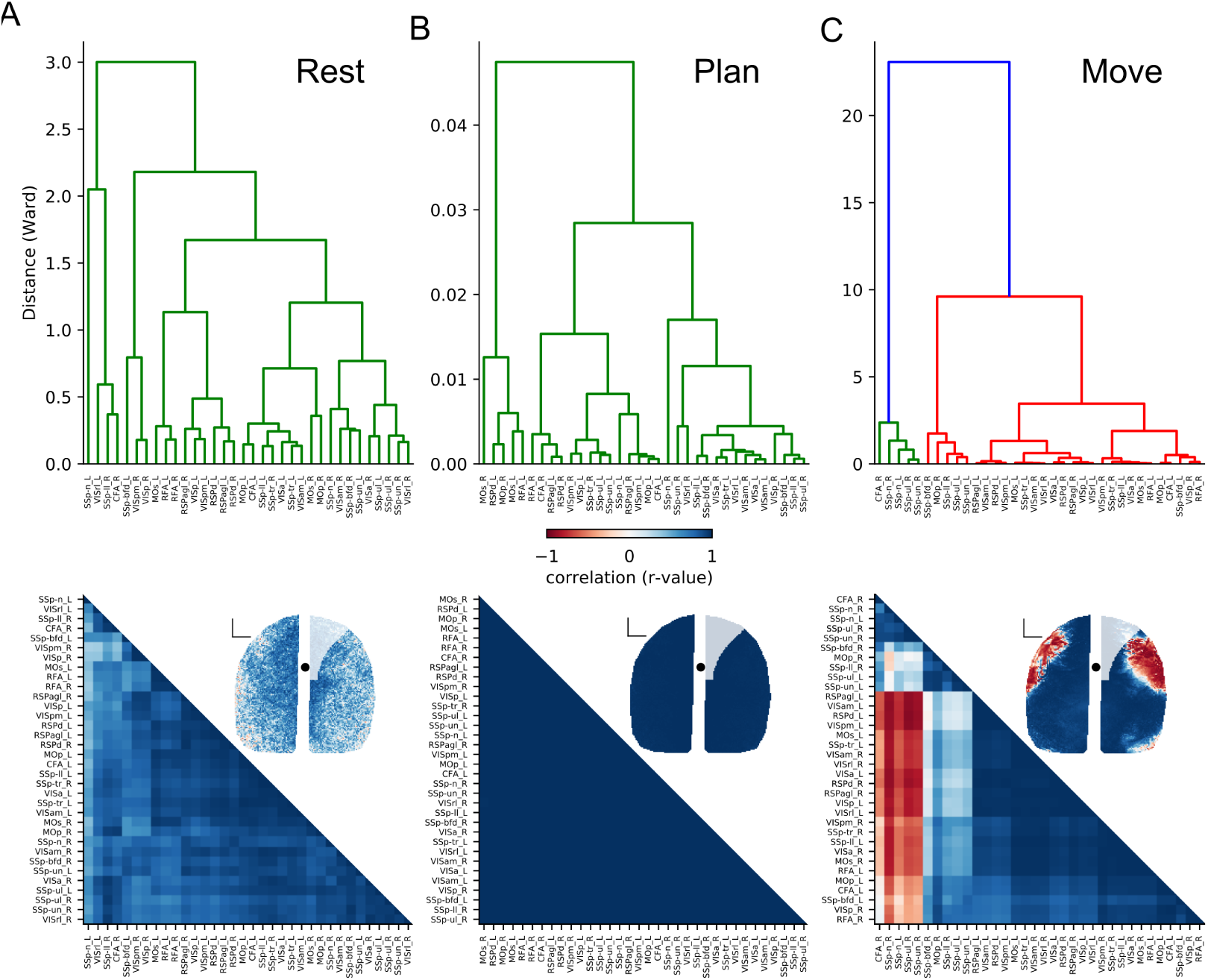
Motor planning is associated with transient increases in FC. The functional connectivity (FC) was analyzed by means of cross-correlation matrices (CCM), for each of the three behavioral phases and are shown in columns from left to right. Cross-correlation coefficient in the matrices are shown from +1 (in blue) to -1 (in red). The order of appereance of each cortical area in the CCM is based upon area position along the dendrogram. A, B. Strong increase of FC during planning with respect to rest. This increase is transient, as during the movement phase there is an abrupt shift towards rearrangement of the functional connectivity, shown by high anti-correlations between selected areas. C. During the RtG execution, some areas maintain highly positive cross-correlation values. A marked anticorrelation arises between the associative and the executive clusters during movement. In panels A-C Seed-pixel analysis (SCA) across each behavioral phase is . The ipsilateral MOs (B), MOp (C), and RSPd (D) were used as seeds, respectively. SCA indicate that from rest to plan vs the activity is highly correlated across the neocortex. This analysis also highlights that during movement execution (move) two anticorrelated networks emerge across both hemispheres. The anticorrelated network overlaps, at least partially, with the canonical coordinates of CFA. Correlation coefficients (CC, -1 denote anticorrelation, +1 denote correlation) is color-coded (jet colormap). Black dots = bregma scalebar = 1mm.

Upon movement execution, the FC transitions to the opposite direction, i.e., there is a decrease in the correlation coefficients (Fig. 5C, right).

### Two anticorrelated networks emerge during movement execution

We next sought to determine whether such scheme of communication is maintained in a context of single-pixel analysis. To this end, the contralateral MOs was used as seed and the cross-correlation of every pixel with each individual area was computed (Fig. 5A-C, bottom row). The transition from resting to planning was associated with a robust increase in the cross-correlation (Fig. 5A-B, bottom row). Upon movement execution, two anticorrelated networks emerged for all seeds (Fig. 5A-B, bottom row). The first network contained areas associated with higher-level transformation, including all subdivisions of the retrosplenial cortex, the secondary motor areas of both hemispheres (MOs and RFA) as well as the visual areas (which could be regarded as the sensorimotor transformation network). The second network includes almost exclusively subdivisions of the primary sensory cortex, the primary motor cortices of both hemispheres (and could viewed as the executive network). Our approach provides evidence for the existence of segregated functional networks emerging during motor execution. Overall, these FC results suggest that the cortico-cortical communication scheme during goal-directed forelimb movements can be decomposed into two parts: the earlier phase, with high increase in cross-correlation, and a latter, where two anti-correlated networks emerge.

### Neural activity levels in the MOs and RFA scale linearly with movement kinematics

Considering the increase in activity observed across the neocortical areas, we sought to determine the extend of association between neural activity and movement parameters. To this end we first reconstructed the spatial trajectories for the successful RtGs and computed the movement metrics (Fig. 6A-C). The movements distance was 52.8 *±* 18.0mm (Fig. 6D), the movements lasted 0.7 *±* 0.2 seconds (Fig. 6E), while the number of speed peaks was 5.4 *±* 2.7 (Fig. 6F).

**Figure 6:**
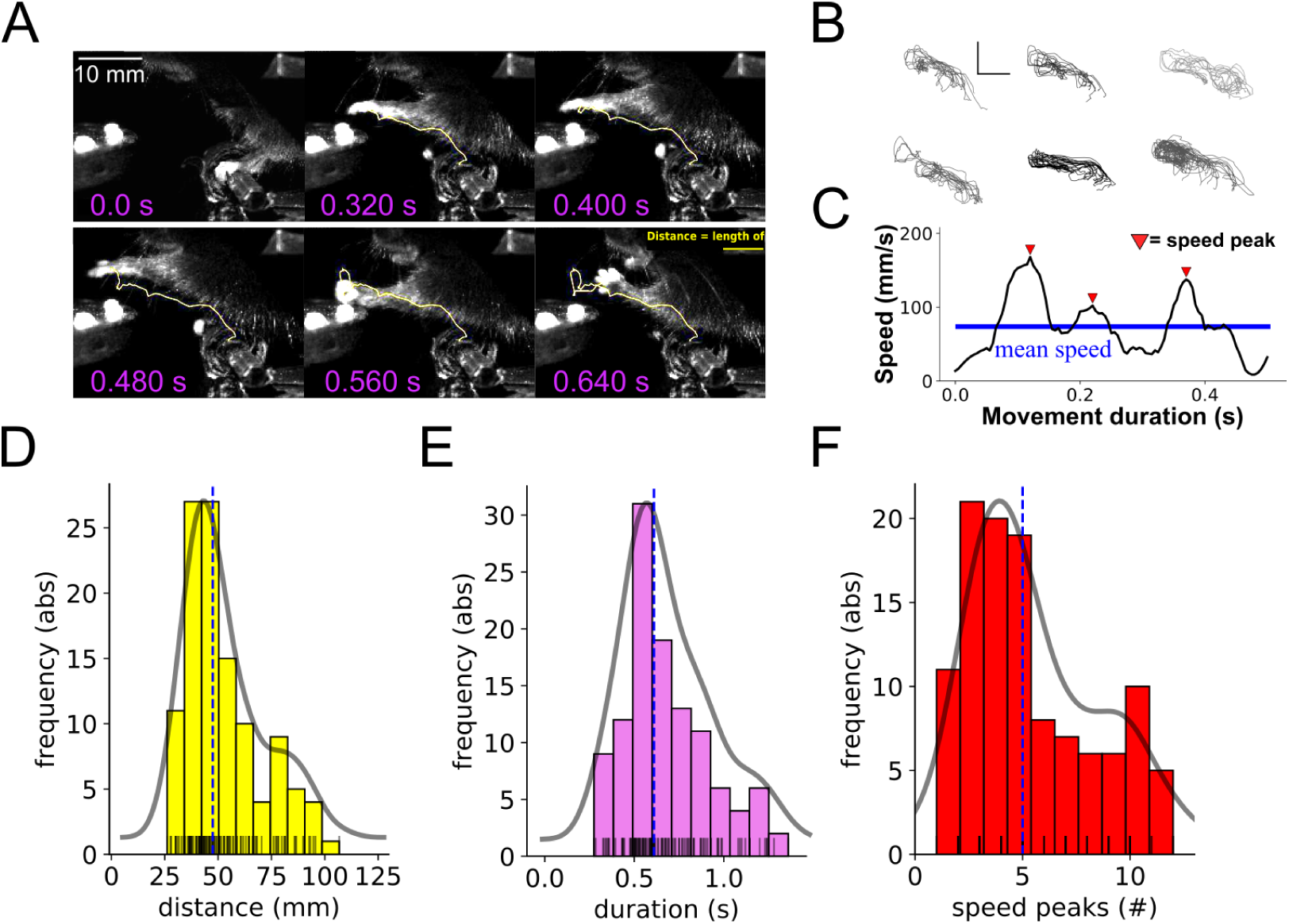
Kinematic trajectories of successful RTGs A. Example of RtG movement tracked for reconstruction and kinematic analysis. A. In this stereotypical reach-to-grasp sequence, the yellow line in each image represent the path covered by the metacarpus. B. Reconstructed reach-to-grasp trajectories of left-paw movements for each animal. Trajectories of successful movements are shown, separated for each animal. Scalebar = 10 mm for both directions. C. Speed profile reconstruction from a successful RtG movement. Speed peaks exceeding the mean speed (horizontal blue line) are used to compute the movement smoothness (red triangles). C. Distribution of the movement distance. Histogram (bin size = 10) showing the absolute frequency distribution for the distances (arc length) covered by the paw to reach the chocolate pellet and eat it. The vertical line illustrate the median value (47.47 mm), the shaded curve indicate the density distribution, lines of the rug plots (bottom) illustrate the raw distribution for each movement. D. Distribution of the movement duration. Histogram (bin size = 10) showing the absolute frequency distribution for the movement durations. The vertical line illustrate the median value (0.61 s), the shaded curve indicate the density distribution, lines of the rug plots (bottom) illustrate the raw distribution for each movement. D. Distribution of the number of speed peaks. Histogram (bin size = 10) showing the absolute frequency distribution for the speed peaks. The vertical line illustrate the median value (5), the shaded curve indicate the density distribution, lines of the rug plots (bottom) illustrate the raw distribution for each movement (note: the speed peaks being computed as natural numbers, overlap frequently in the rug plot).

Since during movement execution we observed the emergence of differentiated activation among areas, we asked to what extent each area contributed to specif kinematic parameter of RtGs movement.

To this end, we performed correlation analysis between PeakMax in selected areas of the neocortex and the extracted parameters of movement kinematics: movement distance, movement durations and movement smoothness (Fig. 7). A moderate association between the distance covered and the Peak Max in the MOs of both hemispheres (ipsi: r=0.42; p=4.5e-06, contra: r=0.42; p=4.2e-06) as well as the RFA (ipsi: r=0.44; p=1.2e-06, contra: r=0.37; p=4.8e-05) was found. Interestingly, in the MOp or the CFA the results indicate that statistical significance was not reached, not for MOp (ipsi: r=0.24; p=0.011, contra: r=0.13; p=0.16) nor for CFA (ipsi: r=0.23;p=0.013, contra: r=0.15; p=0.12), suggesting that in these areas the PeakMax may not be associated with movement distance. No evidence for linear relationship among the same parameter was found in the RSPd and in the VISpm (Fig. 7A). The same pattern was observed when considering the relationship between PeakMax and movement duration. Indeed, the MOs (ipsi: r=0.4; p=1.5e-06, contra: r=0.48; p=7.8e-08) and the RFA (ipsi: r=0.45; p=4.8e-07, contra: r=0.4; p=1.2e-05) were the only areas whose activity was robustly associated with this movement metric (Fig. 7B). We next investigated the relationship between the neural activity levels and the movement smoothness (e.g., Lai et al., 2015). Very high statistical significance for moderate positive association between PeakMax and movement smoothness was found in the MOs (ipsi: r=0.43; p=2.4e-06, contra: r=0.41; p=7.8e-06) and in the RFA (ipsi: r=0.41; p=6.9e-06, contra: r=0.43; p=1.9e-06). No evidence for linear relationship was found in the other areas, including the RSPd and the VISPm (Fig. 7C). Therefore, while activity level increase neocortex-wide, secondary motor areas contained the most information related to kinematic parameters of RtGs.

**Figure 7:**
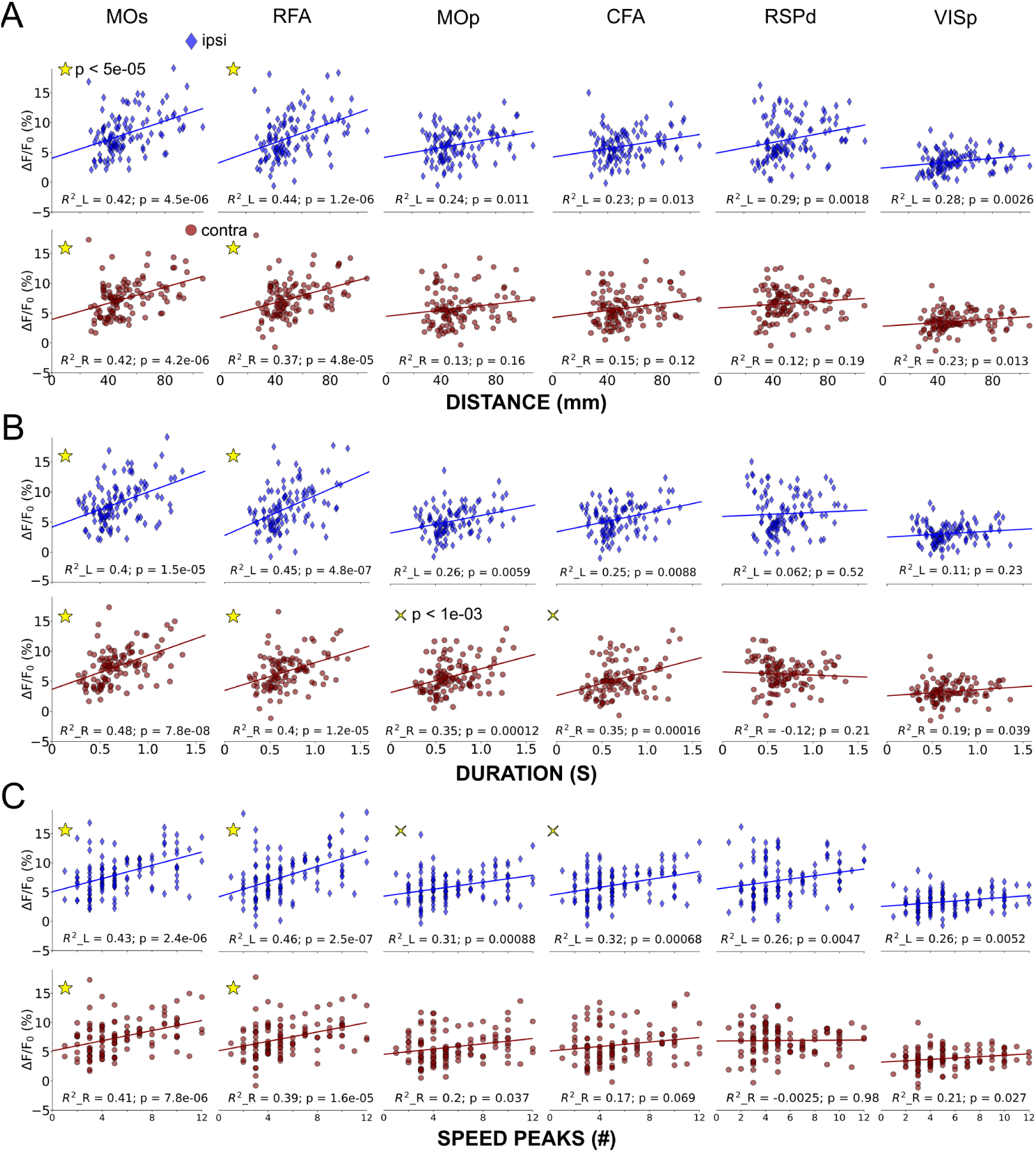
Neural activity levels in both hemispheres scale linearly with movement kinematics. For all panels, the data from the ipsilateral cortical areas are shown as blue diamonds, and the data from the contralateral cortical areas are illustrated as red circles. The PeakMax occuring for each movement within a temporal window of −0.2+0.8 seconds from MoveON was extracted, for each area across both hemispheres and correlate to movement distance (A), movement duration (B), and movement smoothness (C). For the correlation analysis of each pair of parameters, we set a Bonferroni-adjusted significance level of *α<*5 × 10^-5^. Panels with star denote statistical significance, panels with cross denote statistically non-significant trends. The scatterplots show the relationship between PeakMax and the distance covered, across selected homologous areas of the ipsi- and contralateral hemisphere. PeakMax observed in the RFA and MOs of the ipsi- and contralateral hemisphere were found to be positively correlated with the distance covered during the RtGs. This relationship was not observed in the MOp, in the CFA nor in the RSPd nor in the VISp. B. The scatterplots show the relationship between PeakMax and the movement duration, across selected homologous areas of the ipsi- and contralateral hemisphere. PeakMax observed in the bilateral RFA and MOs were found to be positively correlated with the duration of the RtGs. This relationship was not observed in the MOp, in the CFA nor in the RSPd nor in the VISp. C. The scatterplots show the relationship between PeakMax and the speed peaks, across selected homologous areas of the ipsi- and contralateral hemisphere. PeakMax observed in the bilateral RFA and MOs were found to be positively correlated with the speed peaks of the RtGs. This relationship was not observed in the MOp, in the CFA nor in the RSPd nor in the VISp.

## DISCUSSION

In the present work, we report results from experiments in which we employed a wide-field fluorescent imaging approach to provide an unprecedented view of the mesoscale level neocortical dynamics from mice performing a voluntary reach-and-grasp task. To the best of our knowledge, this is the first reported step towards an all-optical characterization of neocortex-wide neural dynamics in rodents performing voluntary RtG movements. We found that during RtG, a distributed network of cortical areas, beyond the regions classically associated with the production of skilled movements, displays increased activity levels. Also, the increased activity is observed in both hemispheres, a finding quite unexpected from a unilateral motor task. The FC among those movement-associated areas undergo a brief (*<*1 sec), transition, i.e., towards an increased connectivity from rest to the planning phase, and towards a decreased connectivity from planning to movement execution. Our analyses indicate the emergence of functional clusters during the movement execution. At variance with the global communication scheme during successful RtG, the internal representation of movement parameters appears localized. Indeed, a robust association between the levels of neural activity and selected RtGs kinematics was found primarily in the secondary motor areas (MOs and RFA of the contra- and ipsilateral hemispheres).

In rodents, it was shown that areas such as the contralateral RFA are both necessary and sufficient to elicit goal-directed and it was also demonstrated that different areas subserve discrete steps of such behaviors (Guo et al., 2015; Morandell and Huber 2017; Wang et al., 2017; Galiñanes et al., 2018). However, these studies were focusing on sensorimotor areas and may have missed important information on the dynamics taking place in other cortical areas during RtG. Our work expands upon this line of research, by providing supporting additional evidence for an involvement of sensorimotor areas during RtG and, more importantly, by directly observing all areas of the neocortex, hence providing an integrative framework not reported previously. The global communication scheme found in the present work is consistent with recent reports on other types of behavioral tasks (Goard et al., 2016; Kyriakatos et al., 2016; Wekselblatt et al., 2016; Allen et al., 2017; Makino et al., 2017; Salkoff et al., 2019; Stringer et al., 2019, Musall et al., 2019). However to our best knowledge, similar approaches were not yet applied to investigate neural dynamics underlying the control of unilateral forelimb movements requiring to locate, reach and grasp for an object. Also, a direct correlative analysis between neocortex-wide neural activity and RtG kinematics was still largely unexplored. Our work provides thus the first direct evidence indicating that while neural activity increases globally during RtG, neuro-kinematic correlations are significant only for the secondary motor areas, suggesting that movement planning and execution may involve large neural networks across the neocortex while encoding of movement parameters occurs in segregated areas. Our work also suggests that during successful reach-to-grasp movements the information processing occurring in the neocortex for multijoint movements is distributed, at least when contextualized to our imaging approach, the temporal windows analyzed and the cortical parcellation employed. Our results point the attention also to areas beyond the MOs, the MOp and their subdivisions (RFA and CFA) towards other territories such as the RSC, the SSp and the visual areas, which were previously not recorded and studied simultaneously within the context of the RtG. For the RSC, while its participation to RtG in the mouse is demonstrated here for the first time, these findings are supported by anatomical evidence, indicating monosynaptic communication between the RSC and the MOs (Yamawaki et al., 2016). At the functional level, the contribution of RSC may be explained by its known involvement in the processing of visual information necessary to locate the target relavant for the behavioral tasks (Czajkowski et al., 2014). There are some aspects that needs to be considered when interpreting the results. The one-photon, wide-field calcium imaging approach is sensitive to light scattering potentially leading to artifacts in the signals extracted from the cortical areas, which could increase the risk of erroneously consider as an area as movement-associated (Yang and Yuste, 2017). Such possibility is reduced by the fact that areas even in close proximity (for example the barrel fields vs the upper limb portion of the somatosensory cortex) do not display the same levels of activity (Fig. 5 A). The analyzed neural activity is inferred from the fluctuations of fluorescence intensity throughout time, a signal composed also by non-neuronal sources, notoriously of hemodynamic origin, which represents another artifact that needs to be taken into account (Ma et al., 2016). However, the hemodynamic component of the signal has a much slower rise and decay time with respect to the neuronal signal. This temporal feature of the hemodynamic component of the GCaMP6f signal reduces its contribution relative to the results reported here (Scott et al., 2018). Even considering such limitations, our findings remains unexpected and provide experimental evidence to put into questions some dogmas of the classical literature in motor control (Omrani et al., 2017), at least when investigating the modus operandi of rodent cortex, adding experimental evidence to the relatively new idea of ipsilateral representation of movements parameters, which is also a computational feature of the human neocortex and is increasingly getting attention in the motor control community (Bundy et al., 2018; Bundy and Leuthardt, 2019). In rodents, a study restricted to the posterior parietal cortex recently reported evidence for ipsilateral control of limb movement (Soma et al., 2019). We concede that the number of neocortical clusters emerging during movement could be influenced by the Allen Institute Mouse Brain Atlas parcellation of the cortex, which may introduce spurious segregations beyond those that are physiologically present. Importantly, the overlap between the results obtained from the SCA and the CCM, supports the existence of two neocortical networks during voluntary movement motor control.

Overall, these associations suggest the secondary motor areas could encode for the movement duration, movement distance, and movement smoothness, encouraging future investigations toward decoding of movement kinematics from neural data. Decoding approaches have been long and very well established for electrophysiological studies of motor control, and more recently applied to fluorescent imaging data of neural activity (Prsa et al., 2017; Li et al., 2019). Our findings extend these studies by providing evidence for correlation between large scale, area-level activity, and specific metrics of movement. Within the context of motor control, our findings also encourage to study the role of areas found to be active during movement (such as the visual areas) commonly disregarded in the association to forelimb movement, at the cellular-level (Galiñanes et al., 2018, Karandell and Huber, 2017, Ebina et al., 2018). In the future, it would also be relevant to assess the neocortical dynamics during RtG in a freely moving condition or during performance refinement, considering that cortical areas undergo functional reorganization during learning (Makino et al., 2017; Whishaw et al., 2017; Bollu et al., 2019; Hwang et al., 2019). In sum, our investigation identified a global network of neocortical area associated to successful RtG execution. Since we observed that several areas across both hemispheres of the cerebral cortex display movement associated activity, we asked whether this activity could be associated with metrics of RtGs movements. We found that among the responsive areas, the secondary motor areas of both hemispheres displayed the most robust association between the neural and the movement kinematics. Previous work showed that these areas is encodes performance rate in a licking task (Salkoff et al., 2019). Our current results suggest that the secondary motor areas could encode for the forelimb movement duration, movement distance, and movement smoothness, encouraging future investigations toward decoding of movement kinematics from neural data. Decoding approaches have been long and very well established for electrophysiological studies of motor control, and more recently applied to fluorescent imaging data of neural activity (Prsa et al., 2017; Li et al., 2019; Salkoff et al., 2019). Our findings extend these studies by providing evidence for correlation between large scale, area-level activity, and specific metrics of RtG movements. Within the context of motor control, our findings also encourage to study the role of areas found to be active during movement (such as the visual areas) commonly disregarded in the association to forelimb movement, at the cellular-level (Galianes et al., 2018, Karandell and Huber, 2017, Ebina et al., 2018). In the future, it would also be relevant to assess the neocortical dynamics during RtG in a freely moving condition or during performance refinement (Makino et al., 2017; Whishaw et al., 2017; Bollu et al., 2019). In sum, our investigation identified a global network of neocortical areas beyond those previously known to be associated with goal-directed arm movements and suggest that the operating mode of the mouse neocortex underlying unilateral goal-directed movements involves distant communication across bothhemispheres, while encoding of movement parameters occurs locally, and, in the context of our investigations, mostly in the secondary motor areas.

## Acknowledgments

We would like to thank the staff of the mechanical and electronic workshops of LENS for their assistance, as well as the members of the Pavone Lab for helpful discussion and suggestions. We would like to thank Costanza Campaioli and Chiara Caldini for their help in developing earlier versions of the experimental setup. Finally, we would like to thank Prof. Matteo Caleo for his useful comments on the manuscript. This project has received funding from the the European Union’s Horizon 2020 Research and Innovation Programme under Grant Agreements 785907 (HBP SGA2) and from the Italian Ministry for Education, University, and Research in the framework of the Project FARE MIMIC. Also, this project has received funding from the European Research Council (ERC) under the European Union’s Horizon 2020 Research and Innovation Programme (grant agreement No. 692943).

## Author contributions

EQ conceived the study, developed the experimental setup, performed the surgeries, performed the experiments, wrote the analysis algorithms, performed the analysis, produced the figures and wrote the manuscript. AS wrote the analysis algorithms, performed the analysis; JL participated in the experiments, participated in the analysis; LS developed the setup; ALAM conceived the study, participated in the experiments, participated in the analysis; FSP conceived the study. All authors discussed and iterated on the analysis and its results, and all authors revised the final manuscript.

