## Supplemental Table 1 for "A distributed neocortical action map associated with reach-to-grasp"

**SUPPLEMENTARY MATERIAL**

Table 1:
Statistical comparison of mean $\Delta F/F_0$ across behavioral states
(pairwise post-hoc comparisons, p-values $< 0.001$ are shown in bold)

| area state vs area state | $pvalue_{adj}$ |
|---|---|
| CFA_R move vs CFA_L move | 1 |
| MOp_L move vs CFA_L move | 1 |
| MOp_R move vs CFA_L move | 1 |
| MOs_L move vs CFA_L move | 0.33 |
| MOs_R move vs CFA_L move | 1 |
| RFA_L move vs CFA_L move | 0.44 |
| RFA_R move vs CFA_L move | 0.99 |
| RSPagl_L move vs CFA_L move | 0.63 |
| RSPagl_R move vs CFA_L move | 1 |
| RSPd_L move vs CFA_L move | 1 |
| RSPd_R move vs CFA_L move | 1 |
| SSp-bfd_L move vs CFA_L move | **0** |
| SSp-bfd_R move vs CFA_L move | **3.6e-06** |
| SSp-ll_L move vs CFA_L move | 1 |
| SSp-ll_R move vs CFA_L move | 1 |
| SSp-n_L move vs CFA_L move | **2.6e-05** |
| SSp-n_R move vs CFA_L move | **0.00011** |
| SSp-tr_L move vs CFA_L move | 1 |
| SSp-tr_R move vs CFA_L move | 1 |
| SSp-ul_L move vs CFA_L move | 1 |
| SSp-ul_R move vs CFA_L move | 1 |
| SSp-un_L move vs CFA_L move | 0.21 |
| SSp-un_R move vs CFA_L move | 1 |
| VISa_L move vs CFA_L move | 0.34 |
| VISam_L move vs CFA_L move | **0.0022** |
| VISam_R move vs CFA_L move | 0.41 |
| VISa_R move vs CFA_L move | 1 |
| VISp_L move vs CFA_L move | **0** |
| VISpm_L move vs CFA_L move | **1.6e-09** |
| VISpm_R move vs CFA_L move | **0.00051** |
| VISp_R move vs CFA_L move | **4.5e-09** |
| VISrl_L move vs CFA_L move | **0** |
| VISrl_R move vs CFA_L move | **0** |
| CFA_L plan vs CFA_L move | **0** |
| CFA_R plan vs CFA_L move | **0** |
| MOp_L plan vs CFA_L move | **0** |

| | |
|---|---|
| MOp_R plan vs CFA_L move | **0** |
| MOs_L plan vs CFA_L move | **1.4e-08** |
| MOs_R plan vs CFA_L move | **0** |
| RFA_L plan vs CFA_L move | **3.3e-09** |
| RFA_R plan vs CFA_L move | **0** |
| RSPagl_L plan vs CFA_L move | **0** |
| RSPagl_R plan vs CFA_L move | **0** |
| RSPd_L plan vs CFA_L move | **0** |
| RSPd_R plan vs CFA_L move | **0** |
| SSp-bfd_L plan vs CFA_L move | **0** |
| SSp-bfd_R plan vs CFA_L move | **0** |
| SSp-ll_L plan vs CFA_L move | **0** |
| SSp-ll_R plan vs CFA_L move | **2.5e-09** |
| SSp-n_L plan vs CFA_L move | **0** |
| SSp-n_R plan vs CFA_L move | **0** |
| SSp-tr_L plan vs CFA_L move | **0** |
| SSp-tr_R plan vs CFA_L move | **0** |
| SSp-ul_L plan vs CFA_L move | **0** |
| SSp-ul_R plan vs CFA_L move | **0** |
| SSp-un_L plan vs CFA_L move | **0** |
| SSp-un_R plan vs CFA_L move | **0** |
| VISa_L plan vs CFA_L move | **0** |
| VISam_L plan vs CFA_L move | **0** |
| VISam_R plan vs CFA_L move | **0** |
| VISa_R plan vs CFA_L move | **0** |
| VISp_L plan vs CFA_L move | **0** |
| VISpm_L plan vs CFA_L move | **0** |
| VISpm_R plan vs CFA_L move | **0** |
| VISp_R plan vs CFA_L move | **0** |
| VISrl_L plan vs CFA_L move | **0** |
| VISrl_R plan vs CFA_L move | **0** |
| CFA_L rest vs CFA_L move | **0** |
| CFA_R rest vs CFA_L move | **0** |
| MOp_L rest vs CFA_L move | **0** |
| MOp_R rest vs CFA_L move | **0** |
| MOs_L rest vs CFA_L move | **0** |
| MOs_R rest vs CFA_L move | **0** |
| RFA_L rest vs CFA_L move | **0** |
| RFA_R rest vs CFA_L move | **0** |
| RSPagl_L rest vs CFA_L move | **0** |
| RSPagl_R rest vs CFA_L move | **0** |
| RSPd_L rest vs CFA_L move | **0** |
| RSPd_R rest vs CFA_L move | **0** |
| SSp-bfd_L rest vs CFA_L move | **0** |
| SSp-bfd_R rest vs CFA_L move | **0** |
| SSp-ll_L rest vs CFA_L move | **0** |

| | |
|---|---|
| SSp-ll_R rest vs CFA_L move | **0** |
| SSp-n_L rest vs CFA_L move | **0** |
| SSp-n_R rest vs CFA_L move | **0** |
| SSp-tr_L rest vs CFA_L move | **0** |
| SSp-tr_R rest vs CFA_L move | **0** |
| SSp-ul_L rest vs CFA_L move | **0** |
| SSp-ul_R rest vs CFA_L move | **0** |
| SSp-un_L rest vs CFA_L move | **0** |
| SSp-un_R rest vs CFA_L move | **0** |
| VISa_L rest vs CFA_L move | **0** |
| VISam_L rest vs CFA_L move | **0** |
| VISam_R rest vs CFA_L move | **0** |
| VISa_R rest vs CFA_L move | **0** |
| VISp_L rest vs CFA_L move | **0** |
| VISpm_L rest vs CFA_L move | **0** |
| VISpm_R rest vs CFA_L move | **0** |
| VISp_R rest vs CFA_L move | **0** |
| VISrl_L rest vs CFA_L move | **0** |
| VISrl_R rest vs CFA_L move | **0** |
| MOp_L move vs CFA_R move | 1 |
| MOp_R move vs CFA_R move | 1 |
| MOs_L move vs CFA_R move | 0.087 |
| MOs_R move vs CFA_R move | 0.9 |
| RFA_L move vs CFA_R move | 0.13 |
| RFA_R move vs CFA_R move | 0.86 |
| RSPagl_L move vs CFA_R move | 0.94 |
| RSPagl_R move vs CFA_R move | 1 |
| RSPd_L move vs CFA_R move | 1 |
| RSPd_R move vs CFA_R move | 1 |
| SSp-bfd_L move vs CFA_R move | **1.6e-09** |
| SSp-bfd_R move vs CFA_R move | **5.4e-05** |
| SSp-ll_L move vs CFA_R move | 1 |
| SSp-ll_R move vs CFA_R move | 1 |
| SSp-n_L move vs CFA_R move | **0.00033** |
| SSp-n_R move vs CFA_R move | **0.0012** |
| SSp-tr_L move vs CFA_R move | 1 |
| SSp-tr_R move vs CFA_R move | 1 |
| SSp-ul_L move vs CFA_R move | 1 |
| SSp-ul_R move vs CFA_R move | 1 |
| SSp-un_L move vs CFA_R move | 0.59 |
| SSp-un_R move vs CFA_R move | 1 |
| VISa_L move vs CFA_R move | 0.75 |
| VISam_L move vs CFA_R move | 0.017 |
| VISam_R move vs CFA_R move | 0.81 |
| VISa_R move vs CFA_R move | 1 |
| VISp_L move vs CFA_R move | **0** |

| | |
|---|---|
| VISpm_L move vs CFA_R move | **6.2e-08** |
| VISpm_R move vs CFA_R move | **0.0048** |
| VISp_R move vs CFA_R move | **1.3e-07** |
| VISrl_L move vs CFA_R move | **0** |
| VISrl_R move vs CFA_R move | **4.2e-09** |
| CFA_L plan vs CFA_R move | **0** |
| CFA_R plan vs CFA_R move | **0** |
| MOp_L plan vs CFA_R move | **0** |
| MOp_R plan vs CFA_R move | **0** |
| MOs_L plan vs CFA_R move | **3.3e-07** |
| MOs_R plan vs CFA_R move | **2.8e-09** |
| RFA_L plan vs CFA_R move | **1e-07** |
| RFA_R plan vs CFA_R move | **0** |
| RSPagl_L plan vs CFA_R move | **0** |
| RSPagl_R plan vs CFA_R move | **0** |
| RSPd_L plan vs CFA_R move | **0** |
| RSPd_R plan vs CFA_R move | **0** |
| SSp-bfd_L plan vs CFA_R move | **0** |
| SSp-bfd_R plan vs CFA_R move | **0** |
| SSp-ll_L plan vs CFA_R move | **1.5e-09** |
| SSp-ll_R plan vs CFA_R move | **8.4e-08** |
| SSp-n_L plan vs CFA_R move | **0** |
| SSp-n_R plan vs CFA_R move | **0** |
| SSp-tr_L plan vs CFA_R move | **0** |
| SSp-tr_R plan vs CFA_R move | **1.5e-10** |
| SSp-ul_L plan vs CFA_R move | **0** |
| SSp-ul_R plan vs CFA_R move | **0** |
| SSp-un_L plan vs CFA_R move | **0** |
| SSp-un_R plan vs CFA_R move | **0** |
| VISa_L plan vs CFA_R move | **0** |
| VISam_L plan vs CFA_R move | **0** |
| VISam_R plan vs CFA_R move | **0** |
| VISa_R plan vs CFA_R move | **0** |
| VISp_L plan vs CFA_R move | **0** |
| VISpm_L plan vs CFA_R move | **0** |
| VISpm_R plan vs CFA_R move | **0** |
| VISp_R plan vs CFA_R move | **0** |
| VISrl_L plan vs CFA_R move | **0** |
| VISrl_R plan vs CFA_R move | **0** |
| CFA_L rest vs CFA_R move | **0** |
| CFA_R rest vs CFA_R move | **0** |
| MOp_L rest vs CFA_R move | **0** |
| MOp_R rest vs CFA_R move | **0** |
| MOs_L rest vs CFA_R move | **0** |
| MOs_R rest vs CFA_R move | **0** |
| RFA_L rest vs CFA_R move | **0** |

| | |
|---|---|
| RFA_R rest vs CFA_R move | **0** |
| RSPagl_L rest vs CFA_R move | **0** |
| RSPagl_R rest vs CFA_R move | **0** |
| RSPd_L rest vs CFA_R move | **0** |
| RSPd_R rest vs CFA_R move | **0** |
| SSp-bfd_L rest vs CFA_R move | **0** |
| SSp-bfd_R rest vs CFA_R move | **0** |
| SSp-ll_L rest vs CFA_R move | **0** |
| SSp-ll_R rest vs CFA_R move | **0** |
| SSp-n_L rest vs CFA_R move | **0** |
| SSp-n_R rest vs CFA_R move | **0** |
| SSp-tr_L rest vs CFA_R move | **0** |
| SSp-tr_R rest vs CFA_R move | **0** |
| SSp-ul_L rest vs CFA_R move | **0** |
| SSp-ul_R rest vs CFA_R move | **0** |
| SSp-un_L rest vs CFA_R move | **0** |
| SSp-un_R rest vs CFA_R move | **0** |
| VISa_L rest vs CFA_R move | **0** |
| VISam_L rest vs CFA_R move | **0** |
| VISam_R rest vs CFA_R move | **0** |
| VISa_R rest vs CFA_R move | **0** |
| VISp_L rest vs CFA_R move | **0** |
| VISpm_L rest vs CFA_R move | **0** |
| VISpm_R rest vs CFA_R move | **0** |
| VISp_R rest vs CFA_R move | **0** |
| VISrl_L rest vs CFA_R move | **0** |
| VISrl_R rest vs CFA_R move | **0** |
| MOp_R move vs MOp_L move | 1 |
| MOs_L move vs MOp_L move | 0.014 |
| MOs_R move vs MOp_L move | 0.51 |
| RFA_L move vs MOp_L move | 0.023 |
| RFA_R move vs MOp_L move | 0.45 |
| RSPagl_L move vs MOp_L move | 1 |
| RSPagl_R move vs MOp_L move | 1 |
| RSPd_L move vs MOp_L move | 1 |
| RSPd_R move vs MOp_L move | 1 |
| SSp-bfd_L move vs MOp_L move | **7e-08** |
| SSp-bfd_R move vs MOp_L move | **0.0007** |
| SSp-ll_L move vs MOp_L move | 1 |
| SSp-ll_R move vs MOp_L move | 0.91 |
| SSp-n_L move vs MOp_L move | **0.0036** |
| SSp-n_R move vs MOp_L move | 0.011 |
| SSp-tr_L move vs MOp_L move | 1 |
| SSp-tr_R move vs MOp_L move | 1 |
| SSp-ul_L move vs MOp_L move | 1 |
| SSp-ul_R move vs MOp_L move | 1 |

| | |
|---|---|
| SSp-un_L move vs MOp_L move | 0.93 |
| SSp-un_R move vs MOp_L move | 1 |
| VISa_L move vs MOp_L move | 0.98 |
| VISam_L move vs MOp_L move | 0.1 |
| VISam_R move vs MOp_L move | 0.99 |
| VISa_R move vs MOp_L move | 1 |
| VISp_L move vs MOp_L move | **0** |
| VISpm_L move vs MOp_L move | **1.4e-06** |
| VISpm_R move vs MOp_L move | 0.036 |
| VISp_R move vs MOp_L move | **2.7e-06** |
| VISrl_L move vs MOp_L move | **0** |
| VISrl_R move vs MOp_L move | **1.4e-07** |
| CFA_L plan vs MOp_L move | **0** |
| CFA_R plan vs MOp_L move | **0** |
| MOp_L plan vs MOp_L move | **0** |
| MOp_R plan vs MOp_L move | **0** |
| MOs_L plan vs MOp_L move | **6.6e-06** |
| MOs_R plan vs MOp_L move | **1e-07** |
| RFA_L plan vs MOp_L move | **2.2e-06** |
| RFA_R plan vs MOp_L move | **1.6e-08** |
| RSPagl_L plan vs MOp_L move | **0** |
| RSPagl_R plan vs MOp_L move | **0** |
| RSPd_L plan vs MOp_L move | **0** |
| RSPd_R plan vs MOp_L move | **0** |
| SSp-bfd_L plan vs MOp_L move | **0** |
| SSp-bfd_R plan vs MOp_L move | **0** |
| SSp-ll_L plan vs MOp_L move | **6.9e-08** |
| SSp-ll_R plan vs MOp_L move | **1.9e-06** |
| SSp-n_L plan vs MOp_L move | **0** |
| SSp-n_R plan vs MOp_L move | **0** |
| SSp-tr_L plan vs MOp_L move | **0** |
| SSp-tr_R plan vs MOp_L move | **3.3e-08** |
| SSp-ul_L plan vs MOp_L move | **0** |
| SSp-ul_R plan vs MOp_L move | **0** |
| SSp-un_L plan vs MOp_L move | **0** |
| SSp-un_R plan vs MOp_L move | **0** |
| VISa_L plan vs MOp_L move | **0** |
| VISam_L plan vs MOp_L move | **0** |
| VISam_R plan vs MOp_L move | **0** |
| VISa_R plan vs MOp_L move | **0** |
| VISp_L plan vs MOp_L move | **0** |
| VISpm_L plan vs MOp_L move | **0** |
| VISpm_R plan vs MOp_L move | **0** |
| VISp_R plan vs MOp_L move | **0** |
| VISrl_L plan vs MOp_L move | **0** |
| VISrl_R plan vs MOp_L move | **0** |

| | |
|---|---|
| CFA_L rest vs MOp_L move | **0** |
| CFA_R rest vs MOp_L move | **0** |
| MOp_L rest vs MOp_L move | **0** |
| MOp_R rest vs MOp_L move | **0** |
| MOs_L rest vs MOp_L move | **0** |
| MOs_R rest vs MOp_L move | **0** |
| RFA_L rest vs MOp_L move | **0** |
| RFA_R rest vs MOp_L move | **0** |
| RSPagl_L rest vs MOp_L move | **0** |
| RSPagl_R rest vs MOp_L move | **0** |
| RSPd_L rest vs MOp_L move | **0** |
| RSPd_R rest vs MOp_L move | **0** |
| SSp-bfd_L rest vs MOp_L move | **0** |
| SSp-bfd_R rest vs MOp_L move | **0** |
| SSp-ll_L rest vs MOp_L move | **0** |
| SSp-ll_R rest vs MOp_L move | **0** |
| SSp-n_L rest vs MOp_L move | **0** |
| SSp-n_R rest vs MOp_L move | **0** |
| SSp-tr_L rest vs MOp_L move | **0** |
| SSp-tr_R rest vs MOp_L move | **0** |
| SSp-ul_L rest vs MOp_L move | **0** |
| SSp-ul_R rest vs MOp_L move | **0** |
| SSp-un_L rest vs MOp_L move | **0** |
| SSp-un_R rest vs MOp_L move | **0** |
| VISa_L rest vs MOp_L move | **0** |
| VISam_L rest vs MOp_L move | **0** |
| VISam_R rest vs MOp_L move | **0** |
| VISa_R rest vs MOp_L move | **0** |
| VISp_L rest vs MOp_L move | **0** |
| VISpm_L rest vs MOp_L move | **0** |
| VISpm_R rest vs MOp_L move | **0** |
| VISp_R rest vs MOp_L move | **0** |
| VISrl_L rest vs MOp_L move | **0** |
| VISrl_R rest vs MOp_L move | **0** |
| MOs_L move vs MOp_R move | **0.0057** |
| MOs_R move vs MOp_R move | 0.33 |
| RFA_L move vs MOp_R move | **0.0099** |
| RFA_R move vs MOp_R move | 0.28 |
| RSPagl_L move vs MOp_R move | 1 |
| RSPagl_R move vs MOp_R move | 1 |
| RSPd_L move vs MOp_R move | 1 |
| RSPd_R move vs MOp_R move | 1 |
| SSp-bfd_L move vs MOp_R move | **2.7e-07** |
| SSp-bfd_R move vs MOp_R move | **0.0019** |
| SSp-ll_L move vs MOp_R move | 1 |
| SSp-ll_R move vs MOp_R move | 0.78 |

| | |
|---|---|
| SSp-n_L move vs MOp_R move | **0.009** |
| SSp-n_R move vs MOp_R move | 0.026 |
| SSp-tr_L move vs MOp_R move | 1 |
| SSp-tr_R move vs MOp_R move | 1 |
| SSp-ul_L move vs MOp_R move | 1 |
| SSp-ul_R move vs MOp_R move | 1 |
| SSp-un_L move vs MOp_R move | 0.98 |
| SSp-un_R move vs MOp_R move | 1 |
| VISa_L move vs MOp_R move | 1 |
| VISam_L move vs MOp_R move | 0.2 |
| VISam_R move vs MOp_R move | 1 |
| VISa_R move vs MOp_R move | 1 |
| VISp_L move vs MOp_R move | **0** |
| VISpm_L move vs MOp_R move | **5e-06** |
| VISpm_R move vs MOp_R move | 0.077 |
| VISp_R move vs MOp_R move | **9.4e-06** |
| VISrl_L move vs MOp_R move | **0** |
| VISrl_R move vs MOp_R move | **5.1e-07** |
| CFA_L plan vs MOp_R move | **0** |
| CFA_R plan vs MOp_R move | **0** |
| MOp_L plan vs MOp_R move | **0** |
| MOp_R plan vs MOp_R move | **0** |
| MOs_L plan vs MOp_R move | **2.2e-05** |
| MOs_R plan vs MOp_R move | **3.9e-07** |
| RFA_L plan vs MOp_R move | **7.5e-06** |
| RFA_R plan vs MOp_R move | **6.6e-08** |
| RSPagl_L plan vs MOp_R move | **0** |
| RSPagl_R plan vs MOp_R move | **0** |
| RSPd_L plan vs MOp_R move | **0** |
| RSPd_R plan vs MOp_R move | **0** |
| SSp-bfd_L plan vs MOp_R move | **0** |
| SSp-bfd_R plan vs MOp_R move | **0** |
| SSp-ll_L plan vs MOp_R move | **2.7e-07** |
| SSp-ll_R plan vs MOp_R move | **6.4e-06** |
| SSp-n_L plan vs MOp_R move | **0** |
| SSp-n_R plan vs MOp_R move | **0** |
| SSp-tr_L plan vs MOp_R move | **6.2e-10** |
| SSp-tr_R plan vs MOp_R move | **1.3e-07** |
| SSp-ul_L plan vs MOp_R move | **0** |
| SSp-ul_R plan vs MOp_R move | **0** |
| SSp-un_L plan vs MOp_R move | **0** |
| SSp-un_R plan vs MOp_R move | **0** |
| VISa_L plan vs MOp_R move | **0** |
| VISam_L plan vs MOp_R move | **0** |
| VISam_R plan vs MOp_R move | **0** |
| VISa_R plan vs MOp_R move | **0** |

| | |
|---|---|
| VISp_L plan vs MOp_R move | **0** |
| VISpm_L plan vs MOp_R move | **0** |
| VISpm_R plan vs MOp_R move | **0** |
| VISp_R plan vs MOp_R move | **0** |
| VISrl_L plan vs MOp_R move | **0** |
| VISrl_R plan vs MOp_R move | **0** |
| CFA_L rest vs MOp_R move | **0** |
| CFA_R rest vs MOp_R move | **0** |
| MOp_L rest vs MOp_R move | **0** |
| MOp_R rest vs MOp_R move | **0** |
| MOs_L rest vs MOp_R move | **0** |
| MOs_R rest vs MOp_R move | **0** |
| RFA_L rest vs MOp_R move | **0** |
| RFA_R rest vs MOp_R move | **0** |
| RSPagl_L rest vs MOp_R move | **0** |
| RSPagl_R rest vs MOp_R move | **0** |
| RSPd_L rest vs MOp_R move | **0** |
| RSPd_R rest vs MOp_R move | **0** |
| SSp-bfd_L rest vs MOp_R move | **0** |
| SSp-bfd_R rest vs MOp_R move | **0** |
| SSp-ll_L rest vs MOp_R move | **0** |
| SSp-ll_R rest vs MOp_R move | **0** |
| SSp-n_L rest vs MOp_R move | **0** |
| SSp-n_R rest vs MOp_R move | **0** |
| SSp-tr_L rest vs MOp_R move | **0** |
| SSp-tr_R rest vs MOp_R move | **0** |
| SSp-ul_L rest vs MOp_R move | **0** |
| SSp-ul_R rest vs MOp_R move | **0** |
| SSp-un_L rest vs MOp_R move | **0** |
| SSp-un_R rest vs MOp_R move | **0** |
| VISa_L rest vs MOp_R move | **0** |
| VISam_L rest vs MOp_R move | **0** |
| VISam_R rest vs MOp_R move | **0** |
| VISa_R rest vs MOp_R move | **0** |
| VISp_L rest vs MOp_R move | **0** |
| VISpm_L rest vs MOp_R move | **0** |
| VISpm_R rest vs MOp_R move | **0** |
| VISp_R rest vs MOp_R move | **0** |
| VISrl_L rest vs MOp_R move | **0** |
| VISrl_R rest vs MOp_R move | **0** |
| MOs_R move vs MOs_L move | 1 |
| RFA_L move vs MOs_L move | 1 |
| RFA_R move vs MOs_L move | 1 |
| RSPagl_L move vs MOs_L move | **5.9e-09** |
| RSPagl_R move vs MOs_L move | **3.3e-06** |
| RSPd_L move vs MOs_L move | 0.016 |

| | |
|---|---|
| RSPd_R move vs MOs_L move | **0.0064** |
| SSp-bfd_L move vs MOs_L move | **0** |
| SSp-bfd_R move vs MOs_L move | **0** |
| SSp-ll_L move vs MOs_L move | 1 |
| SSp-ll_R move vs MOs_L move | 1 |
| SSp-n_L move vs MOs_L move | **0** |
| SSp-n_R move vs MOs_L move | **0** |
| SSp-tr_L move vs MOs_L move | 0.081 |
| SSp-tr_R move vs MOs_L move | 0.85 |
| SSp-ul_L move vs MOs_L move | **0.003** |
| SSp-ul_R move vs MOs_L move | 0.24 |
| SSp-un_L move vs MOs_L move | **0** |
| SSp-un_R move vs MOs_L move | **1.4e-05** |
| VISa_L move vs MOs_L move | **0** |
| VISam_L move vs MOs_L move | **0** |
| VISam_R move vs MOs_L move | **1.7e-10** |
| VISa_R move vs MOs_L move | **2.5e-06** |
| VISp_L move vs MOs_L move | **0** |
| VISpm_L move vs MOs_L move | **0** |
| VISpm_R move vs MOs_L move | **0** |
| VISp_R move vs MOs_L move | **0** |
| VISrl_L move vs MOs_L move | **0** |
| VISrl_R move vs MOs_L move | **0** |
| CFA_L plan vs MOs_L move | **0** |
| CFA_R plan vs MOs_L move | **0** |
| MOp_L plan vs MOs_L move | **0** |
| MOp_R plan vs MOs_L move | **0** |
| MOs_L plan vs MOs_L move | **0** |
| MOs_R plan vs MOs_L move | **0** |
| RFA_L plan vs MOs_L move | **0** |
| RFA_R plan vs MOs_L move | **0** |
| RSPagl_L plan vs MOs_L move | **0** |
| RSPagl_R plan vs MOs_L move | **0** |
| RSPd_L plan vs MOs_L move | **0** |
| RSPd_R plan vs MOs_L move | **0** |
| SSp-bfd_L plan vs MOs_L move | **0** |
| SSp-bfd_R plan vs MOs_L move | **0** |
| SSp-ll_L plan vs MOs_L move | **0** |
| SSp-ll_R plan vs MOs_L move | **0** |
| SSp-n_L plan vs MOs_L move | **0** |
| SSp-n_R plan vs MOs_L move | **0** |
| SSp-tr_L plan vs MOs_L move | **0** |
| SSp-tr_R plan vs MOs_L move | **0** |
| SSp-ul_L plan vs MOs_L move | **0** |
| SSp-ul_R plan vs MOs_L move | **0** |
| SSp-un_L plan vs MOs_L move | **0** |

| | |
|---|---|
| SSp-un_R plan vs MOs_L move | **0** |
| VISa_L plan vs MOs_L move | **0** |
| VISam_L plan vs MOs_L move | **0** |
| VISam_R plan vs MOs_L move | **0** |
| VISa_R plan vs MOs_L move | **0** |
| VISp_L plan vs MOs_L move | **0** |
| VISpm_L plan vs MOs_L move | **0** |
| VISpm_R plan vs MOs_L move | **0** |
| VISp_R plan vs MOs_L move | **0** |
| VISrl_L plan vs MOs_L move | **0** |
| VISrl_R plan vs MOs_L move | **0** |
| CFA_L rest vs MOs_L move | **0** |
| CFA_R rest vs MOs_L move | **0** |
| MOp_L rest vs MOs_L move | **0** |
| MOp_R rest vs MOs_L move | **0** |
| MOs_L rest vs MOs_L move | **0** |
| MOs_R rest vs MOs_L move | **0** |
| RFA_L rest vs MOs_L move | **0** |
| RFA_R rest vs MOs_L move | **0** |
| RSPagl_L rest vs MOs_L move | **0** |
| RSPagl_R rest vs MOs_L move | **0** |
| RSPd_L rest vs MOs_L move | **0** |
| RSPd_R rest vs MOs_L move | **0** |
| SSp-bfd_L rest vs MOs_L move | **0** |
| SSp-bfd_R rest vs MOs_L move | **0** |
| SSp-ll_L rest vs MOs_L move | **0** |
| SSp-ll_R rest vs MOs_L move | **0** |
| SSp-n_L rest vs MOs_L move | **0** |
| SSp-n_R rest vs MOs_L move | **0** |
| SSp-tr_L rest vs MOs_L move | **0** |
| SSp-tr_R rest vs MOs_L move | **0** |
| SSp-ul_L rest vs MOs_L move | **0** |
| SSp-ul_R rest vs MOs_L move | **0** |
| SSp-un_L rest vs MOs_L move | **0** |
| SSp-un_R rest vs MOs_L move | **0** |
| VISa_L rest vs MOs_L move | **0** |
| VISam_L rest vs MOs_L move | **0** |
| VISam_R rest vs MOs_L move | **0** |
| VISa_R rest vs MOs_L move | **0** |
| VISp_L rest vs MOs_L move | **0** |
| VISpm_L rest vs MOs_L move | **0** |
| VISpm_R rest vs MOs_L move | **0** |
| VISp_R rest vs MOs_L move | **0** |
| VISrl_L rest vs MOs_L move | **0** |
| VISrl_R rest vs MOs_L move | **0** |
| RFA_L move vs MOs_R move | 1 |

| | |
|---|---|
| RFA_R move vs MOs_R move | 1 |
| RSPagl_L move vs MOs_R move | **1.1e-05** |
| RSPagl_R move vs MOs_R move | **0.0018** |
| RSPd_L move vs MOs_R move | 0.54 |
| RSPd_R move vs MOs_R move | 0.35 |
| SSp-bfd_L move vs MOs_R move | **0** |
| SSp-bfd_R move vs MOs_R move | **0** |
| SSp-ll_L move vs MOs_R move | 1 |
| SSp-ll_R move vs MOs_R move | 1 |
| SSp-n_L move vs MOs_R move | **0** |
| SSp-n_R move vs MOs_R move | **0** |
| SSp-tr_L move vs MOs_R move | 0.89 |
| SSp-tr_R move vs MOs_R move | 1 |
| SSp-ul_L move vs MOs_R move | 0.24 |
| SSp-ul_R move vs MOs_R move | 0.99 |
| SSp-un_L move vs MOs_R move | **5e-07** |
| SSp-un_R move vs MOs_R move | **0.0057** |
| VISa_L move vs MOs_R move | **1.5e-06** |
| VISam_L move vs MOs_R move | **0** |
| VISam_R move vs MOs_R move | **2.6e-06** |
| VISa_R move vs MOs_R move | **0.0015** |
| VISp_L move vs MOs_R move | **0** |
| VISpm_L move vs MOs_R move | **0** |
| VISpm_R move vs MOs_R move | **0** |
| VISp_R move vs MOs_R move | **0** |
| VISrl_L move vs MOs_R move | **0** |
| VISrl_R move vs MOs_R move | **0** |
| CFA_L plan vs MOs_R move | **0** |
| CFA_R plan vs MOs_R move | **0** |
| MOp_L plan vs MOs_R move | **0** |
| MOp_R plan vs MOs_R move | **0** |
| MOs_L plan vs MOs_R move | **0** |
| MOs_R plan vs MOs_R move | **0** |
| RFA_L plan vs MOs_R move | **0** |
| RFA_R plan vs MOs_R move | **0** |
| RSPagl_L plan vs MOs_R move | **0** |
| RSPagl_R plan vs MOs_R move | **0** |
| RSPd_L plan vs MOs_R move | **0** |
| RSPd_R plan vs MOs_R move | **0** |
| SSp-bfd_L plan vs MOs_R move | **0** |
| SSp-bfd_R plan vs MOs_R move | **0** |
| SSp-ll_L plan vs MOs_R move | **0** |
| SSp-ll_R plan vs MOs_R move | **0** |
| SSp-n_L plan vs MOs_R move | **0** |
| SSp-n_R plan vs MOs_R move | **0** |
| SSp-tr_L plan vs MOs_R move | **0** |

| | |
|---|---|
| SSp-tr_R plan vs MOs_R move | **0** |
| SSp-ul_L plan vs MOs_R move | **0** |
| SSp-ul_R plan vs MOs_R move | **0** |
| SSp-un_L plan vs MOs_R move | **0** |
| SSp-un_R plan vs MOs_R move | **0** |
| VISa_L plan vs MOs_R move | **0** |
| VISam_L plan vs MOs_R move | **0** |
| VISam_R plan vs MOs_R move | **0** |
| VISa_R plan vs MOs_R move | **0** |
| VISp_L plan vs MOs_R move | **0** |
| VISpm_L plan vs MOs_R move | **0** |
| VISpm_R plan vs MOs_R move | **0** |
| VISp_R plan vs MOs_R move | **0** |
| VISrl_L plan vs MOs_R move | **0** |
| VISrl_R plan vs MOs_R move | **0** |
| CFA_L rest vs MOs_R move | **0** |
| CFA_R rest vs MOs_R move | **0** |
| MOp_L rest vs MOs_R move | **0** |
| MOp_R rest vs MOs_R move | **0** |
| MOs_L rest vs MOs_R move | **0** |
| MOs_R rest vs MOs_R move | **0** |
| RFA_L rest vs MOs_R move | **0** |
| RFA_R rest vs MOs_R move | **0** |
| RSPagl_L rest vs MOs_R move | **0** |
| RSPagl_R rest vs MOs_R move | **0** |
| RSPd_L rest vs MOs_R move | **0** |
| RSPd_R rest vs MOs_R move | **0** |
| SSp-bfd_L rest vs MOs_R move | **0** |
| SSp-bfd_R rest vs MOs_R move | **0** |
| SSp-ll_L rest vs MOs_R move | **0** |
| SSp-ll_R rest vs MOs_R move | **0** |
| SSp-n_L rest vs MOs_R move | **0** |
| SSp-n_R rest vs MOs_R move | **0** |
| SSp-tr_L rest vs MOs_R move | **0** |
| SSp-tr_R rest vs MOs_R move | **0** |
| SSp-ul_L rest vs MOs_R move | **0** |
| SSp-ul_R rest vs MOs_R move | **0** |
| SSp-un_L rest vs MOs_R move | **0** |
| SSp-un_R rest vs MOs_R move | **0** |
| VISa_L rest vs MOs_R move | **0** |
| VISam_L rest vs MOs_R move | **0** |
| VISam_R rest vs MOs_R move | **0** |
| VISa_R rest vs MOs_R move | **0** |
| VISp_L rest vs MOs_R move | **0** |
| VISpm_L rest vs MOs_R move | **0** |
| VISpm_R rest vs MOs_R move | **0** |

| | |
|---|---|
| VISp_R rest vs MOs_R move | **0** |
| VISrl_L rest vs MOs_R move | **0** |
| VISrl_R rest vs MOs_R move | **0** |
| RFA_R move vs RFA_L move | 1 |
| RSPagl_L move vs RFA_L move | **1.6e-08** |
| RSPagl_R move vs RFA_L move | **7.1e-06** |
| RSPd_L move vs RFA_L move | 0.026 |
| RSPd_R move vs RFA_L move | 0.011 |
| SSp-bfd_L move vs RFA_L move | **0** |
| SSp-bfd_R move vs RFA_L move | **0** |
| SSp-ll_L move vs RFA_L move | 1 |
| SSp-ll_R move vs RFA_L move | 1 |
| SSp-n_L move vs RFA_L move | **0** |
| SSp-n_R move vs RFA_L move | **0** |
| SSp-tr_L move vs RFA_L move | 0.12 |
| SSp-tr_R move vs RFA_L move | 0.92 |
| SSp-ul_L move vs RFA_L move | **0.0054** |
| SSp-ul_R move vs RFA_L move | 0.33 |
| SSp-un_L move vs RFA_L move | **0** |
| SSp-un_R move vs RFA_L move | **3e-05** |
| VISa_L move vs RFA_L move | **6e-10** |
| VISam_L move vs RFA_L move | **0** |
| VISam_R move vs RFA_L move | **1.9e-09** |
| VISa_R move vs RFA_L move | **5.5e-06** |
| VISp_L move vs RFA_L move | **0** |
| VISpm_L move vs RFA_L move | **0** |
| VISpm_R move vs RFA_L move | **0** |
| VISp_R move vs RFA_L move | **0** |
| VISrl_L move vs RFA_L move | **0** |
| VISrl_R move vs RFA_L move | **0** |
| CFA_L plan vs RFA_L move | **0** |
| CFA_R plan vs RFA_L move | **0** |
| MOp_L plan vs RFA_L move | **0** |
| MOp_R plan vs RFA_L move | **0** |
| MOs_L plan vs RFA_L move | **0** |
| MOs_R plan vs RFA_L move | **0** |
| RFA_L plan vs RFA_L move | **0** |
| RFA_R plan vs RFA_L move | **0** |
| RSPagl_L plan vs RFA_L move | **0** |
| RSPagl_R plan vs RFA_L move | **0** |
| RSPd_L plan vs RFA_L move | **0** |
| RSPd_R plan vs RFA_L move | **0** |
| SSp-bfd_L plan vs RFA_L move | **0** |
| SSp-bfd_R plan vs RFA_L move | **0** |
| SSp-ll_L plan vs RFA_L move | **0** |
| SSp-ll_R plan vs RFA_L move | **0** |

| | |
|---|---|
| SSp-n_L plan vs RFA_L move | **0** |
| SSp-n_R plan vs RFA_L move | **0** |
| SSp-tr_L plan vs RFA_L move | **0** |
| SSp-tr_R plan vs RFA_L move | **0** |
| SSp-ul_L plan vs RFA_L move | **0** |
| SSp-ul_R plan vs RFA_L move | **0** |
| SSp-un_L plan vs RFA_L move | **0** |
| SSp-un_R plan vs RFA_L move | **0** |
| VISa_L plan vs RFA_L move | **0** |
| VISam_L plan vs RFA_L move | **0** |
| VISam_R plan vs RFA_L move | **0** |
| VISa_R plan vs RFA_L move | **0** |
| VISp_L plan vs RFA_L move | **0** |
| VISpm_L plan vs RFA_L move | **0** |
| VISpm_R plan vs RFA_L move | **0** |
| VISp_R plan vs RFA_L move | **0** |
| VISrl_L plan vs RFA_L move | **0** |
| VISrl_R plan vs RFA_L move | **0** |
| CFA_L rest vs RFA_L move | **0** |
| CFA_R rest vs RFA_L move | **0** |
| MOp_L rest vs RFA_L move | **0** |
| MOp_R rest vs RFA_L move | **0** |
| MOs_L rest vs RFA_L move | **0** |
| MOs_R rest vs RFA_L move | **0** |
| RFA_L rest vs RFA_L move | **0** |
| RFA_R rest vs RFA_L move | **0** |
| RSPagl_L rest vs RFA_L move | **0** |
| RSPagl_R rest vs RFA_L move | **0** |
| RSPd_L rest vs RFA_L move | **0** |
| RSPd_R rest vs RFA_L move | **0** |
| SSp-bfd_L rest vs RFA_L move | **0** |
| SSp-bfd_R rest vs RFA_L move | **0** |
| SSp-ll_L rest vs RFA_L move | **0** |
| SSp-ll_R rest vs RFA_L move | **0** |
| SSp-n_L rest vs RFA_L move | **0** |
| SSp-n_R rest vs RFA_L move | **0** |
| SSp-tr_L rest vs RFA_L move | **0** |
| SSp-tr_R rest vs RFA_L move | **0** |
| SSp-ul_L rest vs RFA_L move | **0** |
| SSp-ul_R rest vs RFA_L move | **0** |
| SSp-un_L rest vs RFA_L move | **0** |
| SSp-un_R rest vs RFA_L move | **0** |
| VISa_L rest vs RFA_L move | **0** |
| VISam_L rest vs RFA_L move | **0** |
| VISam_R rest vs RFA_L move | **0** |
| VISa_R rest vs RFA_L move | **0** |

| | |
|---|---|
| VISp_L rest vs RFA_L move | **0** |
| VISpm_L rest vs RFA_L move | **0** |
| VISpm_R rest vs RFA_L move | **0** |
| VISp_R rest vs RFA_L move | **0** |
| VISrl_L rest vs RFA_L move | **0** |
| VISrl_R rest vs RFA_L move | **0** |
| RSPagl_L move vs RFA_R move | **7.5e-06** |
| RSPagl_R move vs RFA_R move | **0.0013** |
| RSPd_L move vs RFA_R move | 0.47 |
| RSPd_R move vs RFA_R move | 0.3 |
| SSp-bfd_L move vs RFA_R move | **0** |
| SSp-bfd_R move vs RFA_R move | **0** |
| SSp-ll_L move vs RFA_R move | 1 |
| SSp-ll_R move vs RFA_R move | 1 |
| SSp-n_L move vs RFA_R move | **0** |
| SSp-n_R move vs RFA_R move | **0** |
| SSp-tr_L move vs RFA_R move | 0.84 |
| SSp-tr_R move vs RFA_R move | 1 |
| SSp-ul_L move vs RFA_R move | 0.19 |
| SSp-ul_R move vs RFA_R move | 0.98 |
| SSp-un_L move vs RFA_R move | **3.2e-07** |
| SSp-un_R move vs RFA_R move | **0.0041** |
| VISa_L move vs RFA_R move | **9.9e-07** |
| VISam_L move vs RFA_R move | **0** |
| VISam_R move vs RFA_R move | **1.7e-06** |
| VISa_R move vs RFA_R move | **0.001** |
| VISp_L move vs RFA_R move | **0** |
| VISpm_L move vs RFA_R move | **0** |
| VISpm_R move vs RFA_R move | **0** |
| VISp_R move vs RFA_R move | **0** |
| VISrl_L move vs RFA_R move | **0** |
| VISrl_R move vs RFA_R move | **0** |
| CFA_L plan vs RFA_R move | **0** |
| CFA_R plan vs RFA_R move | **0** |
| MOp_L plan vs RFA_R move | **0** |
| MOp_R plan vs RFA_R move | **0** |
| MOs_L plan vs RFA_R move | **0** |
| MOs_R plan vs RFA_R move | **0** |
| RFA_L plan vs RFA_R move | **0** |
| RFA_R plan vs RFA_R move | **0** |
| RSPagl_L plan vs RFA_R move | **0** |
| RSPagl_R plan vs RFA_R move | **0** |
| RSPd_L plan vs RFA_R move | **0** |
| RSPd_R plan vs RFA_R move | **0** |
| SSp-bfd_L plan vs RFA_R move | **0** |
| SSp-bfd_R plan vs RFA_R move | **0** |

| | |
|---|---|
| SSp-ll_L plan vs RFA_R move | **0** |
| SSp-ll_R plan vs RFA_R move | **0** |
| SSp-n_L plan vs RFA_R move | **0** |
| SSp-n_R plan vs RFA_R move | **0** |
| SSp-tr_L plan vs RFA_R move | **0** |
| SSp-tr_R plan vs RFA_R move | **0** |
| SSp-ul_L plan vs RFA_R move | **0** |
| SSp-ul_R plan vs RFA_R move | **0** |
| SSp-un_L plan vs RFA_R move | **0** |
| SSp-un_R plan vs RFA_R move | **0** |
| VISa_L plan vs RFA_R move | **0** |
| VISam_L plan vs RFA_R move | **0** |
| VISam_R plan vs RFA_R move | **0** |
| VISa_R plan vs RFA_R move | **0** |
| VISp_L plan vs RFA_R move | **0** |
| VISpm_L plan vs RFA_R move | **0** |
| VISpm_R plan vs RFA_R move | **0** |
| VISp_R plan vs RFA_R move | **0** |
| VISrl_L plan vs RFA_R move | **0** |
| VISrl_R plan vs RFA_R move | **0** |
| CFA_L rest vs RFA_R move | **0** |
| CFA_R rest vs RFA_R move | **0** |
| MOp_L rest vs RFA_R move | **0** |
| MOp_R rest vs RFA_R move | **0** |
| MOs_L rest vs RFA_R move | **0** |
| MOs_R rest vs RFA_R move | **0** |
| RFA_L rest vs RFA_R move | **0** |
| RFA_R rest vs RFA_R move | **0** |
| RSPagl_L rest vs RFA_R move | **0** |
| RSPagl_R rest vs RFA_R move | **0** |
| RSPd_L rest vs RFA_R move | **0** |
| RSPd_R rest vs RFA_R move | **0** |
| SSp-bfd_L rest vs RFA_R move | **0** |
| SSp-bfd_R rest vs RFA_R move | **0** |
| SSp-ll_L rest vs RFA_R move | **0** |
| SSp-ll_R rest vs RFA_R move | **0** |
| SSp-n_L rest vs RFA_R move | **0** |
| SSp-n_R rest vs RFA_R move | **0** |
| SSp-tr_L rest vs RFA_R move | **0** |
| SSp-tr_R rest vs RFA_R move | **0** |
| SSp-ul_L rest vs RFA_R move | **0** |
| SSp-ul_R rest vs RFA_R move | **0** |
| SSp-un_L rest vs RFA_R move | **0** |
| SSp-un_R rest vs RFA_R move | **0** |
| VISa_L rest vs RFA_R move | **0** |
| VISam_L rest vs RFA_R move | **0** |

| | |
|---|---|
| VISam_R rest vs RFA_R move | **0** |
| VISa_R rest vs RFA_R move | **0** |
| VISp_L rest vs RFA_R move | **0** |
| VISpm_L rest vs RFA_R move | **0** |
| VISpm_R rest vs RFA_R move | **0** |
| VISp_R rest vs RFA_R move | **0** |
| VISrl_L rest vs RFA_R move | **0** |
| VISrl_R rest vs RFA_R move | **0** |
| RSPagl_R move vs RSPagl_L move | 1 |
| RSPd_L move vs RSPagl_L move | 1 |
| RSPd_R move vs RSPagl_L move | 1 |
| SSp-bfd_L move vs RSPagl_L move | 0.053 |
| SSp-bfd_R move vs RSPagl_L move | 0.98 |
| SSp-ll_L move vs RSPagl_L move | 0.017 |
| SSp-ll_R move vs RSPagl_L move | **0.00019** |
| SSp-n_L move vs RSPagl_L move | 1 |
| SSp-n_R move vs RSPagl_L move | 1 |
| SSp-tr_L move vs RSPagl_L move | 0.95 |
| SSp-tr_R move vs RSPagl_L move | 0.16 |
| SSp-ul_L move vs RSPagl_L move | 1 |
| SSp-ul_R move vs RSPagl_L move | 0.74 |
| SSp-un_L move vs RSPagl_L move | 1 |
| SSp-un_R move vs RSPagl_L move | 1 |
| VISa_L move vs RSPagl_L move | 1 |
| VISam_L move vs RSPagl_L move | 1 |
| VISam_R move vs RSPagl_L move | 1 |
| VISa_R move vs RSPagl_L move | 1 |
| VISp_L move vs RSPagl_L move | **0.001** |
| VISpm_L move vs RSPagl_L move | 0.23 |
| VISpm_R move vs RSPagl_L move | 1 |
| VISp_R move vs RSPagl_L move | 0.31 |
| VISrl_L move vs RSPagl_L move | **0.00053** |
| VISrl_R move vs RSPagl_L move | 0.075 |
| CFA_L plan vs RSPagl_L move | **0.0012** |
| CFA_R plan vs RSPagl_L move | **0.00025** |
| MOp_L plan vs RSPagl_L move | **0.00011** |
| MOp_R plan vs RSPagl_L move | **5.7e-05** |
| MOs_L plan vs RSPagl_L move | 0.43 |
| MOs_R plan vs RSPagl_L move | 0.065 |
| RFA_L plan vs RSPagl_L move | 0.28 |
| RFA_R plan vs RSPagl_L move | 0.023 |
| RSPagl_L plan vs RSPagl_L move | **5.5e-07** |
| RSPagl_R plan vs RSPagl_L move | **3.3e-06** |
| RSPd_L plan vs RSPagl_L move | **0.00071** |
| RSPd_R plan vs RSPagl_L move | **0.00038** |
| SSp-bfd_L plan vs RSPagl_L move | **0** |

| | |
|---|---|
| SSp-bfd_R plan vs RSPagl_L move | **0** |
| SSp-ll_L plan vs RSPagl_L move | 0.053 |
| SSp-ll_R plan vs RSPagl_L move | 0.26 |
| SSp-n_L plan vs RSPagl_L move | **0** |
| SSp-n_R plan vs RSPagl_L move | **0** |
| SSp-tr_L plan vs RSPagl_L move | **0.0022** |
| SSp-tr_R plan vs RSPagl_L move | 0.035 |
| SSp-ul_L plan vs RSPagl_L move | **6.4e-05** |
| SSp-ul_R plan vs RSPagl_L move | **0.00041** |
| SSp-un_L plan vs RSPagl_L move | **1.2e-07** |
| SSp-un_R plan vs RSPagl_L move | **2.3e-06** |
| VISa_L plan vs RSPagl_L move | **1.1e-06** |
| VISam_L plan vs RSPagl_L move | **2e-08** |
| VISam_R plan vs RSPagl_L move | **2.4e-07** |
| VISa_R plan vs RSPagl_L move | **1e-05** |
| VISp_L plan vs RSPagl_L move | **0** |
| VISpm_L plan vs RSPagl_L move | **0** |
| VISpm_R plan vs RSPagl_L move | **0** |
| VISp_R plan vs RSPagl_L move | **0** |
| VISrl_L plan vs RSPagl_L move | **0** |
| VISrl_R plan vs RSPagl_L move | **0** |
| CFA_L rest vs RSPagl_L move | **0** |
| CFA_R rest vs RSPagl_L move | **0** |
| MOp_L rest vs RSPagl_L move | **0** |
| MOp_R rest vs RSPagl_L move | **0** |
| MOs_L rest vs RSPagl_L move | **0** |
| MOs_R rest vs RSPagl_L move | **0** |
| RFA_L rest vs RSPagl_L move | **0** |
| RFA_R rest vs RSPagl_L move | **0** |
| RSPagl_L rest vs RSPagl_L move | **0** |
| RSPagl_R rest vs RSPagl_L move | **0** |
| RSPd_L rest vs RSPagl_L move | **0** |
| RSPd_R rest vs RSPagl_L move | **0** |
| SSp-bfd_L rest vs RSPagl_L move | **0** |
| SSp-bfd_R rest vs RSPagl_L move | **0** |
| SSp-ll_L rest vs RSPagl_L move | **0** |
| SSp-ll_R rest vs RSPagl_L move | **0** |
| SSp-n_L rest vs RSPagl_L move | **0** |
| SSp-n_R rest vs RSPagl_L move | **0** |
| SSp-tr_L rest vs RSPagl_L move | **0** |
| SSp-tr_R rest vs RSPagl_L move | **0** |
| SSp-ul_L rest vs RSPagl_L move | **0** |
| SSp-ul_R rest vs RSPagl_L move | **0** |
| SSp-un_L rest vs RSPagl_L move | **0** |
| SSp-un_R rest vs RSPagl_L move | **0** |
| VISa_L rest vs RSPagl_L move | **0** |

| | |
|---|---|
| VISam_L rest vs RSPagl_L move | **0** |
| VISam_R rest vs RSPagl_L move | **0** |
| VISa_R rest vs RSPagl_L move | **0** |
| VISp_L rest vs RSPagl_L move | **0** |
| VISpm_L rest vs RSPagl_L move | **0** |
| VISpm_R rest vs RSPagl_L move | **0** |
| VISp_R rest vs RSPagl_L move | **0** |
| VISrl_L rest vs RSPagl_L move | **0** |
| VISrl_R rest vs RSPagl_L move | **0** |
| RSPd_L move vs RSPagl_R move | 1 |
| RSPd_R move vs RSPagl_R move | 1 |
| SSp-bfd_L move vs RSPagl_R move | **0.00085** |
| SSp-bfd_R move vs RSPagl_R move | 0.34 |
| SSp-ll_L move vs RSPagl_R move | 0.37 |
| SSp-ll_R move vs RSPagl_R move | 0.017 |
| SSp-n_L move vs RSPagl_R move | 0.64 |
| SSp-n_R move vs RSPagl_R move | 0.85 |
| SSp-tr_L move vs RSPagl_R move | 1 |
| SSp-tr_R move vs RSPagl_R move | 0.88 |
| SSp-ul_L move vs RSPagl_R move | 1 |
| SSp-ul_R move vs RSPagl_R move | 1 |
| SSp-un_L move vs RSPagl_R move | 1 |
| SSp-un_R move vs RSPagl_R move | 1 |
| VISa_L move vs RSPagl_R move | 1 |
| VISam_L move vs RSPagl_R move | 1 |
| VISam_R move vs RSPagl_R move | 1 |
| VISa_R move vs RSPagl_R move | 1 |
| VISp_L move vs RSPagl_R move | **5.8e-06** |
| VISpm_L move vs RSPagl_R move | **0.0076** |
| VISpm_R move vs RSPagl_R move | 0.97 |
| VISp_R move vs RSPagl_R move | 0.012 |
| VISrl_L move vs RSPagl_R move | **2.6e-06** |
| VISrl_R move vs RSPagl_R move | **0.0014** |
| CFA_L plan vs RSPagl_R move | **7.1e-06** |
| CFA_R plan vs RSPagl_R move | **1e-06** |
| MOp_L plan vs RSPagl_R move | **4.2e-07** |
| MOp_R plan vs RSPagl_R move | **1.9e-07** |
| MOs_L plan vs RSPagl_R move | 0.022 |
| MOs_R plan vs RSPagl_R move | **0.0011** |
| RFA_L plan vs RSPagl_R move | 0.01 |
| RFA_R plan vs RSPagl_R move | **0.00028** |
| RSPagl_L plan vs RSPagl_R move | **0** |
| RSPagl_R plan vs RSPagl_R move | **5.9e-09** |
| RSPd_L plan vs RSPagl_R move | **3.7e-06** |
| RSPd_R plan vs RSPagl_R move | **1.7e-06** |
| SSp-bfd_L plan vs RSPagl_R move | **0** |

| | |
|---|---|
| SSp-bfd_R plan vs RSPagl_R move | **0** |
| SSp-ll_L plan vs RSPagl_R move | **0.00084** |
| SSp-ll_R plan vs RSPagl_R move | **0.0092** |
| SSp-n_L plan vs RSPagl_R move | **0** |
| SSp-n_R plan vs RSPagl_R move | **0** |
| SSp-tr_L plan vs RSPagl_R move | **1.4e-05** |
| SSp-tr_R plan vs RSPagl_R move | **0.00049** |
| SSp-ul_L plan vs RSPagl_R move | **2.1e-07** |
| SSp-ul_R plan vs RSPagl_R move | **1.9e-06** |
| SSp-un_L plan vs RSPagl_R move | **0** |
| SSp-un_R plan vs RSPagl_R move | **3.6e-09** |
| VISa_L plan vs RSPagl_R move | **9.5e-10** |
| VISam_L plan vs RSPagl_R move | **0** |
| VISam_R plan vs RSPagl_R move | **0** |
| VISa_R plan vs RSPagl_R move | **2.3e-08** |
| VISp_L plan vs RSPagl_R move | **0** |
| VISpm_L plan vs RSPagl_R move | **0** |
| VISpm_R plan vs RSPagl_R move | **0** |
| VISp_R plan vs RSPagl_R move | **0** |
| VISrl_L plan vs RSPagl_R move | **0** |
| VISrl_R plan vs RSPagl_R move | **0** |
| CFA_L rest vs RSPagl_R move | **0** |
| CFA_R rest vs RSPagl_R move | **0** |
| MOp_L rest vs RSPagl_R move | **0** |
| MOp_R rest vs RSPagl_R move | **0** |
| MOs_L rest vs RSPagl_R move | **0** |
| MOs_R rest vs RSPagl_R move | **0** |
| RFA_L rest vs RSPagl_R move | **0** |
| RFA_R rest vs RSPagl_R move | **0** |
| RSPagl_L rest vs RSPagl_R move | **0** |
| RSPagl_R rest vs RSPagl_R move | **0** |
| RSPd_L rest vs RSPagl_R move | **0** |
| RSPd_R rest vs RSPagl_R move | **0** |
| SSp-bfd_L rest vs RSPagl_R move | **0** |
| SSp-bfd_R rest vs RSPagl_R move | **0** |
| SSp-ll_L rest vs RSPagl_R move | **0** |
| SSp-ll_R rest vs RSPagl_R move | **0** |
| SSp-n_L rest vs RSPagl_R move | **0** |
| SSp-n_R rest vs RSPagl_R move | **0** |
| SSp-tr_L rest vs RSPagl_R move | **0** |
| SSp-tr_R rest vs RSPagl_R move | **0** |
| SSp-ul_L rest vs RSPagl_R move | **0** |
| SSp-ul_R rest vs RSPagl_R move | **0** |
| SSp-un_L rest vs RSPagl_R move | **0** |
| SSp-un_R rest vs RSPagl_R move | **0** |
| VISa_L rest vs RSPagl_R move | **0** |

| | |
|---|---|
| VISam_L rest vs RSPagl_R move | **0** |
| VISam_R rest vs RSPagl_R move | **0** |
| VISa_R rest vs RSPagl_R move | **0** |
| VISp_L rest vs RSPagl_R move | **0** |
| VISpm_L rest vs RSPagl_R move | **0** |
| VISpm_R rest vs RSPagl_R move | **0** |
| VISp_R rest vs RSPagl_R move | **0** |
| VISrl_L rest vs RSPagl_R move | **0** |
| VISrl_R rest vs RSPagl_R move | **0** |
| RSPd_R move vs RSPd_L move | 1 |
| SSp-bfd_L move vs RSPd_L move | **5.8e-08** |
| SSp-bfd_R move vs RSPd_L move | **0.00061** |
| SSp-ll_L move vs RSPd_L move | 1 |
| SSp-ll_R move vs RSPd_L move | 0.92 |
| SSp-n_L move vs RSPd_L move | **0.0031** |
| SSp-n_R move vs RSPd_L move | 0.01 |
| SSp-tr_L move vs RSPd_L move | 1 |
| SSp-tr_R move vs RSPd_L move | 1 |
| SSp-ul_L move vs RSPd_L move | 1 |
| SSp-ul_R move vs RSPd_L move | 1 |
| SSp-un_L move vs RSPd_L move | 0.92 |
| SSp-un_R move vs RSPd_L move | 1 |
| VISa_L move vs RSPd_L move | 0.97 |
| VISam_L move vs RSPd_L move | 0.095 |
| VISam_R move vs RSPd_L move | 0.99 |
| VISa_R move vs RSPd_L move | 1 |
| VISp_L move vs RSPd_L move | **0** |
| VISpm_L move vs RSPd_L move | **1.2e-06** |
| VISpm_R move vs RSPd_L move | 0.033 |
| VISp_R move vs RSPd_L move | **2.3e-06** |
| VISrl_L move vs RSPd_L move | **0** |
| VISrl_R move vs RSPd_L move | **1.1e-07** |
| CFA_L plan vs RSPd_L move | **0** |
| CFA_R plan vs RSPd_L move | **0** |
| MOp_L plan vs RSPd_L move | **0** |
| MOp_R plan vs RSPd_L move | **0** |
| MOs_L plan vs RSPd_L move | **5.6e-06** |
| MOs_R plan vs RSPd_L move | **8.5e-08** |
| RFA_L plan vs RSPd_L move | **1.8e-06** |
| RFA_R plan vs RSPd_L move | **1.3e-08** |
| RSPagl_L plan vs RSPd_L move | **0** |
| RSPagl_R plan vs RSPd_L move | **0** |
| RSPd_L plan vs RSPd_L move | **0** |
| RSPd_R plan vs RSPd_L move | **0** |
| SSp-bfd_L plan vs RSPd_L move | **0** |
| SSp-bfd_R plan vs RSPd_L move | **0** |

| | |
|---|---|
| SSp-ll_L plan vs RSPd_L move | **5.7e-08** |
| SSp-ll_R plan vs RSPd_L move | **1.6e-06** |
| SSp-n_L plan vs RSPd_L move | **0** |
| SSp-n_R plan vs RSPd_L move | **0** |
| SSp-tr_L plan vs RSPd_L move | **0** |
| SSp-tr_R plan vs RSPd_L move | **2.7e-08** |
| SSp-ul_L plan vs RSPd_L move | **0** |
| SSp-ul_R plan vs RSPd_L move | **0** |
| SSp-un_L plan vs RSPd_L move | **0** |
| SSp-un_R plan vs RSPd_L move | **0** |
| VISa_L plan vs RSPd_L move | **0** |
| VISam_L plan vs RSPd_L move | **0** |
| VISam_R plan vs RSPd_L move | **0** |
| VISa_R plan vs RSPd_L move | **0** |
| VISp_L plan vs RSPd_L move | **0** |
| VISpm_L plan vs RSPd_L move | **0** |
| VISpm_R plan vs RSPd_L move | **0** |
| VISp_R plan vs RSPd_L move | **0** |
| VISrl_L plan vs RSPd_L move | **0** |
| VISrl_R plan vs RSPd_L move | **0** |
| CFA_L rest vs RSPd_L move | **0** |
| CFA_R rest vs RSPd_L move | **0** |
| MOp_L rest vs RSPd_L move | **0** |
| MOp_R rest vs RSPd_L move | **0** |
| MOs_L rest vs RSPd_L move | **0** |
| MOs_R rest vs RSPd_L move | **0** |
| RFA_L rest vs RSPd_L move | **0** |
| RFA_R rest vs RSPd_L move | **0** |
| RSPagl_L rest vs RSPd_L move | **0** |
| RSPagl_R rest vs RSPd_L move | **0** |
| RSPd_L rest vs RSPd_L move | **0** |
| RSPd_R rest vs RSPd_L move | **0** |
| SSp-bfd_L rest vs RSPd_L move | **0** |
| SSp-bfd_R rest vs RSPd_L move | **0** |
| SSp-ll_L rest vs RSPd_L move | **0** |
| SSp-ll_R rest vs RSPd_L move | **0** |
| SSp-n_L rest vs RSPd_L move | **0** |
| SSp-n_R rest vs RSPd_L move | **0** |
| SSp-tr_L rest vs RSPd_L move | **0** |
| SSp-tr_R rest vs RSPd_L move | **0** |
| SSp-ul_L rest vs RSPd_L move | **0** |
| SSp-ul_R rest vs RSPd_L move | **0** |
| SSp-un_L rest vs RSPd_L move | **0** |
| SSp-un_R rest vs RSPd_L move | **0** |
| VISa_L rest vs RSPd_L move | **0** |
| VISam_L rest vs RSPd_L move | **0** |

| | |
|---|---|
| VISam_R rest vs RSPd_L move | **0** |
| VISa_R rest vs RSPd_L move | **0** |
| VISp_L rest vs RSPd_L move | **0** |
| VISpm_L rest vs RSPd_L move | **0** |
| VISpm_R rest vs RSPd_L move | **0** |
| VISp_R rest vs RSPd_L move | **0** |
| VISrl_L rest vs RSPd_L move | **0** |
| VISrl_R rest vs RSPd_L move | **0** |
| SSp-bfd_L move vs RSPd_R move | **2.3e-07** |
| SSp-bfd_R move vs RSPd_R move | **0.0017** |
| SSp-ll_L move vs RSPd_R move | 1 |
| SSp-ll_R move vs RSPd_R move | 0.8 |
| SSp-n_L move vs RSPd_R move | **0.008** |
| SSp-n_R move vs RSPd_R move | 0.024 |
| SSp-tr_L move vs RSPd_R move | 1 |
| SSp-tr_R move vs RSPd_R move | 1 |
| SSp-ul_L move vs RSPd_R move | 1 |
| SSp-ul_R move vs RSPd_R move | 1 |
| SSp-un_L move vs RSPd_R move | 0.98 |
| SSp-un_R move vs RSPd_R move | 1 |
| VISa_L move vs RSPd_R move | 1 |
| VISam_L move vs RSPd_R move | 0.18 |
| VISam_R move vs RSPd_R move | 1 |
| VISa_R move vs RSPd_R move | 1 |
| VISp_L move vs RSPd_R move | **0** |
| VISpm_L move vs RSPd_R move | **4.2e-06** |
| VISpm_R move vs RSPd_R move | 0.07 |
| VISp_R move vs RSPd_R move | **8e-06** |
| VISrl_L move vs RSPd_R move | **0** |
| VISrl_R move vs RSPd_R move | **4.3e-07** |
| CFA_L plan vs RSPd_R move | **0** |
| CFA_R plan vs RSPd_R move | **0** |
| MOp_L plan vs RSPd_R move | **0** |
| MOp_R plan vs RSPd_R move | **0** |
| MOs_L plan vs RSPd_R move | **1.9e-05** |
| MOs_R plan vs RSPd_R move | **3.3e-07** |
| RFA_L plan vs RSPd_R move | **6.4e-06** |
| RFA_R plan vs RSPd_R move | **5.5e-08** |
| RSPagl_L plan vs RSPd_R move | **0** |
| RSPagl_R plan vs RSPd_R move | **0** |
| RSPd_L plan vs RSPd_R move | **0** |
| RSPd_R plan vs RSPd_R move | **0** |
| SSp-bfd_L plan vs RSPd_R move | **0** |
| SSp-bfd_R plan vs RSPd_R move | **0** |
| SSp-ll_L plan vs RSPd_R move | **2.3e-07** |
| SSp-ll_R plan vs RSPd_R move | **5.5e-06** |

| | |
|---|---|
| SSp-n_L plan vs RSPd_R move | **0** |
| SSp-n_R plan vs RSPd_R move | **0** |
| SSp-tr_L plan vs RSPd_R move | **3.3e-10** |
| SSp-tr_R plan vs RSPd_R move | **1.1e-07** |
| SSp-ul_L plan vs RSPd_R move | **0** |
| SSp-ul_R plan vs RSPd_R move | **0** |
| SSp-un_L plan vs RSPd_R move | **0** |
| SSp-un_R plan vs RSPd_R move | **0** |
| VISa_L plan vs RSPd_R move | **0** |
| VISam_L plan vs RSPd_R move | **0** |
| VISam_R plan vs RSPd_R move | **0** |
| VISa_R plan vs RSPd_R move | **0** |
| VISp_L plan vs RSPd_R move | **0** |
| VISpm_L plan vs RSPd_R move | **0** |
| VISpm_R plan vs RSPd_R move | **0** |
| VISp_R plan vs RSPd_R move | **0** |
| VISrl_L plan vs RSPd_R move | **0** |
| VISrl_R plan vs RSPd_R move | **0** |
| CFA_L rest vs RSPd_R move | **0** |
| CFA_R rest vs RSPd_R move | **0** |
| MOp_L rest vs RSPd_R move | **0** |
| MOp_R rest vs RSPd_R move | **0** |
| MOs_L rest vs RSPd_R move | **0** |
| MOs_R rest vs RSPd_R move | **0** |
| RFA_L rest vs RSPd_R move | **0** |
| RFA_R rest vs RSPd_R move | **0** |
| RSPagl_L rest vs RSPd_R move | **0** |
| RSPagl_R rest vs RSPd_R move | **0** |
| RSPd_L rest vs RSPd_R move | **0** |
| RSPd_R rest vs RSPd_R move | **0** |
| SSp-bfd_L rest vs RSPd_R move | **0** |
| SSp-bfd_R rest vs RSPd_R move | **0** |
| SSp-ll_L rest vs RSPd_R move | **0** |
| SSp-ll_R rest vs RSPd_R move | **0** |
| SSp-n_L rest vs RSPd_R move | **0** |
| SSp-n_R rest vs RSPd_R move | **0** |
| SSp-tr_L rest vs RSPd_R move | **0** |
| SSp-tr_R rest vs RSPd_R move | **0** |
| SSp-ul_L rest vs RSPd_R move | **0** |
| SSp-ul_R rest vs RSPd_R move | **0** |
| SSp-un_L rest vs RSPd_R move | **0** |
| SSp-un_R rest vs RSPd_R move | **0** |
| VISa_L rest vs RSPd_R move | **0** |
| VISam_L rest vs RSPd_R move | **0** |
| VISam_R rest vs RSPd_R move | **0** |
| VISa_R rest vs RSPd_R move | **0** |

| | |
|---|---|
| VISp_L rest vs RSPd_R move | **0** |
| VISpm_L rest vs RSPd_R move | **0** |
| VISpm_R rest vs RSPd_R move | **0** |
| VISp_R rest vs RSPd_R move | **0** |
| VISrl_L rest vs RSPd_R move | **0** |
| VISrl_R rest vs RSPd_R move | **0** |
| SSp-bfd_R move vs SSp-bfd_L move | 1 |
| SSp-ll_L move vs SSp-bfd_L move | **0** |
| SSp-ll_R move vs SSp-bfd_L move | **0** |
| SSp-n_L move vs SSp-bfd_L move | 1 |
| SSp-n_R move vs SSp-bfd_L move | 1 |
| SSp-tr_L move vs SSp-bfd_L move | **2e-09** |
| SSp-tr_R move vs SSp-bfd_L move | **0** |
| SSp-ul_L move vs SSp-bfd_L move | **6.5e-07** |
| SSp-ul_R move vs SSp-bfd_L move | **0** |
| SSp-un_L move vs SSp-bfd_L move | 0.26 |
| SSp-un_R move vs SSp-bfd_L move | **0.00024** |
| VISa_L move vs SSp-bfd_L move | 0.16 |
| VISam_L move vs SSp-bfd_L move | 0.99 |
| VISam_R move vs SSp-bfd_L move | 0.12 |
| VISa_R move vs SSp-bfd_L move | **0.0011** |
| VISp_L move vs SSp-bfd_L move | 1 |
| VISpm_L move vs SSp-bfd_L move | 1 |
| VISpm_R move vs SSp-bfd_L move | 1 |
| VISp_R move vs SSp-bfd_L move | 1 |
| VISrl_L move vs SSp-bfd_L move | 1 |
| VISrl_R move vs SSp-bfd_L move | 1 |
| CFA_L plan vs SSp-bfd_L move | 1 |
| CFA_R plan vs SSp-bfd_L move | 1 |
| MOp_L plan vs SSp-bfd_L move | 1 |
| MOp_R plan vs SSp-bfd_L move | 1 |
| MOs_L plan vs SSp-bfd_L move | 1 |
| MOs_R plan vs SSp-bfd_L move | 1 |
| RFA_L plan vs SSp-bfd_L move | 1 |
| RFA_R plan vs SSp-bfd_L move | 1 |
| RSPagl_L plan vs SSp-bfd_L move | 1 |
| RSPagl_R plan vs SSp-bfd_L move | 1 |
| RSPd_L plan vs SSp-bfd_L move | 1 |
| RSPd_R plan vs SSp-bfd_L move | 1 |
| SSp-bfd_L plan vs SSp-bfd_L move | 0.079 |
| SSp-bfd_R plan vs SSp-bfd_L move | 0.57 |
| SSp-ll_L plan vs SSp-bfd_L move | 1 |
| SSp-ll_R plan vs SSp-bfd_L move | 1 |
| SSp-n_L plan vs SSp-bfd_L move | 0.31 |
| SSp-n_R plan vs SSp-bfd_L move | 0.64 |
| SSp-tr_L plan vs SSp-bfd_L move | 1 |

| | |
|---|---|
| SSp-tr_R plan vs SSp-bfd_L move | 1 |
| SSp-ul_L plan vs SSp-bfd_L move | 1 |
| SSp-ul_R plan vs SSp-bfd_L move | 1 |
| SSp-un_L plan vs SSp-bfd_L move | 1 |
| SSp-un_R plan vs SSp-bfd_L move | 1 |
| VISa_L plan vs SSp-bfd_L move | 1 |
| VISam_L plan vs SSp-bfd_L move | 0.99 |
| VISam_R plan vs SSp-bfd_L move | 1 |
| VISa_R plan vs SSp-bfd_L move | 1 |
| VISp_L plan vs SSp-bfd_L move | **0.004** |
| VISpm_L plan vs SSp-bfd_L move | 0.12 |
| VISpm_R plan vs SSp-bfd_L move | 0.53 |
| VISp_R plan vs SSp-bfd_L move | 0.045 |
| VISrl_L plan vs SSp-bfd_L move | 0.019 |
| VISrl_R plan vs SSp-bfd_L move | 0.17 |
| CFA_L rest vs SSp-bfd_L move | **0** |
| CFA_R rest vs SSp-bfd_L move | **4.5e-09** |
| MOp_L rest vs SSp-bfd_L move | **0** |
| MOp_R rest vs SSp-bfd_L move | **0** |
| MOs_L rest vs SSp-bfd_L move | **0** |
| MOs_R rest vs SSp-bfd_L move | **0** |
| RFA_L rest vs SSp-bfd_L move | **0** |
| RFA_R rest vs SSp-bfd_L move | **0** |
| RSPagl_L rest vs SSp-bfd_L move | **0** |
| RSPagl_R rest vs SSp-bfd_L move | **0** |
| RSPd_L rest vs SSp-bfd_L move | **0** |
| RSPd_R rest vs SSp-bfd_L move | **0** |
| SSp-bfd_L rest vs SSp-bfd_L move | **0** |
| SSp-bfd_R rest vs SSp-bfd_L move | **1.9e-09** |
| SSp-ll_L rest vs SSp-bfd_L move | **0** |
| SSp-ll_R rest vs SSp-bfd_L move | **0** |
| SSp-n_L rest vs SSp-bfd_L move | **1.9e-10** |
| SSp-n_R rest vs SSp-bfd_L move | **7.3e-08** |
| SSp-tr_L rest vs SSp-bfd_L move | **0** |
| SSp-tr_R rest vs SSp-bfd_L move | **0** |
| SSp-ul_L rest vs SSp-bfd_L move | **0** |
| SSp-ul_R rest vs SSp-bfd_L move | **6.5e-09** |
| SSp-un_L rest vs SSp-bfd_L move | **1.2e-09** |
| SSp-un_R rest vs SSp-bfd_L move | **2.6e-08** |
| VISa_L rest vs SSp-bfd_L move | **0** |
| VISam_L rest vs SSp-bfd_L move | **0** |
| VISam_R rest vs SSp-bfd_L move | **0** |
| VISa_R rest vs SSp-bfd_L move | **0** |
| VISp_L rest vs SSp-bfd_L move | **0** |
| VISpm_L rest vs SSp-bfd_L move | **0** |
| VISpm_R rest vs SSp-bfd_L move | **0** |

| | |
|---|---|
| VISp_R rest vs SSp-bfd_L move | **0** |
| VISrl_L rest vs SSp-bfd_L move | **0** |
| VISrl_R rest vs SSp-bfd_L move | **0** |
| SSp-ll_L move vs SSp-bfd_R move | **0** |
| SSp-ll_R move vs SSp-bfd_R move | **0** |
| SSp-n_L move vs SSp-bfd_R move | 1 |
| SSp-n_R move vs SSp-bfd_R move | 1 |
| SSp-tr_L move vs SSp-bfd_R move | **6e-05** |
| SSp-tr_R move vs SSp-bfd_R move | **7e-08** |
| SSp-ul_L move vs SSp-bfd_R move | **0.0036** |
| SSp-ul_R move vs SSp-bfd_R move | **7.6e-06** |
| SSp-un_L move vs SSp-bfd_R move | 1 |
| SSp-un_R move vs SSp-bfd_R move | 0.18 |
| VISa_L move vs SSp-bfd_R move | 1 |
| VISam_L move vs SSp-bfd_R move | 1 |
| VISam_R move vs SSp-bfd_R move | 1 |
| VISa_R move vs SSp-bfd_R move | 0.38 |
| VISp_L move vs SSp-bfd_R move | 1 |
| VISpm_L move vs SSp-bfd_R move | 1 |
| VISpm_R move vs SSp-bfd_R move | 1 |
| VISp_R move vs SSp-bfd_R move | 1 |
| VISrl_L move vs SSp-bfd_R move | 1 |
| VISrl_R move vs SSp-bfd_R move | 1 |
| CFA_L plan vs SSp-bfd_R move | 1 |
| CFA_R plan vs SSp-bfd_R move | 0.99 |
| MOp_L plan vs SSp-bfd_R move | 0.98 |
| MOp_R plan vs SSp-bfd_R move | 0.95 |
| MOs_L plan vs SSp-bfd_R move | 1 |
| MOs_R plan vs SSp-bfd_R move | 1 |
| RFA_L plan vs SSp-bfd_R move | 1 |
| RFA_R plan vs SSp-bfd_R move | 1 |
| RSPagl_L plan vs SSp-bfd_R move | 0.36 |
| RSPagl_R plan vs SSp-bfd_R move | 0.62 |
| RSPd_L plan vs SSp-bfd_R move | 1 |
| RSPd_R plan vs SSp-bfd_R move | 1 |
| SSp-bfd_L plan vs SSp-bfd_R move | **5.9e-05** |
| SSp-bfd_R plan vs SSp-bfd_R move | **0.0029** |
| SSp-ll_L plan vs SSp-bfd_R move | 1 |
| SSp-ll_R plan vs SSp-bfd_R move | 1 |
| SSp-n_L plan vs SSp-bfd_R move | **0.0007** |
| SSp-n_R plan vs SSp-bfd_R move | **0.0043** |
| SSp-tr_L plan vs SSp-bfd_R move | 1 |
| SSp-tr_R plan vs SSp-bfd_R move | 1 |
| SSp-ul_L plan vs SSp-bfd_R move | 0.95 |
| SSp-ul_R plan vs SSp-bfd_R move | 1 |
| SSp-un_L plan vs SSp-bfd_R move | 0.2 |

| | |
|---|---|
| SSp-un_R plan vs SSp-bfd_R move | 0.57 |
| VISa_L plan vs SSp-bfd_R move | 0.46 |
| VISam_L plan vs SSp-bfd_R move | 0.087 |
| VISam_R plan vs SSp-bfd_R move | 0.26 |
| VISa_R plan vs SSp-bfd_R move | 0.78 |
| VISp_L plan vs SSp-bfd_R move | **7.5e-07** |
| VISpm_L plan vs SSp-bfd_R move | **0.00011** |
| VISpm_R plan vs SSp-bfd_R move | **0.0024** |
| VISp_R plan vs SSp-bfd_R move | **2.5e-05** |
| VISrl_L plan vs SSp-bfd_R move | **6.6e-06** |
| VISrl_R plan vs SSp-bfd_R move | **0.00022** |
| CFA_L rest vs SSp-bfd_R move | **0** |
| CFA_R rest vs SSp-bfd_R move | **0** |
| MOp_L rest vs SSp-bfd_R move | **0** |
| MOp_R rest vs SSp-bfd_R move | **0** |
| MOs_L rest vs SSp-bfd_R move | **0** |
| MOs_R rest vs SSp-bfd_R move | **0** |
| RFA_L rest vs SSp-bfd_R move | **0** |
| RFA_R rest vs SSp-bfd_R move | **0** |
| RSPagl_L rest vs SSp-bfd_R move | **0** |
| RSPagl_R rest vs SSp-bfd_R move | **0** |
| RSPd_L rest vs SSp-bfd_R move | **0** |
| RSPd_R rest vs SSp-bfd_R move | **0** |
| SSp-bfd_L rest vs SSp-bfd_R move | **0** |
| SSp-bfd_R rest vs SSp-bfd_R move | **0** |
| SSp-ll_L rest vs SSp-bfd_R move | **0** |
| SSp-ll_R rest vs SSp-bfd_R move | **0** |
| SSp-n_L rest vs SSp-bfd_R move | **0** |
| SSp-n_R rest vs SSp-bfd_R move | **0** |
| SSp-tr_L rest vs SSp-bfd_R move | **0** |
| SSp-tr_R rest vs SSp-bfd_R move | **0** |
| SSp-ul_L rest vs SSp-bfd_R move | **0** |
| SSp-ul_R rest vs SSp-bfd_R move | **0** |
| SSp-un_L rest vs SSp-bfd_R move | **0** |
| SSp-un_R rest vs SSp-bfd_R move | **0** |
| VISa_L rest vs SSp-bfd_R move | **0** |
| VISam_L rest vs SSp-bfd_R move | **0** |
| VISam_R rest vs SSp-bfd_R move | **0** |
| VISa_R rest vs SSp-bfd_R move | **0** |
| VISp_L rest vs SSp-bfd_R move | **0** |
| VISpm_L rest vs SSp-bfd_R move | **0** |
| VISpm_R rest vs SSp-bfd_R move | **0** |
| VISp_R rest vs SSp-bfd_R move | **0** |
| VISrl_L rest vs SSp-bfd_R move | **0** |
| VISrl_R rest vs SSp-bfd_R move | **0** |
| SSp-ll_R move vs SSp-ll_L move | 1 |

| | |
|---|---|
| SSp-n_L move vs SSp-ll_L move | **9.8e-09** |
| SSp-n_R move vs SSp-ll_L move | **6.2e-08** |
| SSp-tr_L move vs SSp-ll_L move | 1 |
| SSp-tr_R move vs SSp-ll_L move | 1 |
| SSp-ul_L move vs SSp-ll_L move | 1 |
| SSp-ul_R move vs SSp-ll_L move | 1 |
| SSp-un_L move vs SSp-ll_L move | **0.0017** |
| SSp-un_R move vs SSp-ll_L move | 0.6 |
| VISa_L move vs SSp-ll_L move | **0.004** |
| VISam_L move vs SSp-ll_L move | **2.6e-06** |
| VISam_R move vs SSp-ll_L move | **0.0059** |
| VISa_R move vs SSp-ll_L move | 0.34 |
| VISp_L move vs SSp-ll_L move | **0** |
| VISpm_L move vs SSp-ll_L move | **0** |
| VISpm_R move vs SSp-ll_L move | **4.1e-07** |
| VISp_R move vs SSp-ll_L move | **0** |
| VISrl_L move vs SSp-ll_L move | **0** |
| VISrl_R move vs SSp-ll_L move | **0** |
| CFA_L plan vs SSp-ll_L move | **0** |
| CFA_R plan vs SSp-ll_L move | **0** |
| MOp_L plan vs SSp-ll_L move | **0** |
| MOp_R plan vs SSp-ll_L move | **0** |
| MOs_L plan vs SSp-ll_L move | **0** |
| MOs_R plan vs SSp-ll_L move | **0** |
| RFA_L plan vs SSp-ll_L move | **0** |
| RFA_R plan vs SSp-ll_L move | **0** |
| RSPagl_L plan vs SSp-ll_L move | **0** |
| RSPagl_R plan vs SSp-ll_L move | **0** |
| RSPd_L plan vs SSp-ll_L move | **0** |
| RSPd_R plan vs SSp-ll_L move | **0** |
| SSp-bfd_L plan vs SSp-ll_L move | **0** |
| SSp-bfd_R plan vs SSp-ll_L move | **0** |
| SSp-ll_L plan vs SSp-ll_L move | **0** |
| SSp-ll_R plan vs SSp-ll_L move | **0** |
| SSp-n_L plan vs SSp-ll_L move | **0** |
| SSp-n_R plan vs SSp-ll_L move | **0** |
| SSp-tr_L plan vs SSp-ll_L move | **0** |
| SSp-tr_R plan vs SSp-ll_L move | **0** |
| SSp-ul_L plan vs SSp-ll_L move | **0** |
| SSp-ul_R plan vs SSp-ll_L move | **0** |
| SSp-un_L plan vs SSp-ll_L move | **0** |
| SSp-un_R plan vs SSp-ll_L move | **0** |
| VISa_L plan vs SSp-ll_L move | **0** |
| VISam_L plan vs SSp-ll_L move | **0** |
| VISam_R plan vs SSp-ll_L move | **0** |
| VISa_R plan vs SSp-ll_L move | **0** |

| | |
|---|---|
| VISp_L plan vs SSp-ll_L move | **0** |
| VISpm_L plan vs SSp-ll_L move | **0** |
| VISpm_R plan vs SSp-ll_L move | **0** |
| VISp_R plan vs SSp-ll_L move | **0** |
| VISrl_L plan vs SSp-ll_L move | **0** |
| VISrl_R plan vs SSp-ll_L move | **0** |
| CFA_L rest vs SSp-ll_L move | **0** |
| CFA_R rest vs SSp-ll_L move | **0** |
| MOp_L rest vs SSp-ll_L move | **0** |
| MOp_R rest vs SSp-ll_L move | **0** |
| MOs_L rest vs SSp-ll_L move | **0** |
| MOs_R rest vs SSp-ll_L move | **0** |
| RFA_L rest vs SSp-ll_L move | **0** |
| RFA_R rest vs SSp-ll_L move | **0** |
| RSPagl_L rest vs SSp-ll_L move | **0** |
| RSPagl_R rest vs SSp-ll_L move | **0** |
| RSPd_L rest vs SSp-ll_L move | **0** |
| RSPd_R rest vs SSp-ll_L move | **0** |
| SSp-bfd_L rest vs SSp-ll_L move | **0** |
| SSp-bfd_R rest vs SSp-ll_L move | **0** |
| SSp-ll_L rest vs SSp-ll_L move | **0** |
| SSp-ll_R rest vs SSp-ll_L move | **0** |
| SSp-n_L rest vs SSp-ll_L move | **0** |
| SSp-n_R rest vs SSp-ll_L move | **0** |
| SSp-tr_L rest vs SSp-ll_L move | **0** |
| SSp-tr_R rest vs SSp-ll_L move | **0** |
| SSp-ul_L rest vs SSp-ll_L move | **0** |
| SSp-ul_R rest vs SSp-ll_L move | **0** |
| SSp-un_L rest vs SSp-ll_L move | **0** |
| SSp-un_R rest vs SSp-ll_L move | **0** |
| VISa_L rest vs SSp-ll_L move | **0** |
| VISam_L rest vs SSp-ll_L move | **0** |
| VISam_R rest vs SSp-ll_L move | **0** |
| VISa_R rest vs SSp-ll_L move | **0** |
| VISp_L rest vs SSp-ll_L move | **0** |
| VISpm_L rest vs SSp-ll_L move | **0** |
| VISpm_R rest vs SSp-ll_L move | **0** |
| VISp_R rest vs SSp-ll_L move | **0** |
| VISrl_L rest vs SSp-ll_L move | **0** |
| VISrl_R rest vs SSp-ll_L move | **0** |
| SSp-n_L move vs SSp-ll_R move | **0** |
| SSp-n_R move vs SSp-ll_R move | **0** |
| SSp-tr_L move vs SSp-ll_R move | 1 |
| SSp-tr_R move vs SSp-ll_R move | 1 |
| SSp-ul_L move vs SSp-ll_R move | 0.66 |
| SSp-ul_R move vs SSp-ll_R move | 1 |

| | |
|---|---|
| SSp-un_L move vs SSp-ll_R move | **1.1e-05** |
| SSp-un_R move vs SSp-ll_R move | 0.046 |
| VISa_L move vs SSp-ll_R move | **3.1e-05** |
| VISam_L move vs SSp-ll_R move | **4.4e-09** |
| VISam_R move vs SSp-ll_R move | **5e-05** |
| VISa_R move vs SSp-ll_R move | 0.014 |
| VISp_L move vs SSp-ll_R move | **0** |
| VISpm_L move vs SSp-ll_R move | **0** |
| VISpm_R move vs SSp-ll_R move | **0** |
| VISp_R move vs SSp-ll_R move | **0** |
| VISrl_L move vs SSp-ll_R move | **0** |
| VISrl_R move vs SSp-ll_R move | **0** |
| CFA_L plan vs SSp-ll_R move | **0** |
| CFA_R plan vs SSp-ll_R move | **0** |
| MOp_L plan vs SSp-ll_R move | **0** |
| MOp_R plan vs SSp-ll_R move | **0** |
| MOs_L plan vs SSp-ll_R move | **0** |
| MOs_R plan vs SSp-ll_R move | **0** |
| RFA_L plan vs SSp-ll_R move | **0** |
| RFA_R plan vs SSp-ll_R move | **0** |
| RSPagl_L plan vs SSp-ll_R move | **0** |
| RSPagl_R plan vs SSp-ll_R move | **0** |
| RSPd_L plan vs SSp-ll_R move | **0** |
| RSPd_R plan vs SSp-ll_R move | **0** |
| SSp-bfd_L plan vs SSp-ll_R move | **0** |
| SSp-bfd_R plan vs SSp-ll_R move | **0** |
| SSp-ll_L plan vs SSp-ll_R move | **0** |
| SSp-ll_R plan vs SSp-ll_R move | **0** |
| SSp-n_L plan vs SSp-ll_R move | **0** |
| SSp-n_R plan vs SSp-ll_R move | **0** |
| SSp-tr_L plan vs SSp-ll_R move | **0** |
| SSp-tr_R plan vs SSp-ll_R move | **0** |
| SSp-ul_L plan vs SSp-ll_R move | **0** |
| SSp-ul_R plan vs SSp-ll_R move | **0** |
| SSp-un_L plan vs SSp-ll_R move | **0** |
| SSp-un_R plan vs SSp-ll_R move | **0** |
| VISa_L plan vs SSp-ll_R move | **0** |
| VISam_L plan vs SSp-ll_R move | **0** |
| VISam_R plan vs SSp-ll_R move | **0** |
| VISa_R plan vs SSp-ll_R move | **0** |
| VISp_L plan vs SSp-ll_R move | **0** |
| VISpm_L plan vs SSp-ll_R move | **0** |
| VISpm_R plan vs SSp-ll_R move | **0** |
| VISp_R plan vs SSp-ll_R move | **0** |
| VISrl_L plan vs SSp-ll_R move | **0** |
| VISrl_R plan vs SSp-ll_R move | **0** |

| | |
|---|---|
| CFA_L rest vs SSp-ll_R move | **0** |
| CFA_R rest vs SSp-ll_R move | **0** |
| MOp_L rest vs SSp-ll_R move | **0** |
| MOp_R rest vs SSp-ll_R move | **0** |
| MOs_L rest vs SSp-ll_R move | **0** |
| MOs_R rest vs SSp-ll_R move | **0** |
| RFA_L rest vs SSp-ll_R move | **0** |
| RFA_R rest vs SSp-ll_R move | **0** |
| RSPagl_L rest vs SSp-ll_R move | **0** |
| RSPagl_R rest vs SSp-ll_R move | **0** |
| RSPd_L rest vs SSp-ll_R move | **0** |
| RSPd_R rest vs SSp-ll_R move | **0** |
| SSp-bfd_L rest vs SSp-ll_R move | **0** |
| SSp-bfd_R rest vs SSp-ll_R move | **0** |
| SSp-ll_L rest vs SSp-ll_R move | **0** |
| SSp-ll_R rest vs SSp-ll_R move | **0** |
| SSp-n_L rest vs SSp-ll_R move | **0** |
| SSp-n_R rest vs SSp-ll_R move | **0** |
| SSp-tr_L rest vs SSp-ll_R move | **0** |
| SSp-tr_R rest vs SSp-ll_R move | **0** |
| SSp-ul_L rest vs SSp-ll_R move | **0** |
| SSp-ul_R rest vs SSp-ll_R move | **0** |
| SSp-un_L rest vs SSp-ll_R move | **0** |
| SSp-un_R rest vs SSp-ll_R move | **0** |
| VISa_L rest vs SSp-ll_R move | **0** |
| VISam_L rest vs SSp-ll_R move | **0** |
| VISam_R rest vs SSp-ll_R move | **0** |
| VISa_R rest vs SSp-ll_R move | **0** |
| VISp_L rest vs SSp-ll_R move | **0** |
| VISpm_L rest vs SSp-ll_R move | **0** |
| VISpm_R rest vs SSp-ll_R move | **0** |
| VISp_R rest vs SSp-ll_R move | **0** |
| VISrl_L rest vs SSp-ll_R move | **0** |
| VISrl_R rest vs SSp-ll_R move | **0** |
| SSp-n_R move vs SSp-n_L move | 1 |
| SSp-tr_L move vs SSp-n_L move | **0.00037** |
| SSp-tr_R move vs SSp-n_L move | **6.4e-07** |
| SSp-ul_L move vs SSp-n_L move | 0.016 |
| SSp-ul_R move vs SSp-n_L move | **5.3e-05** |
| SSp-un_L move vs SSp-n_L move | 1 |
| SSp-un_R move vs SSp-n_L move | 0.42 |
| VISa_L move vs SSp-n_L move | 1 |
| VISam_L move vs SSp-n_L move | 1 |
| VISam_R move vs SSp-n_L move | 1 |
| VISa_R move vs SSp-n_L move | 0.68 |
| VISp_L move vs SSp-n_L move | 0.99 |

| | |
|---|---|
| VISpm_L move vs SSp-n_L move | 1 |
| VISpm_R move vs SSp-n_L move | 1 |
| VISp_R move vs SSp-n_L move | 1 |
| VISrl_L move vs SSp-n_L move | 0.97 |
| VISrl_R move vs SSp-n_L move | 1 |
| CFA_L plan vs SSp-n_L move | 0.99 |
| CFA_R plan vs SSp-n_L move | 0.92 |
| MOp_L plan vs SSp-n_L move | 0.84 |
| MOp_R plan vs SSp-n_L move | 0.75 |
| MOs_L plan vs SSp-n_L move | 1 |
| MOs_R plan vs SSp-n_L move | 1 |
| RFA_L plan vs SSp-n_L move | 1 |
| RFA_R plan vs SSp-n_L move | 1 |
| RSPagl_L plan vs SSp-n_L move | 0.15 |
| RSPagl_R plan vs SSp-n_L move | 0.32 |
| RSPd_L plan vs SSp-n_L move | 0.98 |
| RSPd_R plan vs SSp-n_L move | 0.95 |
| SSp-bfd_L plan vs SSp-n_L move | **8.5e-06** |
| SSp-bfd_R plan vs SSp-n_L move | **0.00056** |
| SSp-ll_L plan vs SSp-n_L move | 1 |
| SSp-ll_R plan vs SSp-n_L move | 1 |
| SSp-n_L plan vs SSp-n_L move | **0.00012** |
| SSp-n_R plan vs SSp-n_L move | **0.00085** |
| SSp-tr_L plan vs SSp-n_L move | 1 |
| SSp-tr_R plan vs SSp-n_L move | 1 |
| SSp-ul_L plan vs SSp-n_L move | 0.77 |
| SSp-ul_R plan vs SSp-n_L move | 0.96 |
| SSp-un_L plan vs SSp-n_L move | 0.067 |
| SSp-un_R plan vs SSp-n_L move | 0.28 |
| VISa_L plan vs SSp-n_L move | 0.2 |
| VISam_L plan vs SSp-n_L move | 0.025 |
| VISam_R plan vs SSp-n_L move | 0.097 |
| VISa_R plan vs SSp-n_L move | 0.48 |
| VISp_L plan vs SSp-n_L move | **8.4e-08** |
| VISpm_L plan vs SSp-n_L move | **1.7e-05** |
| VISpm_R plan vs SSp-n_L move | **0.00045** |
| VISp_R plan vs SSp-n_L move | **3.4e-06** |
| VISrl_L plan vs SSp-n_L move | **8.3e-07** |
| VISrl_R plan vs SSp-n_L move | **3.5e-05** |
| CFA_L rest vs SSp-n_L move | **0** |
| CFA_R rest vs SSp-n_L move | **0** |
| MOp_L rest vs SSp-n_L move | **0** |
| MOp_R rest vs SSp-n_L move | **0** |
| MOs_L rest vs SSp-n_L move | **0** |
| MOs_R rest vs SSp-n_L move | **0** |
| RFA_L rest vs SSp-n_L move | **0** |

| | |
|---|---|
| RFA_R rest vs SSp-n_L move | **0** |
| RSPagl_L rest vs SSp-n_L move | **0** |
| RSPagl_R rest vs SSp-n_L move | **0** |
| RSPd_L rest vs SSp-n_L move | **0** |
| RSPd_R rest vs SSp-n_L move | **0** |
| SSp-bfd_L rest vs SSp-n_L move | **0** |
| SSp-bfd_R rest vs SSp-n_L move | **0** |
| SSp-ll_L rest vs SSp-n_L move | **0** |
| SSp-ll_R rest vs SSp-n_L move | **0** |
| SSp-n_L rest vs SSp-n_L move | **0** |
| SSp-n_R rest vs SSp-n_L move | **0** |
| SSp-tr_L rest vs SSp-n_L move | **0** |
| SSp-tr_R rest vs SSp-n_L move | **0** |
| SSp-ul_L rest vs SSp-n_L move | **0** |
| SSp-ul_R rest vs SSp-n_L move | **0** |
| SSp-un_L rest vs SSp-n_L move | **0** |
| SSp-un_R rest vs SSp-n_L move | **0** |
| VISa_L rest vs SSp-n_L move | **0** |
| VISam_L rest vs SSp-n_L move | **0** |
| VISam_R rest vs SSp-n_L move | **0** |
| VISa_R rest vs SSp-n_L move | **0** |
| VISp_L rest vs SSp-n_L move | **0** |
| VISpm_L rest vs SSp-n_L move | **0** |
| VISpm_R rest vs SSp-n_L move | **0** |
| VISp_R rest vs SSp-n_L move | **0** |
| VISrl_L rest vs SSp-n_L move | **0** |
| VISrl_R rest vs SSp-n_L move | **0** |
| SSp-tr_L move vs SSp-n_R move | **0.0014** |
| SSp-tr_R move vs SSp-n_R move | **3.2e-06** |
| SSp-ul_L move vs SSp-n_R move | 0.044 |
| SSp-ul_R move vs SSp-n_R move | **0.00022** |
| SSp-un_L move vs SSp-n_R move | 1 |
| SSp-un_R move vs SSp-n_R move | 0.66 |
| VISa_L move vs SSp-n_R move | 1 |
| VISam_L move vs SSp-n_R move | 1 |
| VISam_R move vs SSp-n_R move | 1 |
| VISa_R move vs SSp-n_R move | 0.88 |
| VISp_L move vs SSp-n_R move | 0.93 |
| VISpm_L move vs SSp-n_R move | 1 |
| VISpm_R move vs SSp-n_R move | 1 |
| VISp_R move vs SSp-n_R move | 1 |
| VISrl_L move vs SSp-n_R move | 0.87 |
| VISrl_R move vs SSp-n_R move | 1 |
| CFA_L plan vs SSp-n_R move | 0.94 |
| CFA_R plan vs SSp-n_R move | 0.76 |
| MOp_L plan vs SSp-n_R move | 0.64 |

| | |
|---|---|
| MOp_R plan vs SSp-n_R move | 0.52 |
| MOs_L plan vs SSp-n_R move | 1 |
| MOs_R plan vs SSp-n_R move | 1 |
| RFA_L plan vs SSp-n_R move | 1 |
| RFA_R plan vs SSp-n_R move | 1 |
| RSPagl_L plan vs SSp-n_R move | 0.062 |
| RSPagl_R plan vs SSp-n_R move | 0.16 |
| RSPd_L plan vs SSp-n_R move | 0.9 |
| RSPd_R plan vs SSp-n_R move | 0.82 |
| SSp-bfd_L plan vs SSp-n_R move | **1.8e-06** |
| SSp-bfd_R plan vs SSp-n_R move | **0.00014** |
| SSp-ll_L plan vs SSp-n_R move | 1 |
| SSp-ll_R plan vs SSp-n_R move | 1 |
| SSp-n_L plan vs SSp-n_R move | **2.8e-05** |
| SSp-n_R plan vs SSp-n_R move | **0.00022** |
| SSp-tr_L plan vs SSp-n_R move | 0.97 |
| SSp-tr_R plan vs SSp-n_R move | 1 |
| SSp-ul_L plan vs SSp-n_R move | 0.54 |
| SSp-ul_R plan vs SSp-n_R move | 0.83 |
| SSp-un_L plan vs SSp-n_R move | 0.025 |
| SSp-un_R plan vs SSp-n_R move | 0.13 |
| VISa_L plan vs SSp-n_R move | 0.091 |
| VISam_L plan vs SSp-n_R move | **0.0085** |
| VISam_R plan vs SSp-n_R move | 0.039 |
| VISa_R plan vs SSp-n_R move | 0.27 |
| VISp_L plan vs SSp-n_R move | **1.4e-08** |
| VISpm_L plan vs SSp-n_R move | **3.7e-06** |
| VISpm_R plan vs SSp-n_R move | **0.00012** |
| VISp_R plan vs SSp-n_R move | **6.7e-07** |
| VISrl_L plan vs SSp-n_R move | **1.6e-07** |
| VISrl_R plan vs SSp-n_R move | **7.8e-06** |
| CFA_L rest vs SSp-n_R move | **0** |
| CFA_R rest vs SSp-n_R move | **0** |
| MOp_L rest vs SSp-n_R move | **0** |
| MOp_R rest vs SSp-n_R move | **0** |
| MOs_L rest vs SSp-n_R move | **0** |
| MOs_R rest vs SSp-n_R move | **0** |
| RFA_L rest vs SSp-n_R move | **0** |
| RFA_R rest vs SSp-n_R move | **0** |
| RSPagl_L rest vs SSp-n_R move | **0** |
| RSPagl_R rest vs SSp-n_R move | **0** |
| RSPd_L rest vs SSp-n_R move | **0** |
| RSPd_R rest vs SSp-n_R move | **0** |
| SSp-bfd_L rest vs SSp-n_R move | **0** |
| SSp-bfd_R rest vs SSp-n_R move | **0** |
| SSp-ll_L rest vs SSp-n_R move | **0** |

| | |
|---|---|
| SSp-ll_R rest vs SSp-n_R move | **0** |
| SSp-n_L rest vs SSp-n_R move | **0** |
| SSp-n_R rest vs SSp-n_R move | **0** |
| SSp-tr_L rest vs SSp-n_R move | **0** |
| SSp-tr_R rest vs SSp-n_R move | **0** |
| SSp-ul_L rest vs SSp-n_R move | **0** |
| SSp-ul_R rest vs SSp-n_R move | **0** |
| SSp-un_L rest vs SSp-n_R move | **0** |
| SSp-un_R rest vs SSp-n_R move | **0** |
| VISa_L rest vs SSp-n_R move | **0** |
| VISam_L rest vs SSp-n_R move | **0** |
| VISam_R rest vs SSp-n_R move | **0** |
| VISa_R rest vs SSp-n_R move | **0** |
| VISp_L rest vs SSp-n_R move | **0** |
| VISpm_L rest vs SSp-n_R move | **0** |
| VISpm_R rest vs SSp-n_R move | **0** |
| VISp_R rest vs SSp-n_R move | **0** |
| VISrl_L rest vs SSp-n_R move | **0** |
| VISrl_R rest vs SSp-n_R move | **0** |
| SSp-tr_R move vs SSp-tr_L move | 1 |
| SSp-ul_L move vs SSp-tr_L move | 1 |
| SSp-ul_R move vs SSp-tr_L move | 1 |
| SSp-un_L move vs SSp-tr_L move | 0.61 |
| SSp-un_R move vs SSp-tr_L move | 1 |
| VISa_L move vs SSp-tr_L move | 0.76 |
| VISam_L move vs SSp-tr_L move | 0.019 |
| VISam_R move vs SSp-tr_L move | 0.83 |
| VISa_R move vs SSp-tr_L move | 1 |
| VISp_L move vs SSp-tr_L move | **0** |
| VISpm_L move vs SSp-tr_L move | **7.2e-08** |
| VISpm_R move vs SSp-tr_L move | **0.0053** |
| VISp_R move vs SSp-tr_L move | **1.5e-07** |
| VISrl_L move vs SSp-tr_L move | **0** |
| VISrl_R move vs SSp-tr_L move | **5e-09** |
| CFA_L plan vs SSp-tr_L move | **0** |
| CFA_R plan vs SSp-tr_L move | **0** |
| MOp_L plan vs SSp-tr_L move | **0** |
| MOp_R plan vs SSp-tr_L move | **0** |
| MOs_L plan vs SSp-tr_L move | **3.8e-07** |
| MOs_R plan vs SSp-tr_L move | **3.4e-09** |
| RFA_L plan vs SSp-tr_L move | **1.1e-07** |
| RFA_R plan vs SSp-tr_L move | **0** |
| RSPagl_L plan vs SSp-tr_L move | **0** |
| RSPagl_R plan vs SSp-tr_L move | **0** |
| RSPd_L plan vs SSp-tr_L move | **0** |
| RSPd_R plan vs SSp-tr_L move | **0** |

| | |
|---|---|
| SSp-bfd_L plan vs SSp-tr_L move | **0** |
| SSp-bfd_R plan vs SSp-tr_L move | **0** |
| SSp-ll_L plan vs SSp-tr_L move | **1.9e-09** |
| SSp-ll_R plan vs SSp-tr_L move | **9.6e-08** |
| SSp-n_L plan vs SSp-tr_L move | **0** |
| SSp-n_R plan vs SSp-tr_L move | **0** |
| SSp-tr_L plan vs SSp-tr_L move | **0** |
| SSp-tr_R plan vs SSp-tr_L move | **3.3e-10** |
| SSp-ul_L plan vs SSp-tr_L move | **0** |
| SSp-ul_R plan vs SSp-tr_L move | **0** |
| SSp-un_L plan vs SSp-tr_L move | **0** |
| SSp-un_R plan vs SSp-tr_L move | **0** |
| VISa_L plan vs SSp-tr_L move | **0** |
| VISam_L plan vs SSp-tr_L move | **0** |
| VISam_R plan vs SSp-tr_L move | **0** |
| VISa_R plan vs SSp-tr_L move | **0** |
| VISp_L plan vs SSp-tr_L move | **0** |
| VISpm_L plan vs SSp-tr_L move | **0** |
| VISpm_R plan vs SSp-tr_L move | **0** |
| VISp_R plan vs SSp-tr_L move | **0** |
| VISrl_L plan vs SSp-tr_L move | **0** |
| VISrl_R plan vs SSp-tr_L move | **0** |
| CFA_L rest vs SSp-tr_L move | **0** |
| CFA_R rest vs SSp-tr_L move | **0** |
| MOp_L rest vs SSp-tr_L move | **0** |
| MOp_R rest vs SSp-tr_L move | **0** |
| MOs_L rest vs SSp-tr_L move | **0** |
| MOs_R rest vs SSp-tr_L move | **0** |
| RFA_L rest vs SSp-tr_L move | **0** |
| RFA_R rest vs SSp-tr_L move | **0** |
| RSPagl_L rest vs SSp-tr_L move | **0** |
| RSPagl_R rest vs SSp-tr_L move | **0** |
| RSPd_L rest vs SSp-tr_L move | **0** |
| RSPd_R rest vs SSp-tr_L move | **0** |
| SSp-bfd_L rest vs SSp-tr_L move | **0** |
| SSp-bfd_R rest vs SSp-tr_L move | **0** |
| SSp-ll_L rest vs SSp-tr_L move | **0** |
| SSp-ll_R rest vs SSp-tr_L move | **0** |
| SSp-n_L rest vs SSp-tr_L move | **0** |
| SSp-n_R rest vs SSp-tr_L move | **0** |
| SSp-tr_L rest vs SSp-tr_L move | **0** |
| SSp-tr_R rest vs SSp-tr_L move | **0** |
| SSp-ul_L rest vs SSp-tr_L move | **0** |
| SSp-ul_R rest vs SSp-tr_L move | **0** |
| SSp-un_L rest vs SSp-tr_L move | **0** |
| SSp-un_R rest vs SSp-tr_L move | **0** |

| | |
|---|---|
| VISa_L rest vs SSp-tr_L move | **0** |
| VISam_L rest vs SSp-tr_L move | **0** |
| VISam_R rest vs SSp-tr_L move | **0** |
| VISa_R rest vs SSp-tr_L move | **0** |
| VISp_L rest vs SSp-tr_L move | **0** |
| VISpm_L rest vs SSp-tr_L move | **0** |
| VISpm_R rest vs SSp-tr_L move | **0** |
| VISp_R rest vs SSp-tr_L move | **0** |
| VISrl_L rest vs SSp-tr_L move | **0** |
| VISrl_R rest vs SSp-tr_L move | **0** |
| SSp-ul_L move vs SSp-tr_R move | 1 |
| SSp-ul_R move vs SSp-tr_R move | 1 |
| SSp-un_L move vs SSp-tr_R move | 0.026 |
| SSp-un_R move vs SSp-tr_R move | 0.97 |
| VISa_L move vs SSp-tr_R move | 0.052 |
| VISam_L move vs SSp-tr_R move | **9.4e-05** |
| VISam_R move vs SSp-tr_R move | 0.07 |
| VISa_R move vs SSp-tr_R move | 0.86 |
| VISp_L move vs SSp-tr_R move | **0** |
| VISpm_L move vs SSp-tr_R move | **0** |
| VISpm_R move vs SSp-tr_R move | **1.8e-05** |
| VISp_R move vs SSp-tr_R move | **0** |
| VISrl_L move vs SSp-tr_R move | **0** |
| VISrl_R move vs SSp-tr_R move | **0** |
| CFA_L plan vs SSp-tr_R move | **0** |
| CFA_R plan vs SSp-tr_R move | **0** |
| MOp_L plan vs SSp-tr_R move | **0** |
| MOp_R plan vs SSp-tr_R move | **0** |
| MOs_L plan vs SSp-tr_R move | **0** |
| MOs_R plan vs SSp-tr_R move | **0** |
| RFA_L plan vs SSp-tr_R move | **0** |
| RFA_R plan vs SSp-tr_R move | **0** |
| RSPagl_L plan vs SSp-tr_R move | **0** |
| RSPagl_R plan vs SSp-tr_R move | **0** |
| RSPd_L plan vs SSp-tr_R move | **0** |
| RSPd_R plan vs SSp-tr_R move | **0** |
| SSp-bfd_L plan vs SSp-tr_R move | **0** |
| SSp-bfd_R plan vs SSp-tr_R move | **0** |
| SSp-ll_L plan vs SSp-tr_R move | **0** |
| SSp-ll_R plan vs SSp-tr_R move | **0** |
| SSp-n_L plan vs SSp-tr_R move | **0** |
| SSp-n_R plan vs SSp-tr_R move | **0** |
| SSp-tr_L plan vs SSp-tr_R move | **0** |
| SSp-tr_R plan vs SSp-tr_R move | **0** |
| SSp-ul_L plan vs SSp-tr_R move | **0** |
| SSp-ul_R plan vs SSp-tr_R move | **0** |

| | |
|---|---|
| SSp-un_L plan vs SSp-tr_R move | **0** |
| SSp-un_R plan vs SSp-tr_R move | **0** |
| VISa_L plan vs SSp-tr_R move | **0** |
| VISam_L plan vs SSp-tr_R move | **0** |
| VISam_R plan vs SSp-tr_R move | **0** |
| VISa_R plan vs SSp-tr_R move | **0** |
| VISp_L plan vs SSp-tr_R move | **0** |
| VISpm_L plan vs SSp-tr_R move | **0** |
| VISpm_R plan vs SSp-tr_R move | **0** |
| VISp_R plan vs SSp-tr_R move | **0** |
| VISrl_L plan vs SSp-tr_R move | **0** |
| VISrl_R plan vs SSp-tr_R move | **0** |
| CFA_L rest vs SSp-tr_R move | **0** |
| CFA_R rest vs SSp-tr_R move | **0** |
| MOp_L rest vs SSp-tr_R move | **0** |
| MOp_R rest vs SSp-tr_R move | **0** |
| MOs_L rest vs SSp-tr_R move | **0** |
| MOs_R rest vs SSp-tr_R move | **0** |
| RFA_L rest vs SSp-tr_R move | **0** |
| RFA_R rest vs SSp-tr_R move | **0** |
| RSPagl_L rest vs SSp-tr_R move | **0** |
| RSPagl_R rest vs SSp-tr_R move | **0** |
| RSPd_L rest vs SSp-tr_R move | **0** |
| RSPd_R rest vs SSp-tr_R move | **0** |
| SSp-bfd_L rest vs SSp-tr_R move | **0** |
| SSp-bfd_R rest vs SSp-tr_R move | **0** |
| SSp-ll_L rest vs SSp-tr_R move | **0** |
| SSp-ll_R rest vs SSp-tr_R move | **0** |
| SSp-n_L rest vs SSp-tr_R move | **0** |
| SSp-n_R rest vs SSp-tr_R move | **0** |
| SSp-tr_L rest vs SSp-tr_R move | **0** |
| SSp-tr_R rest vs SSp-tr_R move | **0** |
| SSp-ul_L rest vs SSp-tr_R move | **0** |
| SSp-ul_R rest vs SSp-tr_R move | **0** |
| SSp-un_L rest vs SSp-tr_R move | **0** |
| SSp-un_R rest vs SSp-tr_R move | **0** |
| VISa_L rest vs SSp-tr_R move | **0** |
| VISam_L rest vs SSp-tr_R move | **0** |
| VISam_R rest vs SSp-tr_R move | **0** |
| VISa_R rest vs SSp-tr_R move | **0** |
| VISp_L rest vs SSp-tr_R move | **0** |
| VISpm_L rest vs SSp-tr_R move | **0** |
| VISpm_R rest vs SSp-tr_R move | **0** |
| VISp_R rest vs SSp-tr_R move | **0** |
| VISrl_L rest vs SSp-tr_R move | **0** |
| VISrl_R rest vs SSp-tr_R move | **0** |

| | |
|---|---|
| SSp-ul_R move vs SSp-ul_L move | 1 |
| SSp-un_L move vs SSp-ul_L move | 0.99 |
| SSp-un_R move vs SSp-ul_L move | 1 |
| VISa_L move vs SSp-ul_L move | 1 |
| VISam_L move vs SSp-ul_L move | 0.28 |
| VISam_R move vs SSp-ul_L move | 1 |
| VISa_R move vs SSp-ul_L move | 1 |
| VISp_L move vs SSp-ul_L move | **4.1e-10** |
| VISpm_L move vs SSp-ul_L move | **1.1e-05** |
| VISpm_R move vs SSp-ul_L move | 0.12 |
| VISp_R move vs SSp-ul_L move | **2.1e-05** |
| VISrl_L move vs SSp-ul_L move | **0** |
| VISrl_R move vs SSp-ul_L move | **1.2e-06** |
| CFA_L plan vs SSp-ul_L move | **7.9e-10** |
| CFA_R plan vs SSp-ul_L move | **0** |
| MOp_L plan vs SSp-ul_L move | **0** |
| MOp_R plan vs SSp-ul_L move | **0** |
| MOs_L plan vs SSp-ul_L move | **4.7e-05** |
| MOs_R plan vs SSp-ul_L move | **9.3e-07** |
| RFA_L plan vs SSp-ul_L move | **1.7e-05** |
| RFA_R plan vs SSp-ul_L move | **1.6e-07** |
| RSPagl_L plan vs SSp-ul_L move | **0** |
| RSPagl_R plan vs SSp-ul_L move | **0** |
| RSPd_L plan vs SSp-ul_L move | **0** |
| RSPd_R plan vs SSp-ul_L move | **0** |
| SSp-bfd_L plan vs SSp-ul_L move | **0** |
| SSp-bfd_R plan vs SSp-ul_L move | **0** |
| SSp-ll_L plan vs SSp-ul_L move | **6.4e-07** |
| SSp-ll_R plan vs SSp-ul_L move | **1.4e-05** |
| SSp-n_L plan vs SSp-ul_L move | **0** |
| SSp-n_R plan vs SSp-ul_L move | **0** |
| SSp-tr_L plan vs SSp-ul_L move | **3.3e-09** |
| SSp-tr_R plan vs SSp-ul_L move | **3.2e-07** |
| SSp-ul_L plan vs SSp-ul_L move | **0** |
| SSp-ul_R plan vs SSp-ul_L move | **0** |
| SSp-un_L plan vs SSp-ul_L move | **0** |
| SSp-un_R plan vs SSp-ul_L move | **0** |
| VISa_L plan vs SSp-ul_L move | **0** |
| VISam_L plan vs SSp-ul_L move | **0** |
| VISam_R plan vs SSp-ul_L move | **0** |
| VISa_R plan vs SSp-ul_L move | **0** |
| VISp_L plan vs SSp-ul_L move | **0** |
| VISpm_L plan vs SSp-ul_L move | **0** |
| VISpm_R plan vs SSp-ul_L move | **0** |
| VISp_R plan vs SSp-ul_L move | **0** |
| VISrl_L plan vs SSp-ul_L move | **0** |

| | |
|---|---|
| VISrl_R plan vs SSp-ul_L move | **0** |
| CFA_L rest vs SSp-ul_L move | **0** |
| CFA_R rest vs SSp-ul_L move | **0** |
| MOp_L rest vs SSp-ul_L move | **0** |
| MOp_R rest vs SSp-ul_L move | **0** |
| MOs_L rest vs SSp-ul_L move | **0** |
| MOs_R rest vs SSp-ul_L move | **0** |
| RFA_L rest vs SSp-ul_L move | **0** |
| RFA_R rest vs SSp-ul_L move | **0** |
| RSPagl_L rest vs SSp-ul_L move | **0** |
| RSPagl_R rest vs SSp-ul_L move | **0** |
| RSPd_L rest vs SSp-ul_L move | **0** |
| RSPd_R rest vs SSp-ul_L move | **0** |
| SSp-bfd_L rest vs SSp-ul_L move | **0** |
| SSp-bfd_R rest vs SSp-ul_L move | **0** |
| SSp-ll_L rest vs SSp-ul_L move | **0** |
| SSp-ll_R rest vs SSp-ul_L move | **0** |
| SSp-n_L rest vs SSp-ul_L move | **0** |
| SSp-n_R rest vs SSp-ul_L move | **0** |
| SSp-tr_L rest vs SSp-ul_L move | **0** |
| SSp-tr_R rest vs SSp-ul_L move | **0** |
| SSp-ul_L rest vs SSp-ul_L move | **0** |
| SSp-ul_R rest vs SSp-ul_L move | **0** |
| SSp-un_L rest vs SSp-ul_L move | **0** |
| SSp-un_R rest vs SSp-ul_L move | **0** |
| VISa_L rest vs SSp-ul_L move | **0** |
| VISam_L rest vs SSp-ul_L move | **0** |
| VISam_R rest vs SSp-ul_L move | **0** |
| VISa_R rest vs SSp-ul_L move | **0** |
| VISp_L rest vs SSp-ul_L move | **0** |
| VISpm_L rest vs SSp-ul_L move | **0** |
| VISpm_R rest vs SSp-ul_L move | **0** |
| VISp_R rest vs SSp-ul_L move | **0** |
| VISrl_L rest vs SSp-ul_L move | **0** |
| VISrl_R rest vs SSp-ul_L move | **0** |
| SSp-un_L move vs SSp-ul_R move | 0.3 |
| SSp-un_R move vs SSp-ul_R move | 1 |
| VISa_L move vs SSp-ul_R move | 0.44 |
| VISam_L move vs SSp-ul_R move | **0.004** |
| VISam_R move vs SSp-ul_R move | 0.52 |
| VISa_R move vs SSp-ul_R move | 1 |
| VISp_L move vs SSp-ul_R move | **0** |
| VISpm_L move vs SSp-ul_R move | **5.3e-09** |
| VISpm_R move vs SSp-ul_R move | **0.00097** |
| VISp_R move vs SSp-ul_R move | **1.2e-08** |
| VISrl_L move vs SSp-ul_R move | **0** |

| | |
|---|---|
| VISrl_R move vs SSp-ul_R move | **0** |
| CFA_L plan vs SSp-ul_R move | **0** |
| CFA_R plan vs SSp-ul_R move | **0** |
| MOp_L plan vs SSp-ul_R move | **0** |
| MOp_R plan vs SSp-ul_R move | **0** |
| MOs_L plan vs SSp-ul_R move | **3.5e-08** |
| MOs_R plan vs SSp-ul_R move | **0** |
| RFA_L plan vs SSp-ul_R move | **9.3e-09** |
| RFA_R plan vs SSp-ul_R move | **0** |
| RSPagl_L plan vs SSp-ul_R move | **0** |
| RSPagl_R plan vs SSp-ul_R move | **0** |
| RSPd_L plan vs SSp-ul_R move | **0** |
| RSPd_R plan vs SSp-ul_R move | **0** |
| SSp-bfd_L plan vs SSp-ul_R move | **0** |
| SSp-bfd_R plan vs SSp-ul_R move | **0** |
| SSp-ll_L plan vs SSp-ul_R move | **0** |
| SSp-ll_R plan vs SSp-ul_R move | **7.5e-09** |
| SSp-n_L plan vs SSp-ul_R move | **0** |
| SSp-n_R plan vs SSp-ul_R move | **0** |
| SSp-tr_L plan vs SSp-ul_R move | **0** |
| SSp-tr_R plan vs SSp-ul_R move | **0** |
| SSp-ul_L plan vs SSp-ul_R move | **0** |
| SSp-ul_R plan vs SSp-ul_R move | **0** |
| SSp-un_L plan vs SSp-ul_R move | **0** |
| SSp-un_R plan vs SSp-ul_R move | **0** |
| VISa_L plan vs SSp-ul_R move | **0** |
| VISam_L plan vs SSp-ul_R move | **0** |
| VISam_R plan vs SSp-ul_R move | **0** |
| VISa_R plan vs SSp-ul_R move | **0** |
| VISp_L plan vs SSp-ul_R move | **0** |
| VISpm_L plan vs SSp-ul_R move | **0** |
| VISpm_R plan vs SSp-ul_R move | **0** |
| VISp_R plan vs SSp-ul_R move | **0** |
| VISrl_L plan vs SSp-ul_R move | **0** |
| VISrl_R plan vs SSp-ul_R move | **0** |
| CFA_L rest vs SSp-ul_R move | **0** |
| CFA_R rest vs SSp-ul_R move | **0** |
| MOp_L rest vs SSp-ul_R move | **0** |
| MOp_R rest vs SSp-ul_R move | **0** |
| MOs_L rest vs SSp-ul_R move | **0** |
| MOs_R rest vs SSp-ul_R move | **0** |
| RFA_L rest vs SSp-ul_R move | **0** |
| RFA_R rest vs SSp-ul_R move | **0** |
| RSPagl_L rest vs SSp-ul_R move | **0** |
| RSPagl_R rest vs SSp-ul_R move | **0** |
| RSPd_L rest vs SSp-ul_R move | **0** |

| | |
|---|---|
| RSPd_R rest vs SSp-ul_R move | **0** |
| SSp-bfd_L rest vs SSp-ul_R move | **0** |
| SSp-bfd_R rest vs SSp-ul_R move | **0** |
| SSp-ll_L rest vs SSp-ul_R move | **0** |
| SSp-ll_R rest vs SSp-ul_R move | **0** |
| SSp-n_L rest vs SSp-ul_R move | **0** |
| SSp-n_R rest vs SSp-ul_R move | **0** |
| SSp-tr_L rest vs SSp-ul_R move | **0** |
| SSp-tr_R rest vs SSp-ul_R move | **0** |
| SSp-ul_L rest vs SSp-ul_R move | **0** |
| SSp-ul_R rest vs SSp-ul_R move | **0** |
| SSp-un_L rest vs SSp-ul_R move | **0** |
| SSp-un_R rest vs SSp-ul_R move | **0** |
| VISa_L rest vs SSp-ul_R move | **0** |
| VISam_L rest vs SSp-ul_R move | **0** |
| VISam_R rest vs SSp-ul_R move | **0** |
| VISa_R rest vs SSp-ul_R move | **0** |
| VISp_L rest vs SSp-ul_R move | **0** |
| VISpm_L rest vs SSp-ul_R move | **0** |
| VISpm_R rest vs SSp-ul_R move | **0** |
| VISp_R rest vs SSp-ul_R move | **0** |
| VISrl_L rest vs SSp-ul_R move | **0** |
| VISrl_R rest vs SSp-ul_R move | **0** |
| SSp-un_R move vs SSp-un_L move | 1 |
| VISa_L move vs SSp-un_L move | 1 |
| VISam_L move vs SSp-un_L move | 1 |
| VISam_R move vs SSp-un_L move | 1 |
| VISa_R move vs SSp-un_L move | 1 |
| VISp_L move vs SSp-un_L move | 0.011 |
| VISpm_L move vs SSp-un_L move | 0.66 |
| VISpm_R move vs SSp-un_L move | 1 |
| VISp_R move vs SSp-un_L move | 0.76 |
| VISrl_L move vs SSp-un_L move | **0.0062** |
| VISrl_R move vs SSp-un_L move | 0.34 |
| CFA_L plan vs SSp-un_L move | 0.013 |
| CFA_R plan vs SSp-un_L move | **0.0031** |
| MOp_L plan vs SSp-un_L move | **0.0016** |
| MOp_R plan vs SSp-un_L move | **0.00086** |
| MOs_L plan vs SSp-un_L move | 0.86 |
| MOs_R plan vs SSp-un_L move | 0.3 |
| RFA_L plan vs SSp-un_L move | 0.73 |
| RFA_R plan vs SSp-un_L move | 0.14 |
| RSPagl_L plan vs SSp-un_L move | **1.2e-05** |
| RSPagl_R plan vs SSp-un_L move | **6.5e-05** |
| RSPd_L plan vs SSp-un_L move | **0.0079** |
| RSPd_R plan vs SSp-un_L move | **0.0046** |

| | |
|---|---|
| SSp-bfd_L plan vs SSp-un_L move | **0** |
| SSp-bfd_R plan vs SSp-un_L move | **1.2e-09** |
| SSp-ll_L plan vs SSp-un_L move | 0.26 |
| SSp-ll_R plan vs SSp-un_L move | 0.7 |
| SSp-n_L plan vs SSp-un_L move | **0** |
| SSp-n_R plan vs SSp-un_L move | **2.9e-09** |
| SSp-tr_L plan vs SSp-un_L move | 0.021 |
| SSp-tr_R plan vs SSp-un_L move | 0.2 |
| SSp-ul_L plan vs SSp-un_L move | **0.00095** |
| SSp-ul_R plan vs SSp-un_L move | **0.005** |
| SSp-un_L plan vs SSp-un_L move | **3e-06** |
| SSp-un_R plan vs SSp-un_L move | **4.7e-05** |
| VISa_L plan vs SSp-un_L move | **2.4e-05** |
| VISam_L plan vs SSp-un_L move | **6e-07** |
| VISam_R plan vs SSp-un_L move | **5.9e-06** |
| VISa_R plan vs SSp-un_L move | **0.00018** |
| VISp_L plan vs SSp-un_L move | **0** |
| VISpm_L plan vs SSp-un_L move | **0** |
| VISpm_R plan vs SSp-un_L move | **6.2e-10** |
| VISp_R plan vs SSp-un_L move | **0** |
| VISrl_L plan vs SSp-un_L move | **0** |
| VISrl_R plan vs SSp-un_L move | **0** |
| CFA_L rest vs SSp-un_L move | **0** |
| CFA_R rest vs SSp-un_L move | **0** |
| MOp_L rest vs SSp-un_L move | **0** |
| MOp_R rest vs SSp-un_L move | **0** |
| MOs_L rest vs SSp-un_L move | **0** |
| MOs_R rest vs SSp-un_L move | **0** |
| RFA_L rest vs SSp-un_L move | **0** |
| RFA_R rest vs SSp-un_L move | **0** |
| RSPagl_L rest vs SSp-un_L move | **0** |
| RSPagl_R rest vs SSp-un_L move | **0** |
| RSPd_L rest vs SSp-un_L move | **0** |
| RSPd_R rest vs SSp-un_L move | **0** |
| SSp-bfd_L rest vs SSp-un_L move | **0** |
| SSp-bfd_R rest vs SSp-un_L move | **0** |
| SSp-ll_L rest vs SSp-un_L move | **0** |
| SSp-ll_R rest vs SSp-un_L move | **0** |
| SSp-n_L rest vs SSp-un_L move | **0** |
| SSp-n_R rest vs SSp-un_L move | **0** |
| SSp-tr_L rest vs SSp-un_L move | **0** |
| SSp-tr_R rest vs SSp-un_L move | **0** |
| SSp-ul_L rest vs SSp-un_L move | **0** |
| SSp-ul_R rest vs SSp-un_L move | **0** |
| SSp-un_L rest vs SSp-un_L move | **0** |
| SSp-un_R rest vs SSp-un_L move | **0** |

| | |
|---|---|
| VISa_L rest vs SSp-un_L move | **0** |
| VISam_L rest vs SSp-un_L move | **0** |
| VISam_R rest vs SSp-un_L move | **0** |
| VISa_R rest vs SSp-un_L move | **0** |
| VISp_L rest vs SSp-un_L move | **0** |
| VISpm_L rest vs SSp-un_L move | **0** |
| VISpm_R rest vs SSp-un_L move | **0** |
| VISp_R rest vs SSp-un_L move | **0** |
| VISrl_L rest vs SSp-un_L move | **0** |
| VISrl_R rest vs SSp-un_L move | **0** |
| VISa_L move vs SSp-un_R move | 1 |
| VISam_L move vs SSp-un_R move | 0.98 |
| VISam_R move vs SSp-un_R move | 1 |
| VISa_R move vs SSp-un_R move | 1 |
| VISp_L move vs SSp-un_R move | **1.3e-06** |
| VISpm_L move vs SSp-un_R move | **0.0025** |
| VISpm_R move vs SSp-un_R move | 0.88 |
| VISp_R move vs SSp-un_R move | **0.0041** |
| VISrl_L move vs SSp-un_R move | **5.6e-07** |
| VISrl_R move vs SSp-un_R move | **0.0004** |
| CFA_L plan vs SSp-un_R move | **1.6e-06** |
| CFA_R plan vs SSp-un_R move | **2.2e-07** |
| MOp_L plan vs SSp-un_R move | **8.3e-08** |
| MOp_R plan vs SSp-un_R move | **3.5e-08** |
| MOs_L plan vs SSp-un_R move | **0.0078** |
| MOs_R plan vs SSp-un_R move | **0.00032** |
| RFA_L plan vs SSp-un_R move | **0.0034** |
| RFA_R plan vs SSp-un_R move | **7.5e-05** |
| RSPagl_L plan vs SSp-un_R move | **0** |
| RSPagl_R plan vs SSp-un_R move | **1.8e-10** |
| RSPd_L plan vs SSp-un_R move | **8e-07** |
| RSPd_R plan vs SSp-un_R move | **3.7e-07** |
| SSp-bfd_L plan vs SSp-un_R move | **0** |
| SSp-bfd_R plan vs SSp-un_R move | **0** |
| SSp-ll_L plan vs SSp-un_R move | **0.00024** |
| SSp-ll_R plan vs SSp-un_R move | **0.003** |
| SSp-n_L plan vs SSp-un_R move | **0** |
| SSp-n_R plan vs SSp-un_R move | **0** |
| SSp-tr_L plan vs SSp-un_R move | **3.3e-06** |
| SSp-tr_R plan vs SSp-un_R move | **0.00013** |
| SSp-ul_L plan vs SSp-un_R move | **4.1e-08** |
| SSp-ul_R plan vs SSp-un_R move | **4.1e-07** |
| SSp-un_L plan vs SSp-un_R move | **0** |
| SSp-un_R plan vs SSp-un_R move | **0** |
| VISa_L plan vs SSp-un_R move | **0** |
| VISam_L plan vs SSp-un_R move | **0** |

| | |
|---|---|
| VISam_R plan vs SSp-un_R move | **0** |
| VISa_R plan vs SSp-un_R move | **3.5e-09** |
| VISp_L plan vs SSp-un_R move | **0** |
| VISpm_L plan vs SSp-un_R move | **0** |
| VISpm_R plan vs SSp-un_R move | **0** |
| VISp_R plan vs SSp-un_R move | **0** |
| VISrl_L plan vs SSp-un_R move | **0** |
| VISrl_R plan vs SSp-un_R move | **0** |
| CFA_L rest vs SSp-un_R move | **0** |
| CFA_R rest vs SSp-un_R move | **0** |
| MOp_L rest vs SSp-un_R move | **0** |
| MOp_R rest vs SSp-un_R move | **0** |
| MOs_L rest vs SSp-un_R move | **0** |
| MOs_R rest vs SSp-un_R move | **0** |
| RFA_L rest vs SSp-un_R move | **0** |
| RFA_R rest vs SSp-un_R move | **0** |
| RSPagl_L rest vs SSp-un_R move | **0** |
| RSPagl_R rest vs SSp-un_R move | **0** |
| RSPd_L rest vs SSp-un_R move | **0** |
| RSPd_R rest vs SSp-un_R move | **0** |
| SSp-bfd_L rest vs SSp-un_R move | **0** |
| SSp-bfd_R rest vs SSp-un_R move | **0** |
| SSp-ll_L rest vs SSp-un_R move | **0** |
| SSp-ll_R rest vs SSp-un_R move | **0** |
| SSp-n_L rest vs SSp-un_R move | **0** |
| SSp-n_R rest vs SSp-un_R move | **0** |
| SSp-tr_L rest vs SSp-un_R move | **0** |
| SSp-tr_R rest vs SSp-un_R move | **0** |
| SSp-ul_L rest vs SSp-un_R move | **0** |
| SSp-ul_R rest vs SSp-un_R move | **0** |
| SSp-un_L rest vs SSp-un_R move | **0** |
| SSp-un_R rest vs SSp-un_R move | **0** |
| VISa_L rest vs SSp-un_R move | **0** |
| VISam_L rest vs SSp-un_R move | **0** |
| VISam_R rest vs SSp-un_R move | **0** |
| VISa_R rest vs SSp-un_R move | **0** |
| VISp_L rest vs SSp-un_R move | **0** |
| VISpm_L rest vs SSp-un_R move | **0** |
| VISpm_R rest vs SSp-un_R move | **0** |
| VISp_R rest vs SSp-un_R move | **0** |
| VISrl_L rest vs SSp-un_R move | **0** |
| VISrl_R rest vs SSp-un_R move | **0** |
| VISam_L move vs VISa_L move | 1 |
| VISam_R move vs VISa_L move | 1 |
| VISa_R move vs VISa_L move | 1 |
| VISp_L move vs VISa_L move | **0.005** |

| | |
|---|---|
| VISpm_L move vs VISa_L move | 0.5 |
| VISpm_R move vs VISa_L move | 1 |
| VISp_R move vs VISa_L move | 0.6 |
| VISrl_L move vs VISa_L move | **0.0027** |
| VISrl_R move vs VISa_L move | 0.21 |
| CFA_L plan vs VISa_L move | **0.0058** |
| CFA_R plan vs VISa_L move | **0.0013** |
| MOp_L plan vs VISa_L move | **0.00065** |
| MOp_R plan vs VISa_L move | **0.00034** |
| MOs_L plan vs VISa_L move | 0.73 |
| MOs_R plan vs VISa_L move | 0.19 |
| RFA_L plan vs VISa_L move | 0.56 |
| RFA_R plan vs VISa_L move | 0.08 |
| RSPagl_L plan vs VISa_L move | **4.3e-06** |
| RSPagl_R plan vs VISa_L move | **2.4e-05** |
| RSPd_L plan vs VISa_L move | **0.0035** |
| RSPd_R plan vs VISa_L move | **0.002** |
| SSp-bfd_L plan vs VISa_L move | **0** |
| SSp-bfd_R plan vs VISa_L move | **0** |
| SSp-ll_L plan vs VISa_L move | 0.16 |
| SSp-ll_R plan vs VISa_L move | 0.54 |
| SSp-n_L plan vs VISa_L move | **0** |
| SSp-n_R plan vs VISa_L move | **7.1e-11** |
| SSp-tr_L plan vs VISa_L move | 0.01 |
| SSp-tr_R plan vs VISa_L move | 0.11 |
| SSp-ul_L plan vs VISa_L move | **0.00038** |
| SSp-ul_R plan vs VISa_L move | **0.0022** |
| SSp-un_L plan vs VISa_L move | **1e-06** |
| SSp-un_R plan vs VISa_L move | **1.7e-05** |
| VISa_L plan vs VISa_L move | **8.4e-06** |
| VISam_L plan vs VISa_L move | **1.9e-07** |
| VISam_R plan vs VISa_L move | **2e-06** |
| VISa_R plan vs VISa_L move | **6.7e-05** |
| VISp_L plan vs VISa_L move | **0** |
| VISpm_L plan vs VISa_L move | **0** |
| VISpm_R plan vs VISa_L move | **0** |
| VISp_R plan vs VISa_L move | **0** |
| VISrl_L plan vs VISa_L move | **0** |
| VISrl_R plan vs VISa_L move | **0** |
| CFA_L rest vs VISa_L move | **0** |
| CFA_R rest vs VISa_L move | **0** |
| MOp_L rest vs VISa_L move | **0** |
| MOp_R rest vs VISa_L move | **0** |
| MOs_L rest vs VISa_L move | **0** |
| MOs_R rest vs VISa_L move | **0** |
| RFA_L rest vs VISa_L move | **0** |

| | |
|---|---|
| RFA_R rest vs VISa_L move | **0** |
| RSPagl_L rest vs VISa_L move | **0** |
| RSPagl_R rest vs VISa_L move | **0** |
| RSPd_L rest vs VISa_L move | **0** |
| RSPd_R rest vs VISa_L move | **0** |
| SSp-bfd_L rest vs VISa_L move | **0** |
| SSp-bfd_R rest vs VISa_L move | **0** |
| SSp-ll_L rest vs VISa_L move | **0** |
| SSp-ll_R rest vs VISa_L move | **0** |
| SSp-n_L rest vs VISa_L move | **0** |
| SSp-n_R rest vs VISa_L move | **0** |
| SSp-tr_L rest vs VISa_L move | **0** |
| SSp-tr_R rest vs VISa_L move | **0** |
| SSp-ul_L rest vs VISa_L move | **0** |
| SSp-ul_R rest vs VISa_L move | **0** |
| SSp-un_L rest vs VISa_L move | **0** |
| SSp-un_R rest vs VISa_L move | **0** |
| VISa_L rest vs VISa_L move | **0** |
| VISam_L rest vs VISa_L move | **0** |
| VISam_R rest vs VISa_L move | **0** |
| VISa_R rest vs VISa_L move | **0** |
| VISp_L rest vs VISa_L move | **0** |
| VISpm_L rest vs VISa_L move | **0** |
| VISpm_R rest vs VISa_L move | **0** |
| VISp_R rest vs VISa_L move | **0** |
| VISrl_L rest vs VISa_L move | **0** |
| VISrl_R rest vs VISa_L move | **0** |
| VISam_R move vs VISam_L move | 1 |
| VISa_R move vs VISam_L move | 1 |
| VISp_L move vs VISam_L move | 0.49 |
| VISpm_L move vs VISam_L move | 1 |
| VISpm_R move vs VISam_L move | 1 |
| VISp_R move vs VISam_L move | 1 |
| VISrl_L move vs VISam_L move | 0.37 |
| VISrl_R move vs VISam_L move | 1 |
| CFA_L plan vs VISam_L move | 0.52 |
| CFA_R plan vs VISam_L move | 0.26 |
| MOp_L plan vs VISam_L move | 0.17 |
| MOp_R plan vs VISam_L move | 0.12 |
| MOs_L plan vs VISam_L move | 1 |
| MOs_R plan vs VISam_L move | 1 |
| RFA_L plan vs VISam_L move | 1 |
| RFA_R plan vs VISam_L move | 0.96 |
| RSPagl_L plan vs VISam_L move | **0.0058** |
| RSPagl_R plan vs VISam_L move | 0.02 |
| RSPd_L plan vs VISam_L move | 0.42 |

| | |
|---|---|
| RSPd_R plan vs VISam_L move | 0.32 |
| SSp-bfd_L plan vs VISam_L move | **4e-08** |
| SSp-bfd_R plan vs VISam_L move | **5.1e-06** |
| SSp-ll_L plan vs VISam_L move | 0.99 |
| SSp-ll_R plan vs VISam_L move | 1 |
| SSp-n_L plan vs VISam_L move | **8.4e-07** |
| SSp-n_R plan vs VISam_L move | **8.3e-06** |
| SSp-tr_L plan vs VISam_L move | 0.63 |
| SSp-tr_R plan vs VISam_L move | 0.98 |
| SSp-ul_L plan vs VISam_L move | 0.13 |
| SSp-ul_R plan vs VISam_L move | 0.33 |
| SSp-un_L plan vs VISam_L move | **0.0019** |
| SSp-un_R plan vs VISam_L move | 0.016 |
| VISa_L plan vs VISam_L move | **0.0095** |
| VISam_L plan vs VISam_L move | **0.00053** |
| VISam_R plan vs VISam_L move | **0.0032** |
| VISa_R plan vs VISam_L move | 0.041 |
| VISp_L plan vs VISam_L move | **0** |
| VISpm_L plan vs VISam_L move | **9e-08** |
| VISpm_R plan vs VISam_L move | **4e-06** |
| VISp_R plan vs VISam_L move | **1.3e-08** |
| VISrl_L plan vs VISam_L move | **1.9e-09** |
| VISrl_R plan vs VISam_L move | **2.1e-07** |
| CFA_L rest vs VISam_L move | **0** |
| CFA_R rest vs VISam_L move | **0** |
| MOp_L rest vs VISam_L move | **0** |
| MOp_R rest vs VISam_L move | **0** |
| MOs_L rest vs VISam_L move | **0** |
| MOs_R rest vs VISam_L move | **0** |
| RFA_L rest vs VISam_L move | **0** |
| RFA_R rest vs VISam_L move | **0** |
| RSPagl_L rest vs VISam_L move | **0** |
| RSPagl_R rest vs VISam_L move | **0** |
| RSPd_L rest vs VISam_L move | **0** |
| RSPd_R rest vs VISam_L move | **0** |
| SSp-bfd_L rest vs VISam_L move | **0** |
| SSp-bfd_R rest vs VISam_L move | **0** |
| SSp-ll_L rest vs VISam_L move | **0** |
| SSp-ll_R rest vs VISam_L move | **0** |
| SSp-n_L rest vs VISam_L move | **0** |
| SSp-n_R rest vs VISam_L move | **0** |
| SSp-tr_L rest vs VISam_L move | **0** |
| SSp-tr_R rest vs VISam_L move | **0** |
| SSp-ul_L rest vs VISam_L move | **0** |
| SSp-ul_R rest vs VISam_L move | **0** |
| SSp-un_L rest vs VISam_L move | **0** |

| | |
|---|---|
| SSp-un_R rest vs VISam_L move | **0** |
| VISa_L rest vs VISam_L move | **0** |
| VISam_L rest vs VISam_L move | **0** |
| VISam_R rest vs VISam_L move | **0** |
| VISa_R rest vs VISam_L move | **0** |
| VISp_L rest vs VISam_L move | **0** |
| VISpm_L rest vs VISam_L move | **0** |
| VISpm_R rest vs VISam_L move | **0** |
| VISp_R rest vs VISam_L move | **0** |
| VISrl_L rest vs VISam_L move | **0** |
| VISrl_R rest vs VISam_L move | **0** |
| VISa_R move vs VISam_R move | 1 |
| VISp_L move vs VISam_R move | **0.0034** |
| VISpm_L move vs VISam_R move | 0.42 |
| VISpm_R move vs VISam_R move | 1 |
| VISp_R move vs VISam_R move | 0.52 |
| VISrl_L move vs VISam_R move | **0.0018** |
| VISrl_R move vs VISam_R move | 0.17 |
| CFA_L plan vs VISam_R move | **0.004** |
| CFA_R plan vs VISam_R move | **0.00088** |
| MOp_L plan vs VISam_R move | **0.00042** |
| MOp_R plan vs VISam_R move | **0.00022** |
| MOs_L plan vs VISam_R move | 0.65 |
| MOs_R plan vs VISam_R move | 0.15 |
| RFA_L plan vs VISam_R move | 0.48 |
| RFA_R plan vs VISam_R move | 0.06 |
| RSPagl_L plan vs VISam_R move | **2.6e-06** |
| RSPagl_R plan vs VISam_R move | **1.4e-05** |
| RSPd_L plan vs VISam_R move | **0.0024** |
| RSPd_R plan vs VISam_R move | **0.0013** |
| SSp-bfd_L plan vs VISam_R move | **0** |
| SSp-bfd_R plan vs VISam_R move | **0** |
| SSp-ll_L plan vs VISam_R move | 0.12 |
| SSp-ll_R plan vs VISam_R move | 0.46 |
| SSp-n_L plan vs VISam_R move | **0** |
| SSp-n_R plan vs VISam_R move | **0** |
| SSp-tr_L plan vs VISam_R move | **0.0069** |
| SSp-tr_R plan vs VISam_R move | 0.086 |
| SSp-ul_L plan vs VISam_R move | **0.00024** |
| SSp-ul_R plan vs VISam_R move | **0.0014** |
| SSp-un_L plan vs VISam_R move | **5.9e-07** |
| SSp-un_R plan vs VISam_R move | **1e-05** |
| VISa_L plan vs VISam_R move | **5.1e-06** |
| VISam_L plan vs VISam_R move | **1.1e-07** |
| VISam_R plan vs VISam_R move | **1.2e-06** |
| VISa_R plan vs VISam_R move | **4.2e-05** |

| | |
|---|---|
| VISp_L plan vs VISam_R move | **0** |
| VISpm_L plan vs VISam_R move | **0** |
| VISpm_R plan vs VISam_R move | **0** |
| VISp_R plan vs VISam_R move | **0** |
| VISrl_L plan vs VISam_R move | **0** |
| VISrl_R plan vs VISam_R move | **0** |
| CFA_L rest vs VISam_R move | **0** |
| CFA_R rest vs VISam_R move | **0** |
| MOp_L rest vs VISam_R move | **0** |
| MOp_R rest vs VISam_R move | **0** |
| MOs_L rest vs VISam_R move | **0** |
| MOs_R rest vs VISam_R move | **0** |
| RFA_L rest vs VISam_R move | **0** |
| RFA_R rest vs VISam_R move | **0** |
| RSPagl_L rest vs VISam_R move | **0** |
| RSPagl_R rest vs VISam_R move | **0** |
| RSPd_L rest vs VISam_R move | **0** |
| RSPd_R rest vs VISam_R move | **0** |
| SSp-bfd_L rest vs VISam_R move | **0** |
| SSp-bfd_R rest vs VISam_R move | **0** |
| SSp-ll_L rest vs VISam_R move | **0** |
| SSp-ll_R rest vs VISam_R move | **0** |
| SSp-n_L rest vs VISam_R move | **0** |
| SSp-n_R rest vs VISam_R move | **0** |
| SSp-tr_L rest vs VISam_R move | **0** |
| SSp-tr_R rest vs VISam_R move | **0** |
| SSp-ul_L rest vs VISam_R move | **0** |
| SSp-ul_R rest vs VISam_R move | **0** |
| SSp-un_L rest vs VISam_R move | **0** |
| SSp-un_R rest vs VISam_R move | **0** |
| VISa_L rest vs VISam_R move | **0** |
| VISam_L rest vs VISam_R move | **0** |
| VISam_R rest vs VISam_R move | **0** |
| VISa_R rest vs VISam_R move | **0** |
| VISp_L rest vs VISam_R move | **0** |
| VISpm_L rest vs VISam_R move | **0** |
| VISpm_R rest vs VISam_R move | **0** |
| VISp_R rest vs VISam_R move | **0** |
| VISrl_L rest vs VISam_R move | **0** |
| VISrl_R rest vs VISam_R move | **0** |
| VISp_L move vs VISa_R move | **7.5e-06** |
| VISpm_L move vs VISa_R move | **0.0092** |
| VISpm_R move vs VISa_R move | 0.98 |
| VISp_R move vs VISa_R move | 0.015 |
| VISrl_L move vs VISa_R move | **3.4e-06** |
| VISrl_R move vs VISa_R move | **0.0017** |

| | |
|---|---|
| CFA_L plan vs VISa_R move | **9.2e-06** |
| CFA_R plan vs VISa_R move | **1.4e-06** |
| MOp_L plan vs VISa_R move | **5.5e-07** |
| MOp_R plan vs VISa_R move | **2.5e-07** |
| MOs_L plan vs VISa_R move | 0.026 |
| MOs_R plan vs VISa_R move | **0.0014** |
| RFA_L plan vs VISa_R move | 0.012 |
| RFA_R plan vs VISa_R move | **0.00035** |
| RSPagl_L plan vs VISa_R move | **1.8e-10** |
| RSPagl_R plan vs VISa_R move | **8.3e-09** |
| RSPd_L plan vs VISa_R move | **4.8e-06** |
| RSPd_R plan vs VISa_R move | **2.3e-06** |
| SSp-bfd_L plan vs VISa_R move | **0** |
| SSp-bfd_R plan vs VISa_R move | **0** |
| SSp-ll_L plan vs VISa_R move | **0.001** |
| SSp-ll_R plan vs VISa_R move | 0.011 |
| SSp-n_L plan vs VISa_R move | **0** |
| SSp-n_R plan vs VISa_R move | **0** |
| SSp-tr_L plan vs VISa_R move | **1.9e-05** |
| SSp-tr_R plan vs VISa_R move | **0.00061** |
| SSp-ul_L plan vs VISa_R move | **2.8e-07** |
| SSp-ul_R plan vs VISa_R move | **2.5e-06** |
| SSp-un_L plan vs VISa_R move | **0** |
| SSp-un_R plan vs VISa_R move | **5.2e-09** |
| VISa_L plan vs VISa_R move | **1.7e-09** |
| VISam_L plan vs VISa_R move | **0** |
| VISam_R plan vs VISa_R move | **0** |
| VISa_R plan vs VISa_R move | **3.2e-08** |
| VISp_L plan vs VISa_R move | **0** |
| VISpm_L plan vs VISa_R move | **0** |
| VISpm_R plan vs VISa_R move | **0** |
| VISp_R plan vs VISa_R move | **0** |
| VISrl_L plan vs VISa_R move | **0** |
| VISrl_R plan vs VISa_R move | **0** |
| CFA_L rest vs VISa_R move | **0** |
| CFA_R rest vs VISa_R move | **0** |
| MOp_L rest vs VISa_R move | **0** |
| MOp_R rest vs VISa_R move | **0** |
| MOs_L rest vs VISa_R move | **0** |
| MOs_R rest vs VISa_R move | **0** |
| RFA_L rest vs VISa_R move | **0** |
| RFA_R rest vs VISa_R move | **0** |
| RSPagl_L rest vs VISa_R move | **0** |
| RSPagl_R rest vs VISa_R move | **0** |
| RSPd_L rest vs VISa_R move | **0** |
| RSPd_R rest vs VISa_R move | **0** |

| | |
|---|---|
| SSp-bfd_L rest vs VISa_R move | **0** |
| SSp-bfd_R rest vs VISa_R move | **0** |
| SSp-ll_L rest vs VISa_R move | **0** |
| SSp-ll_R rest vs VISa_R move | **0** |
| SSp-n_L rest vs VISa_R move | **0** |
| SSp-n_R rest vs VISa_R move | **0** |
| SSp-tr_L rest vs VISa_R move | **0** |
| SSp-tr_R rest vs VISa_R move | **0** |
| SSp-ul_L rest vs VISa_R move | **0** |
| SSp-ul_R rest vs VISa_R move | **0** |
| SSp-un_L rest vs VISa_R move | **0** |
| SSp-un_R rest vs VISa_R move | **0** |
| VISa_L rest vs VISa_R move | **0** |
| VISam_L rest vs VISa_R move | **0** |
| VISam_R rest vs VISa_R move | **0** |
| VISa_R rest vs VISa_R move | **0** |
| VISp_L rest vs VISa_R move | **0** |
| VISpm_L rest vs VISa_R move | **0** |
| VISpm_R rest vs VISa_R move | **0** |
| VISp_R rest vs VISa_R move | **0** |
| VISrl_L rest vs VISa_R move | **0** |
| VISrl_R rest vs VISa_R move | **0** |
| VISpm_L move vs VISp_L move | 1 |
| VISpm_R move vs VISp_L move | 0.75 |
| VISp_R move vs VISp_L move | 1 |
| VISrl_L move vs VISp_L move | 1 |
| VISrl_R move vs VISp_L move | 1 |
| CFA_L plan vs VISp_L move | 1 |
| CFA_R plan vs VISp_L move | 1 |
| MOp_L plan vs VISp_L move | 1 |
| MOp_R plan vs VISp_L move | 1 |
| MOs_L plan vs VISp_L move | 1 |
| MOs_R plan vs VISp_L move | 1 |
| RFA_L plan vs VISp_L move | 1 |
| RFA_R plan vs VISp_L move | 1 |
| RSPagl_L plan vs VISp_L move | 1 |
| RSPagl_R plan vs VISp_L move | 1 |
| RSPd_L plan vs VISp_L move | 1 |
| RSPd_R plan vs VISp_L move | 1 |
| SSp-bfd_L plan vs VISp_L move | 0.7 |
| SSp-bfd_R plan vs VISp_L move | 1 |
| SSp-ll_L plan vs VISp_L move | 1 |
| SSp-ll_R plan vs VISp_L move | 1 |
| SSp-n_L plan vs VISp_L move | 0.97 |
| SSp-n_R plan vs VISp_L move | 1 |
| SSp-tr_L plan vs VISp_L move | 1 |

| | |
|---|---|
| SSp-tr_R plan vs VISp_L move | 1 |
| SSp-ul_L plan vs VISp_L move | 1 |
| SSp-ul_R plan vs VISp_L move | 1 |
| SSp-un_L plan vs VISp_L move | 1 |
| SSp-un_R plan vs VISp_L move | 1 |
| VISa_L plan vs VISp_L move | 1 |
| VISam_L plan vs VISp_L move | 1 |
| VISam_R plan vs VISp_L move | 1 |
| VISa_R plan vs VISp_L move | 1 |
| VISp_L plan vs VISp_L move | 0.14 |
| VISpm_L plan vs VISp_L move | 0.8 |
| VISpm_R plan vs VISp_L move | 1 |
| VISp_R plan vs VISp_L move | 0.56 |
| VISrl_L plan vs VISp_L move | 0.36 |
| VISrl_R plan vs VISp_L move | 0.88 |
| CFA_L rest vs VISp_L move | **3.1e-07** |
| CFA_R rest vs VISp_L move | **2.1e-06** |
| MOp_L rest vs VISp_L move | **7.8e-08** |
| MOp_R rest vs VISp_L move | **3.2e-07** |
| MOs_L rest vs VISp_L move | **1.4e-08** |
| MOs_R rest vs VISp_L move | **8.5e-09** |
| RFA_L rest vs VISp_L move | **7.3e-11** |
| RFA_R rest vs VISp_L move | **0** |
| RSPagl_L rest vs VISp_L move | **0** |
| RSPagl_R rest vs VISp_L move | **0** |
| RSPd_L rest vs VISp_L move | **0** |
| RSPd_R rest vs VISp_L move | **0** |
| SSp-bfd_L rest vs VISp_L move | **5.6e-08** |
| SSp-bfd_R rest vs VISp_L move | **1.2e-06** |
| SSp-ll_L rest vs VISp_L move | **7.4e-09** |
| SSp-ll_R rest vs VISp_L move | **4.9e-07** |
| SSp-n_L rest vs VISp_L move | **5.7e-07** |
| SSp-n_R rest vs VISp_L move | **2.1e-05** |
| SSp-tr_L rest vs VISp_L move | **6.4e-10** |
| SSp-tr_R rest vs VISp_L move | **3.1e-08** |
| SSp-ul_L rest vs VISp_L move | **4.8e-07** |
| SSp-ul_R rest vs VISp_L move | **2.8e-06** |
| SSp-un_L rest vs VISp_L move | **9.6e-07** |
| SSp-un_R rest vs VISp_L move | **8.7e-06** |
| VISa_L rest vs VISp_L move | **0** |
| VISam_L rest vs VISp_L move | **0** |
| VISam_R rest vs VISp_L move | **0** |
| VISa_R rest vs VISp_L move | **9.8e-10** |
| VISp_L rest vs VISp_L move | **0** |
| VISpm_L rest vs VISp_L move | **0** |
| VISpm_R rest vs VISp_L move | **0** |

| | |
|---|---|
| VISp_R rest vs VISp_L move | **0** |
| VISrl_L rest vs VISp_L move | **0** |
| VISrl_R rest vs VISp_L move | **1.3e-09** |
| VISpm_R move vs VISpm_L move | 1 |
| VISp_R move vs VISpm_L move | 1 |
| VISrl_L move vs VISpm_L move | 1 |
| VISrl_R move vs VISpm_L move | 1 |
| CFA_L plan vs VISpm_L move | 1 |
| CFA_R plan vs VISpm_L move | 1 |
| MOp_L plan vs VISpm_L move | 1 |
| MOp_R plan vs VISpm_L move | 1 |
| MOs_L plan vs VISpm_L move | 1 |
| MOs_R plan vs VISpm_L move | 1 |
| RFA_L plan vs VISpm_L move | 1 |
| RFA_R plan vs VISpm_L move | 1 |
| RSPagl_L plan vs VISpm_L move | 1 |
| RSPagl_R plan vs VISpm_L move | 1 |
| RSPd_L plan vs VISpm_L move | 1 |
| RSPd_R plan vs VISpm_L move | 1 |
| SSp-bfd_L plan vs VISpm_L move | 0.013 |
| SSp-bfd_R plan vs VISpm_L move | 0.2 |
| SSp-ll_L plan vs VISpm_L move | 1 |
| SSp-ll_R plan vs VISpm_L move | 1 |
| SSp-n_L plan vs VISpm_L move | 0.079 |
| SSp-n_R plan vs VISpm_L move | 0.25 |
| SSp-tr_L plan vs VISpm_L move | 1 |
| SSp-tr_R plan vs VISpm_L move | 1 |
| SSp-ul_L plan vs VISpm_L move | 1 |
| SSp-ul_R plan vs VISpm_L move | 1 |
| SSp-un_L plan vs VISpm_L move | 0.97 |
| SSp-un_R plan vs VISpm_L move | 1 |
| VISa_L plan vs VISpm_L move | 1 |
| VISam_L plan vs VISpm_L move | 0.87 |
| VISam_R plan vs VISpm_L move | 0.99 |
| VISa_R plan vs VISpm_L move | 1 |
| VISp_L plan vs VISpm_L move | **0.00042** |
| VISpm_L plan vs VISpm_L move | 0.022 |
| VISpm_R plan vs VISpm_L move | 0.18 |
| VISp_R plan vs VISpm_L move | **0.0068** |
| VISrl_L plan vs VISpm_L move | **0.0024** |
| VISrl_R plan vs VISpm_L move | 0.036 |
| CFA_L rest vs VISpm_L move | **0** |
| CFA_R rest vs VISpm_L move | **0** |
| MOp_L rest vs VISpm_L move | **0** |
| MOp_R rest vs VISpm_L move | **0** |
| MOs_L rest vs VISpm_L move | **0** |

| | |
|---|---|
| MOs_R rest vs VISpm_L move | **0** |
| RFA_L rest vs VISpm_L move | **0** |
| RFA_R rest vs VISpm_L move | **0** |
| RSPagl_L rest vs VISpm_L move | **0** |
| RSPagl_R rest vs VISpm_L move | **0** |
| RSPd_L rest vs VISpm_L move | **0** |
| RSPd_R rest vs VISpm_L move | **0** |
| SSp-bfd_L rest vs VISpm_L move | **0** |
| SSp-bfd_R rest vs VISpm_L move | **0** |
| SSp-ll_L rest vs VISpm_L move | **0** |
| SSp-ll_R rest vs VISpm_L move | **0** |
| SSp-n_L rest vs VISpm_L move | **0** |
| SSp-n_R rest vs VISpm_L move | **2.1e-09** |
| SSp-tr_L rest vs VISpm_L move | **0** |
| SSp-tr_R rest vs VISpm_L move | **0** |
| SSp-ul_L rest vs VISpm_L move | **0** |
| SSp-ul_R rest vs VISpm_L move | **0** |
| SSp-un_L rest vs VISpm_L move | **0** |
| SSp-un_R rest vs VISpm_L move | **3.2e-11** |
| VISa_L rest vs VISpm_L move | **0** |
| VISam_L rest vs VISpm_L move | **0** |
| VISam_R rest vs VISpm_L move | **0** |
| VISa_R rest vs VISpm_L move | **0** |
| VISp_L rest vs VISpm_L move | **0** |
| VISpm_L rest vs VISpm_L move | **0** |
| VISpm_R rest vs VISpm_L move | **0** |
| VISp_R rest vs VISpm_L move | **0** |
| VISrl_L rest vs VISpm_L move | **0** |
| VISrl_R rest vs VISpm_L move | **0** |
| VISp_R move vs VISpm_R move | 1 |
| VISrl_L move vs VISpm_R move | 0.64 |
| VISrl_R move vs VISpm_R move | 1 |
| CFA_L plan vs VISpm_R move | 0.78 |
| CFA_R plan vs VISpm_R move | 0.5 |
| MOp_L plan vs VISpm_R move | 0.37 |
| MOp_R plan vs VISpm_R move | 0.28 |
| MOs_L plan vs VISpm_R move | 1 |
| MOs_R plan vs VISpm_R move | 1 |
| RFA_L plan vs VISpm_R move | 1 |
| RFA_R plan vs VISpm_R move | 1 |
| RSPagl_L plan vs VISpm_R move | 0.02 |
| RSPagl_R plan vs VISpm_R move | 0.062 |
| RSPd_L plan vs VISpm_R move | 0.69 |
| RSPd_R plan vs VISpm_R move | 0.58 |
| SSp-bfd_L plan vs VISpm_R move | **2.9e-07** |
| SSp-bfd_R plan vs VISpm_R move | **2.9e-05** |

| | |
|---|---|
| SSp-ll_L plan vs VISpm_R move | 1 |
| SSp-ll_R plan vs VISpm_R move | 1 |
| SSp-n_L plan vs VISpm_R move | **5.2e-06** |
| SSp-n_R plan vs VISpm_R move | **4.6e-05** |
| SSp-tr_L plan vs VISpm_R move | 0.86 |
| SSp-tr_R plan vs VISpm_R move | 1 |
| SSp-ul_L plan vs VISpm_R move | 0.29 |
| SSp-ul_R plan vs VISpm_R move | 0.59 |
| SSp-un_L plan vs VISpm_R move | **0.0076** |
| SSp-un_R plan vs VISpm_R move | 0.05 |
| VISa_L plan vs VISpm_R move | 0.032 |
| VISam_L plan vs VISpm_R move | **0.0023** |
| VISam_R plan vs VISpm_R move | 0.012 |
| VISa_R plan vs VISpm_R move | 0.12 |
| VISp_L plan vs VISpm_R move | **9.4e-10** |
| VISpm_L plan vs VISpm_R move | **6.2e-07** |
| VISpm_R plan vs VISpm_R move | **2.3e-05** |
| VISp_R plan vs VISpm_R move | **1e-07** |
| VISrl_L plan vs VISpm_R move | **2.2e-08** |
| VISrl_R plan vs VISpm_R move | **1.4e-06** |
| CFA_L rest vs VISpm_R move | **0** |
| CFA_R rest vs VISpm_R move | **0** |
| MOp_L rest vs VISpm_R move | **0** |
| MOp_R rest vs VISpm_R move | **0** |
| MOs_L rest vs VISpm_R move | **0** |
| MOs_R rest vs VISpm_R move | **0** |
| RFA_L rest vs VISpm_R move | **0** |
| RFA_R rest vs VISpm_R move | **0** |
| RSPagl_L rest vs VISpm_R move | **0** |
| RSPagl_R rest vs VISpm_R move | **0** |
| RSPd_L rest vs VISpm_R move | **0** |
| RSPd_R rest vs VISpm_R move | **0** |
| SSp-bfd_L rest vs VISpm_R move | **0** |
| SSp-bfd_R rest vs VISpm_R move | **0** |
| SSp-ll_L rest vs VISpm_R move | **0** |
| SSp-ll_R rest vs VISpm_R move | **0** |
| SSp-n_L rest vs VISpm_R move | **0** |
| SSp-n_R rest vs VISpm_R move | **0** |
| SSp-tr_L rest vs VISpm_R move | **0** |
| SSp-tr_R rest vs VISpm_R move | **0** |
| SSp-ul_L rest vs VISpm_R move | **0** |
| SSp-ul_R rest vs VISpm_R move | **0** |
| SSp-un_L rest vs VISpm_R move | **0** |
| SSp-un_R rest vs VISpm_R move | **0** |
| VISa_L rest vs VISpm_R move | **0** |
| VISam_L rest vs VISpm_R move | **0** |

| | |
|---|---|
| VISam_R rest vs VISpm_R move | **0** |
| VISa_R rest vs VISpm_R move | **0** |
| VISp_L rest vs VISpm_R move | **0** |
| VISpm_L rest vs VISpm_R move | **0** |
| VISpm_R rest vs VISpm_R move | **0** |
| VISp_R rest vs VISpm_R move | **0** |
| VISrl_L rest vs VISpm_R move | **0** |
| VISrl_R rest vs VISpm_R move | **0** |
| VISrl_L move vs VISp_R move | 1 |
| VISrl_R move vs VISp_R move | 1 |
| CFA_L plan vs VISp_R move | 1 |
| CFA_R plan vs VISp_R move | 1 |
| MOp_L plan vs VISp_R move | 1 |
| MOp_R plan vs VISp_R move | 1 |
| MOs_L plan vs VISp_R move | 1 |
| MOs_R plan vs VISp_R move | 1 |
| RFA_L plan vs VISp_R move | 1 |
| RFA_R plan vs VISp_R move | 1 |
| RSPagl_L plan vs VISp_R move | 0.99 |
| RSPagl_R plan vs VISp_R move | 1 |
| RSPd_L plan vs VISp_R move | 1 |
| RSPd_R plan vs VISp_R move | 1 |
| SSp-bfd_L plan vs VISp_R move | **0.0084** |
| SSp-bfd_R plan vs VISp_R move | 0.14 |
| SSp-ll_L plan vs VISp_R move | 1 |
| SSp-ll_R plan vs VISp_R move | 1 |
| SSp-n_L plan vs VISp_R move | 0.054 |
| SSp-n_R plan vs VISp_R move | 0.18 |
| SSp-tr_L plan vs VISp_R move | 1 |
| SSp-tr_R plan vs VISp_R move | 1 |
| SSp-ul_L plan vs VISp_R move | 1 |
| SSp-ul_R plan vs VISp_R move | 1 |
| SSp-un_L plan vs VISp_R move | 0.94 |
| SSp-un_R plan vs VISp_R move | 1 |
| VISa_L plan vs VISp_R move | 1 |
| VISam_L plan vs VISp_R move | 0.79 |
| VISam_R plan vs VISp_R move | 0.97 |
| VISa_R plan vs VISp_R move | 1 |
| VISp_L plan vs VISp_R move | **0.00024** |
| VISpm_L plan vs VISp_R move | 0.014 |
| VISpm_R plan vs VISp_R move | 0.13 |
| VISp_R plan vs VISp_R move | **0.0042** |
| VISrl_L plan vs VISp_R move | **0.0014** |
| VISrl_R plan vs VISp_R move | 0.023 |
| CFA_L rest vs VISp_R move | **0** |
| CFA_R rest vs VISp_R move | **0** |

| | |
|---|---|
| MOp_L rest vs VISp_R move | **0** |
| MOp_R rest vs VISp_R move | **0** |
| MOs_L rest vs VISp_R move | **0** |
| MOs_R rest vs VISp_R move | **0** |
| RFA_L rest vs VISp_R move | **0** |
| RFA_R rest vs VISp_R move | **0** |
| RSPagl_L rest vs VISp_R move | **0** |
| RSPagl_R rest vs VISp_R move | **0** |
| RSPd_L rest vs VISp_R move | **0** |
| RSPd_R rest vs VISp_R move | **0** |
| SSp-bfd_L rest vs VISp_R move | **0** |
| SSp-bfd_R rest vs VISp_R move | **0** |
| SSp-ll_L rest vs VISp_R move | **0** |
| SSp-ll_R rest vs VISp_R move | **0** |
| SSp-n_L rest vs VISp_R move | **0** |
| SSp-n_R rest vs VISp_R move | **4e-10** |
| SSp-tr_L rest vs VISp_R move | **0** |
| SSp-tr_R rest vs VISp_R move | **0** |
| SSp-ul_L rest vs VISp_R move | **0** |
| SSp-ul_R rest vs VISp_R move | **0** |
| SSp-un_L rest vs VISp_R move | **0** |
| SSp-un_R rest vs VISp_R move | **0** |
| VISa_L rest vs VISp_R move | **0** |
| VISam_L rest vs VISp_R move | **0** |
| VISam_R rest vs VISp_R move | **0** |
| VISa_R rest vs VISp_R move | **0** |
| VISp_L rest vs VISp_R move | **0** |
| VISpm_L rest vs VISp_R move | **0** |
| VISpm_R rest vs VISp_R move | **0** |
| VISp_R rest vs VISp_R move | **0** |
| VISrl_L rest vs VISp_R move | **0** |
| VISrl_R rest vs VISp_R move | **0** |
| VISrl_R move vs VISrl_L move | 1 |
| CFA_L plan vs VISrl_L move | 1 |
| CFA_R plan vs VISrl_L move | 1 |
| MOp_L plan vs VISrl_L move | 1 |
| MOp_R plan vs VISrl_L move | 1 |
| MOs_L plan vs VISrl_L move | 1 |
| MOs_R plan vs VISrl_L move | 1 |
| RFA_L plan vs VISrl_L move | 1 |
| RFA_R plan vs VISrl_L move | 1 |
| RSPagl_L plan vs VISrl_L move | 1 |
| RSPagl_R plan vs VISrl_L move | 1 |
| RSPd_L plan vs VISrl_L move | 1 |
| RSPd_R plan vs VISrl_L move | 1 |
| SSp-bfd_L plan vs VISrl_L move | 0.81 |

| | |
|---|---|
| SSp-bfd_R plan vs VISrl_L move | 1 |
| SSp-ll_L plan vs VISrl_L move | 1 |
| SSp-ll_R plan vs VISrl_L move | 1 |
| SSp-n_L plan vs VISrl_L move | 0.99 |
| SSp-n_R plan vs VISrl_L move | 1 |
| SSp-tr_L plan vs VISrl_L move | 1 |
| SSp-tr_R plan vs VISrl_L move | 1 |
| SSp-ul_L plan vs VISrl_L move | 1 |
| SSp-ul_R plan vs VISrl_L move | 1 |
| SSp-un_L plan vs VISrl_L move | 1 |
| SSp-un_R plan vs VISrl_L move | 1 |
| VISa_L plan vs VISrl_L move | 1 |
| VISam_L plan vs VISrl_L move | 1 |
| VISam_R plan vs VISrl_L move | 1 |
| VISa_R plan vs VISrl_L move | 1 |
| VISp_L plan vs VISrl_L move | 0.2 |
| VISpm_L plan vs VISrl_L move | 0.88 |
| VISpm_R plan vs VISrl_L move | 1 |
| VISp_R plan vs VISrl_L move | 0.68 |
| VISrl_L plan vs VISrl_L move | 0.47 |
| VISrl_R plan vs VISrl_L move | 0.94 |
| CFA_L rest vs VISrl_L move | **7.2e-07** |
| CFA_R rest vs VISrl_L move | **4.8e-06** |
| MOp_L rest vs VISrl_L move | **1.9e-07** |
| MOp_R rest vs VISrl_L move | **7.5e-07** |
| MOs_L rest vs VISrl_L move | **3.7e-08** |
| MOs_R rest vs VISrl_L move | **2.2e-08** |
| RFA_L rest vs VISrl_L move | **1.8e-09** |
| RFA_R rest vs VISrl_L move | **0** |
| RSPagl_L rest vs VISrl_L move | **0** |
| RSPagl_R rest vs VISrl_L move | **0** |
| RSPd_L rest vs VISrl_L move | **0** |
| RSPd_R rest vs VISrl_L move | **0** |
| SSp-bfd_L rest vs VISrl_L move | **1.4e-07** |
| SSp-bfd_R rest vs VISrl_L move | **2.8e-06** |
| SSp-ll_L rest vs VISrl_L move | **2e-08** |
| SSp-ll_R rest vs VISrl_L move | **1.1e-06** |
| SSp-n_L rest vs VISrl_L move | **1.3e-06** |
| SSp-n_R rest vs VISrl_L move | **4.4e-05** |
| SSp-tr_L rest vs VISrl_L move | **3.2e-09** |
| SSp-tr_R rest vs VISrl_L move | **7.6e-08** |
| SSp-ul_L rest vs VISrl_L move | **1.1e-06** |
| SSp-ul_R rest vs VISrl_L move | **6.2e-06** |
| SSp-un_L rest vs VISrl_L move | **2.2e-06** |
| SSp-un_R rest vs VISrl_L move | **1.9e-05** |
| VISa_L rest vs VISrl_L move | **1.2e-09** |

| | |
|---|---|
| VISam_L rest vs VISrl_L move | **0** |
| VISam_R rest vs VISrl_L move | **0** |
| VISa_R rest vs VISrl_L move | **4.1e-09** |
| VISp_L rest vs VISrl_L move | **0** |
| VISpm_L rest vs VISrl_L move | **0** |
| VISpm_R rest vs VISrl_L move | **0** |
| VISp_R rest vs VISrl_L move | **0** |
| VISrl_L rest vs VISrl_L move | **2.7e-10** |
| VISrl_R rest vs VISrl_L move | **5e-09** |
| CFA_L plan vs VISrl_R move | 1 |
| CFA_R plan vs VISrl_R move | 1 |
| MOp_L plan vs VISrl_R move | 1 |
| MOp_R plan vs VISrl_R move | 1 |
| MOs_L plan vs VISrl_R move | 1 |
| MOs_R plan vs VISrl_R move | 1 |
| RFA_L plan vs VISrl_R move | 1 |
| RFA_R plan vs VISrl_R move | 1 |
| RSPagl_L plan vs VISrl_R move | 1 |
| RSPagl_R plan vs VISrl_R move | 1 |
| RSPd_L plan vs VISrl_R move | 1 |
| RSPd_R plan vs VISrl_R move | 1 |
| SSp-bfd_L plan vs VISrl_R move | 0.056 |
| SSp-bfd_R plan vs VISrl_R move | 0.48 |
| SSp-ll_L plan vs VISrl_R move | 1 |
| SSp-ll_R plan vs VISrl_R move | 1 |
| SSp-n_L plan vs VISrl_R move | 0.24 |
| SSp-n_R plan vs VISrl_R move | 0.55 |
| SSp-tr_L plan vs VISrl_R move | 1 |
| SSp-tr_R plan vs VISrl_R move | 1 |
| SSp-ul_L plan vs VISrl_R move | 1 |
| SSp-ul_R plan vs VISrl_R move | 1 |
| SSp-un_L plan vs VISrl_R move | 1 |
| SSp-un_R plan vs VISrl_R move | 1 |
| VISa_L plan vs VISrl_R move | 1 |
| VISam_L plan vs VISrl_R move | 0.99 |
| VISam_R plan vs VISrl_R move | 1 |
| VISa_R plan vs VISrl_R move | 1 |
| VISp_L plan vs VISrl_R move | **0.0026** |
| VISpm_L plan vs VISrl_R move | 0.085 |
| VISpm_R plan vs VISrl_R move | 0.44 |
| VISp_R plan vs VISrl_R move | 0.031 |
| VISrl_L plan vs VISrl_R move | 0.012 |
| VISrl_R plan vs VISrl_R move | 0.13 |
| CFA_L rest vs VISrl_R move | **0** |
| CFA_R rest vs VISrl_R move | **1.8e-09** |
| MOp_L rest vs VISrl_R move | **0** |

| | |
|---|---|
| MOp_R rest vs VISrl_R move | **0** |
| MOs_L rest vs VISrl_R move | **0** |
| MOs_R rest vs VISrl_R move | **0** |
| RFA_L rest vs VISrl_R move | **0** |
| RFA_R rest vs VISrl_R move | **0** |
| RSPagl_L rest vs VISrl_R move | **0** |
| RSPagl_R rest vs VISrl_R move | **0** |
| RSPd_L rest vs VISrl_R move | **0** |
| RSPd_R rest vs VISrl_R move | **0** |
| SSp-bfd_L rest vs VISrl_R move | **0** |
| SSp-bfd_R rest vs VISrl_R move | **4.4e-10** |
| SSp-ll_L rest vs VISrl_R move | **0** |
| SSp-ll_R rest vs VISrl_R move | **0** |
| SSp-n_L rest vs VISrl_R move | **0** |
| SSp-n_R rest vs VISrl_R move | **3.8e-08** |
| SSp-tr_L rest vs VISrl_R move | **0** |
| SSp-tr_R rest vs VISrl_R move | **0** |
| SSp-ul_L rest vs VISrl_R move | **0** |
| SSp-ul_R rest vs VISrl_R move | **2.8e-09** |
| SSp-un_L rest vs VISrl_R move | **7.8e-11** |
| SSp-un_R rest vs VISrl_R move | **1.3e-08** |
| VISa_L rest vs VISrl_R move | **0** |
| VISam_L rest vs VISrl_R move | **0** |
| VISam_R rest vs VISrl_R move | **0** |
| VISa_R rest vs VISrl_R move | **0** |
| VISp_L rest vs VISrl_R move | **0** |
| VISpm_L rest vs VISrl_R move | **0** |
| VISpm_R rest vs VISrl_R move | **0** |
| VISp_R rest vs VISrl_R move | **0** |
| VISrl_L rest vs VISrl_R move | **0** |
| VISrl_R rest vs VISrl_R move | **0** |
| CFA_R plan vs CFA_L plan | 1 |
| MOp_L plan vs CFA_L plan | 1 |
| MOp_R plan vs CFA_L plan | 1 |
| MOs_L plan vs CFA_L plan | 1 |
| MOs_R plan vs CFA_L plan | 1 |
| RFA_L plan vs CFA_L plan | 1 |
| RFA_R plan vs CFA_L plan | 1 |
| RSPagl_L plan vs CFA_L plan | 1 |
| RSPagl_R plan vs CFA_L plan | 1 |
| RSPd_L plan vs CFA_L plan | 1 |
| RSPd_R plan vs CFA_L plan | 1 |
| SSp-bfd_L plan vs CFA_L plan | 0.67 |
| SSp-bfd_R plan vs CFA_L plan | 1 |
| SSp-ll_L plan vs CFA_L plan | 1 |
| SSp-ll_R plan vs CFA_L plan | 1 |

| | |
|---|---|
| SSp-n_L plan vs CFA_L plan | 0.96 |
| SSp-n_R plan vs CFA_L plan | 1 |
| SSp-tr_L plan vs CFA_L plan | 1 |
| SSp-tr_R plan vs CFA_L plan | 1 |
| SSp-ul_L plan vs CFA_L plan | 1 |
| SSp-ul_R plan vs CFA_L plan | 1 |
| SSp-un_L plan vs CFA_L plan | 1 |
| SSp-un_R plan vs CFA_L plan | 1 |
| VISa_L plan vs CFA_L plan | 1 |
| VISam_L plan vs CFA_L plan | 1 |
| VISam_R plan vs CFA_L plan | 1 |
| VISa_R plan vs CFA_L plan | 1 |
| VISp_L plan vs CFA_L plan | 0.12 |
| VISpm_L plan vs CFA_L plan | 0.77 |
| VISpm_R plan vs CFA_L plan | 0.99 |
| VISp_R plan vs CFA_L plan | 0.53 |
| VISrl_L plan vs CFA_L plan | 0.33 |
| VISrl_R plan vs CFA_L plan | 0.86 |
| CFA_L rest vs CFA_L plan | **2.5e-07** |
| CFA_R rest vs CFA_L plan | **1.7e-06** |
| MOp_L rest vs CFA_L plan | **6.2e-08** |
| MOp_R rest vs CFA_L plan | **2.6e-07** |
| MOs_L rest vs CFA_L plan | **1.1e-08** |
| MOs_R rest vs CFA_L plan | **6.5e-09** |
| RFA_L rest vs CFA_L plan | **0** |
| RFA_R rest vs CFA_L plan | **0** |
| RSPagl_L rest vs CFA_L plan | **0** |
| RSPagl_R rest vs CFA_L plan | **0** |
| RSPd_L rest vs CFA_L plan | **0** |
| RSPd_R rest vs CFA_L plan | **0** |
| SSp-bfd_L rest vs CFA_L plan | **4.4e-08** |
| SSp-bfd_R rest vs CFA_L plan | **1e-06** |
| SSp-ll_L rest vs CFA_L plan | **5.6e-09** |
| SSp-ll_R rest vs CFA_L plan | **3.9e-07** |
| SSp-n_L rest vs CFA_L plan | **4.6e-07** |
| SSp-n_R rest vs CFA_L plan | **1.7e-05** |
| SSp-tr_L rest vs CFA_L plan | **2.8e-10** |
| SSp-tr_R rest vs CFA_L plan | **2.4e-08** |
| SSp-ul_L rest vs CFA_L plan | **3.9e-07** |
| SSp-ul_R rest vs CFA_L plan | **2.3e-06** |
| SSp-un_L rest vs CFA_L plan | **7.8e-07** |
| SSp-un_R rest vs CFA_L plan | **7.1e-06** |
| VISa_L rest vs CFA_L plan | **0** |
| VISam_L rest vs CFA_L plan | **0** |
| VISam_R rest vs CFA_L plan | **0** |
| VISa_R rest vs CFA_L plan | **5.5e-10** |

| | |
|---|---|
| VISp_L rest vs CFA_L plan | **0** |
| VISpm_L rest vs CFA_L plan | **0** |
| VISpm_R rest vs CFA_L plan | **0** |
| VISp_R rest vs CFA_L plan | **0** |
| VISrl_L rest vs CFA_L plan | **0** |
| VISrl_R rest vs CFA_L plan | **8.3e-10** |
| MOp_L plan vs CFA_R plan | 1 |
| MOp_R plan vs CFA_R plan | 1 |
| MOs_L plan vs CFA_R plan | 1 |
| MOs_R plan vs CFA_R plan | 1 |
| RFA_L plan vs CFA_R plan | 1 |
| RFA_R plan vs CFA_R plan | 1 |
| RSPagl_L plan vs CFA_R plan | 1 |
| RSPagl_R plan vs CFA_R plan | 1 |
| RSPd_L plan vs CFA_R plan | 1 |
| RSPd_R plan vs CFA_R plan | 1 |
| SSp-bfd_L plan vs CFA_R plan | 0.9 |
| SSp-bfd_R plan vs CFA_R plan | 1 |
| SSp-ll_L plan vs CFA_R plan | 1 |
| SSp-ll_R plan vs CFA_R plan | 1 |
| SSp-n_L plan vs CFA_R plan | 1 |
| SSp-n_R plan vs CFA_R plan | 1 |
| SSp-tr_L plan vs CFA_R plan | 1 |
| SSp-tr_R plan vs CFA_R plan | 1 |
| SSp-ul_L plan vs CFA_R plan | 1 |
| SSp-ul_R plan vs CFA_R plan | 1 |
| SSp-un_L plan vs CFA_R plan | 1 |
| SSp-un_R plan vs CFA_R plan | 1 |
| VISa_L plan vs CFA_R plan | 1 |
| VISam_L plan vs CFA_R plan | 1 |
| VISam_R plan vs CFA_R plan | 1 |
| VISa_R plan vs CFA_R plan | 1 |
| VISp_L plan vs CFA_R plan | 0.3 |
| VISpm_L plan vs CFA_R plan | 0.95 |
| VISpm_R plan vs CFA_R plan | 1 |
| VISp_R plan vs CFA_R plan | 0.8 |
| VISrl_L plan vs CFA_R plan | 0.61 |
| VISrl_R plan vs CFA_R plan | 0.98 |
| CFA_L rest vs CFA_R plan | **1.8e-06** |
| CFA_R rest vs CFA_R plan | **1.2e-05** |
| MOp_L rest vs CFA_R plan | **4.9e-07** |
| MOp_R rest vs CFA_R plan | **1.9e-06** |
| MOs_L rest vs CFA_R plan | **1e-07** |
| MOs_R rest vs CFA_R plan | **6.3e-08** |
| RFA_L rest vs CFA_R plan | **7e-09** |
| RFA_R rest vs CFA_R plan | **1.1e-09** |

| | |
|---|---|
| RSPagl_L rest vs CFA_R plan | **0** |
| RSPagl_R rest vs CFA_R plan | **0** |
| RSPd_L rest vs CFA_R plan | **0** |
| RSPd_R rest vs CFA_R plan | **0** |
| SSp-bfd_L rest vs CFA_R plan | **3.6e-07** |
| SSp-bfd_R rest vs CFA_R plan | **6.8e-06** |
| SSp-ll_L rest vs CFA_R plan | **5.6e-08** |
| SSp-ll_R rest vs CFA_R plan | **2.8e-06** |
| SSp-n_L rest vs CFA_R plan | **3.3e-06** |
| SSp-n_R rest vs CFA_R plan | **0.0001** |
| SSp-tr_L rest vs CFA_R plan | **1.1e-08** |
| SSp-tr_R rest vs CFA_R plan | **2e-07** |
| SSp-ul_L rest vs CFA_R plan | **2.8e-06** |
| SSp-ul_R rest vs CFA_R plan | **1.5e-05** |
| SSp-un_L rest vs CFA_R plan | **5.4e-06** |
| SSp-un_R rest vs CFA_R plan | **4.4e-05** |
| VISa_L rest vs CFA_R plan | **5.3e-09** |
| VISam_L rest vs CFA_R plan | **0** |
| VISam_R rest vs CFA_R plan | **6.5e-10** |
| VISa_R rest vs CFA_R plan | **1.3e-08** |
| VISp_L rest vs CFA_R plan | **0** |
| VISpm_L rest vs CFA_R plan | **0** |
| VISpm_R rest vs CFA_R plan | **0** |
| VISp_R rest vs CFA_R plan | **0** |
| VISrl_L rest vs CFA_R plan | **2.7e-09** |
| VISrl_R rest vs CFA_R plan | **1.6e-08** |
| MOp_R plan vs MOp_L plan | 1 |
| MOs_L plan vs MOp_L plan | 1 |
| MOs_R plan vs MOp_L plan | 1 |
| RFA_L plan vs MOp_L plan | 1 |
| RFA_R plan vs MOp_L plan | 1 |
| RSPagl_L plan vs MOp_L plan | 1 |
| RSPagl_R plan vs MOp_L plan | 1 |
| RSPd_L plan vs MOp_L plan | 1 |
| RSPd_R plan vs MOp_L plan | 1 |
| SSp-bfd_L plan vs MOp_L plan | 0.96 |
| SSp-bfd_R plan vs MOp_L plan | 1 |
| SSp-ll_L plan vs MOp_L plan | 1 |
| SSp-ll_R plan vs MOp_L plan | 1 |
| SSp-n_L plan vs MOp_L plan | 1 |
| SSp-n_R plan vs MOp_L plan | 1 |
| SSp-tr_L plan vs MOp_L plan | 1 |
| SSp-tr_R plan vs MOp_L plan | 1 |
| SSp-ul_L plan vs MOp_L plan | 1 |
| SSp-ul_R plan vs MOp_L plan | 1 |
| SSp-un_L plan vs MOp_L plan | 1 |

| | |
|---|---|
| SSp-un_R plan vs MOp_L plan | 1 |
| VISa_L plan vs MOp_L plan | 1 |
| VISam_L plan vs MOp_L plan | 1 |
| VISam_R plan vs MOp_L plan | 1 |
| VISa_R plan vs MOp_L plan | 1 |
| VISp_L plan vs MOp_L plan | 0.42 |
| VISpm_L plan vs MOp_L plan | 0.98 |
| VISpm_R plan vs MOp_L plan | 1 |
| VISp_R plan vs MOp_L plan | 0.89 |
| VISrl_L plan vs MOp_L plan | 0.74 |
| VISrl_R plan vs MOp_L plan | 0.99 |
| CFA_L rest vs MOp_L plan | **4.4e-06** |
| CFA_R rest vs MOp_L plan | **2.7e-05** |
| MOp_L rest vs MOp_L plan | **1.2e-06** |
| MOp_R rest vs MOp_L plan | **4.6e-06** |
| MOs_L rest vs MOp_L plan | **2.6e-07** |
| MOs_R rest vs MOp_L plan | **1.6e-07** |
| RFA_L rest vs MOp_L plan | **2.1e-08** |
| RFA_R rest vs MOp_L plan | **4.8e-09** |
| RSPagl_L rest vs MOp_L plan | **0** |
| RSPagl_R rest vs MOp_L plan | **8.3e-10** |
| RSPd_L rest vs MOp_L plan | **0** |
| RSPd_R rest vs MOp_L plan | **4.5e-10** |
| SSp-bfd_L rest vs MOp_L plan | **8.9e-07** |
| SSp-bfd_R rest vs MOp_L plan | **1.6e-05** |
| SSp-ll_L rest vs MOp_L plan | **1.5e-07** |
| SSp-ll_R rest vs MOp_L plan | **6.7e-06** |
| SSp-n_L rest vs MOp_L plan | **7.7e-06** |
| SSp-n_R rest vs MOp_L plan | **0.00022** |
| SSp-tr_L rest vs MOp_L plan | **3.1e-08** |
| SSp-tr_R rest vs MOp_L plan | **5.1e-07** |
| SSp-ul_L rest vs MOp_L plan | **6.6e-06** |
| SSp-ul_R rest vs MOp_L plan | **3.4e-05** |
| SSp-un_L rest vs MOp_L plan | **1.3e-05** |
| SSp-un_R rest vs MOp_L plan | **9.7e-05** |
| VISa_L rest vs MOp_L plan | **1.6e-08** |
| VISam_L rest vs MOp_L plan | **7.5e-10** |
| VISam_R rest vs MOp_L plan | **3.6e-09** |
| VISa_R rest vs MOp_L plan | **3.7e-08** |
| VISp_L rest vs MOp_L plan | **0** |
| VISpm_L rest vs MOp_L plan | **0** |
| VISpm_R rest vs MOp_L plan | **0** |
| VISp_R rest vs MOp_L plan | **2e-10** |
| VISrl_L rest vs MOp_L plan | **9.2e-09** |
| VISrl_R rest vs MOp_L plan | **4.3e-08** |
| MOs_L plan vs MOp_R plan | 1 |

| | |
|---|---|
| MOs_R plan vs MOp_R plan | 1 |
| RFA_L plan vs MOp_R plan | 1 |
| RFA_R plan vs MOp_R plan | 1 |
| RSPagl_L plan vs MOp_R plan | 1 |
| RSPagl_R plan vs MOp_R plan | 1 |
| RSPd_L plan vs MOp_R plan | 1 |
| RSPd_R plan vs MOp_R plan | 1 |
| SSp-bfd_L plan vs MOp_R plan | 0.98 |
| SSp-bfd_R plan vs MOp_R plan | 1 |
| SSp-ll_L plan vs MOp_R plan | 1 |
| SSp-ll_R plan vs MOp_R plan | 1 |
| SSp-n_L plan vs MOp_R plan | 1 |
| SSp-n_R plan vs MOp_R plan | 1 |
| SSp-tr_L plan vs MOp_R plan | 1 |
| SSp-tr_R plan vs MOp_R plan | 1 |
| SSp-ul_L plan vs MOp_R plan | 1 |
| SSp-ul_R plan vs MOp_R plan | 1 |
| SSp-un_L plan vs MOp_R plan | 1 |
| SSp-un_R plan vs MOp_R plan | 1 |
| VISa_L plan vs MOp_R plan | 1 |
| VISam_L plan vs MOp_R plan | 1 |
| VISam_R plan vs MOp_R plan | 1 |
| VISa_R plan vs MOp_R plan | 1 |
| VISp_L plan vs MOp_R plan | 0.53 |
| VISpm_L plan vs MOp_R plan | 0.99 |
| VISpm_R plan vs MOp_R plan | 1 |
| VISp_R plan vs MOp_R plan | 0.95 |
| VISrl_L plan vs MOp_R plan | 0.83 |
| VISrl_R plan vs MOp_R plan | 1 |
| CFA_L rest vs MOp_R plan | **9.3e-06** |
| CFA_R rest vs MOp_R plan | **5.4e-05** |
| MOp_L rest vs MOp_R plan | **2.6e-06** |
| MOp_R rest vs MOp_R plan | **9.7e-06** |
| MOs_L rest vs MOp_R plan | **5.8e-07** |
| MOs_R rest vs MOp_R plan | **3.7e-07** |
| RFA_L rest vs MOp_R plan | **4.9e-08** |
| RFA_R rest vs MOp_R plan | **1.3e-08** |
| RSPagl_L rest vs MOp_R plan | **2.8e-10** |
| RSPagl_R rest vs MOp_R plan | **3.5e-09** |
| RSPd_L rest vs MOp_R plan | **1.3e-09** |
| RSPd_R rest vs MOp_R plan | **2.6e-09** |
| SSp-bfd_L rest vs MOp_R plan | **1.9e-06** |
| SSp-bfd_R rest vs MOp_R plan | **3.3e-05** |
| SSp-ll_L rest vs MOp_R plan | **3.3e-07** |
| SSp-ll_R rest vs MOp_R plan | **1.4e-05** |
| SSp-n_L rest vs MOp_R plan | **1.6e-05** |

| | |
|---|---|
| SSp-n_R rest vs MOp_R plan | **0.00042** |
| SSp-tr_L rest vs MOp_R plan | **7.3e-08** |
| SSp-tr_R rest vs MOp_R plan | **1.1e-06** |
| SSp-ul_L rest vs MOp_R plan | **1.4e-05** |
| SSp-ul_R rest vs MOp_R plan | **6.9e-05** |
| SSp-un_L rest vs MOp_R plan | **2.6e-05** |
| SSp-un_R rest vs MOp_R plan | **0.00019** |
| VISa_L rest vs MOp_R plan | **3.9e-08** |
| VISam_L rest vs MOp_R plan | **3.3e-09** |
| VISam_R rest vs MOp_R plan | **1e-08** |
| VISa_R rest vs MOp_R plan | **8.7e-08** |
| VISp_L rest vs MOp_R plan | **0** |
| VISpm_L rest vs MOp_R plan | **0** |
| VISpm_R rest vs MOp_R plan | **0** |
| VISp_R rest vs MOp_R plan | **2e-09** |
| VISrl_L rest vs MOp_R plan | **2.3e-08** |
| VISrl_R rest vs MOp_R plan | **1e-07** |
| MOs_R plan vs MOs_L plan | 1 |
| RFA_L plan vs MOs_L plan | 1 |
| RFA_R plan vs MOs_L plan | 1 |
| RSPagl_L plan vs MOs_L plan | 0.97 |
| RSPagl_R plan vs MOs_L plan | 1 |
| RSPd_L plan vs MOs_L plan | 1 |
| RSPd_R plan vs MOs_L plan | 1 |
| SSp-bfd_L plan vs MOs_L plan | **0.0044** |
| SSp-bfd_R plan vs MOs_L plan | 0.09 |
| SSp-ll_L plan vs MOs_L plan | 1 |
| SSp-ll_R plan vs MOs_L plan | 1 |
| SSp-n_L plan vs MOs_L plan | 0.031 |
| SSp-n_R plan vs MOs_L plan | 0.12 |
| SSp-tr_L plan vs MOs_L plan | 1 |
| SSp-tr_R plan vs MOs_L plan | 1 |
| SSp-ul_L plan vs MOs_L plan | 1 |
| SSp-ul_R plan vs MOs_L plan | 1 |
| SSp-un_L plan vs MOs_L plan | 0.87 |
| SSp-un_R plan vs MOs_L plan | 0.99 |
| VISa_L plan vs MOs_L plan | 0.98 |
| VISam_L plan vs MOs_L plan | 0.67 |
| VISam_R plan vs MOs_L plan | 0.93 |
| VISa_R plan vs MOs_L plan | 1 |
| VISp_L plan vs MOs_L plan | **0.00011** |
| VISpm_L plan vs MOs_L plan | **0.0075** |
| VISpm_R plan vs MOs_L plan | 0.079 |
| VISp_R plan vs MOs_L plan | **0.0021** |
| VISrl_L plan vs MOs_L plan | **0.0007** |
| VISrl_R plan vs MOs_L plan | 0.013 |

| | |
|---|---|
| CFA_L rest vs MOs_L plan | **0** |
| CFA_R rest vs MOs_L plan | **0** |
| MOp_L rest vs MOs_L plan | **0** |
| MOp_R rest vs MOs_L plan | **0** |
| MOs_L rest vs MOs_L plan | **0** |
| MOs_R rest vs MOs_L plan | **0** |
| RFA_L rest vs MOs_L plan | **0** |
| RFA_R rest vs MOs_L plan | **0** |
| RSPagl_L rest vs MOs_L plan | **0** |
| RSPagl_R rest vs MOs_L plan | **0** |
| RSPd_L rest vs MOs_L plan | **0** |
| RSPd_R rest vs MOs_L plan | **0** |
| SSp-bfd_L rest vs MOs_L plan | **0** |
| SSp-bfd_R rest vs MOs_L plan | **0** |
| SSp-ll_L rest vs MOs_L plan | **0** |
| SSp-ll_R rest vs MOs_L plan | **0** |
| SSp-n_L rest vs MOs_L plan | **0** |
| SSp-n_R rest vs MOs_L plan | **0** |
| SSp-tr_L rest vs MOs_L plan | **0** |
| SSp-tr_R rest vs MOs_L plan | **0** |
| SSp-ul_L rest vs MOs_L plan | **0** |
| SSp-ul_R rest vs MOs_L plan | **0** |
| SSp-un_L rest vs MOs_L plan | **0** |
| SSp-un_R rest vs MOs_L plan | **0** |
| VISa_L rest vs MOs_L plan | **0** |
| VISam_L rest vs MOs_L plan | **0** |
| VISam_R rest vs MOs_L plan | **0** |
| VISa_R rest vs MOs_L plan | **0** |
| VISp_L rest vs MOs_L plan | **0** |
| VISpm_L rest vs MOs_L plan | **0** |
| VISpm_R rest vs MOs_L plan | **0** |
| VISp_R rest vs MOs_L plan | **0** |
| VISrl_L rest vs MOs_L plan | **0** |
| VISrl_R rest vs MOs_L plan | **0** |
| RFA_L plan vs MOs_R plan | 1 |
| RFA_R plan vs MOs_R plan | 1 |
| RSPagl_L plan vs MOs_R plan | 1 |
| RSPagl_R plan vs MOs_R plan | 1 |
| RSPd_L plan vs MOs_R plan | 1 |
| RSPd_R plan vs MOs_R plan | 1 |
| SSp-bfd_L plan vs MOs_R plan | 0.065 |
| SSp-bfd_R plan vs MOs_R plan | 0.51 |
| SSp-ll_L plan vs MOs_R plan | 1 |
| SSp-ll_R plan vs MOs_R plan | 1 |
| SSp-n_L plan vs MOs_R plan | 0.27 |
| SSp-n_R plan vs MOs_R plan | 0.59 |

| | |
|---|---|
| SSp-tr_L plan vs MOs_R plan | 1 |
| SSp-tr_R plan vs MOs_R plan | 1 |
| SSp-ul_L plan vs MOs_R plan | 1 |
| SSp-ul_R plan vs MOs_R plan | 1 |
| SSp-un_L plan vs MOs_R plan | 1 |
| SSp-un_R plan vs MOs_R plan | 1 |
| VISa_L plan vs MOs_R plan | 1 |
| VISam_L plan vs MOs_R plan | 0.99 |
| VISam_R plan vs MOs_R plan | 1 |
| VISa_R plan vs MOs_R plan | 1 |
| VISp_L plan vs MOs_R plan | **0.0031** |
| VISpm_L plan vs MOs_R plan | 0.098 |
| VISpm_R plan vs MOs_R plan | 0.48 |
| VISp_R plan vs MOs_R plan | 0.037 |
| VISrl_L plan vs MOs_R plan | 0.015 |
| VISrl_R plan vs MOs_R plan | 0.15 |
| CFA_L rest vs MOs_R plan | **0** |
| CFA_R rest vs MOs_R plan | **2.7e-09** |
| MOp_L rest vs MOs_R plan | **0** |
| MOp_R rest vs MOs_R plan | **0** |
| MOs_L rest vs MOs_R plan | **0** |
| MOs_R rest vs MOs_R plan | **0** |
| RFA_L rest vs MOs_R plan | **0** |
| RFA_R rest vs MOs_R plan | **0** |
| RSPagl_L rest vs MOs_R plan | **0** |
| RSPagl_R rest vs MOs_R plan | **0** |
| RSPd_L rest vs MOs_R plan | **0** |
| RSPd_R rest vs MOs_R plan | **0** |
| SSp-bfd_L rest vs MOs_R plan | **0** |
| SSp-bfd_R rest vs MOs_R plan | **9.6e-10** |
| SSp-ll_L rest vs MOs_R plan | **0** |
| SSp-ll_R rest vs MOs_R plan | **0** |
| SSp-n_L rest vs MOs_R plan | **0** |
| SSp-n_R rest vs MOs_R plan | **5e-08** |
| SSp-tr_L rest vs MOs_R plan | **0** |
| SSp-tr_R rest vs MOs_R plan | **0** |
| SSp-ul_L rest vs MOs_R plan | **0** |
| SSp-ul_R rest vs MOs_R plan | **4.1e-09** |
| SSp-un_L rest vs MOs_R plan | **4.7e-10** |
| SSp-un_R rest vs MOs_R plan | **1.8e-08** |
| VISa_L rest vs MOs_R plan | **0** |
| VISam_L rest vs MOs_R plan | **0** |
| VISam_R rest vs MOs_R plan | **0** |
| VISa_R rest vs MOs_R plan | **0** |
| VISp_L rest vs MOs_R plan | **0** |
| VISpm_L rest vs MOs_R plan | **0** |

| | |
|---|---|
| VISpm_R rest vs MOs_R plan | **0** |
| VISp_R rest vs MOs_R plan | **0** |
| VISrl_L rest vs MOs_R plan | **0** |
| VISrl_R rest vs MOs_R plan | **0** |
| RFA_R plan vs RFA_L plan | 1 |
| RSPagl_L plan vs RFA_L plan | 0.99 |
| RSPagl_R plan vs RFA_L plan | 1 |
| RSPd_L plan vs RFA_L plan | 1 |
| RSPd_R plan vs RFA_L plan | 1 |
| SSp-bfd_L plan vs RFA_L plan | **0.0098** |
| SSp-bfd_R plan vs RFA_L plan | 0.16 |
| SSp-ll_L plan vs RFA_L plan | 1 |
| SSp-ll_R plan vs RFA_L plan | 1 |
| SSp-n_L plan vs RFA_L plan | 0.062 |
| SSp-n_R plan vs RFA_L plan | 0.21 |
| SSp-tr_L plan vs RFA_L plan | 1 |
| SSp-tr_R plan vs RFA_L plan | 1 |
| SSp-ul_L plan vs RFA_L plan | 1 |
| SSp-ul_R plan vs RFA_L plan | 1 |
| SSp-un_L plan vs RFA_L plan | 0.95 |
| SSp-un_R plan vs RFA_L plan | 1 |
| VISa_L plan vs RFA_L plan | 1 |
| VISam_L plan vs RFA_L plan | 0.82 |
| VISam_R plan vs RFA_L plan | 0.98 |
| VISa_R plan vs RFA_L plan | 1 |
| VISp_L plan vs RFA_L plan | **0.00029** |
| VISpm_L plan vs RFA_L plan | 0.016 |
| VISpm_R plan vs RFA_L plan | 0.14 |
| VISp_R plan vs RFA_L plan | **0.0049** |
| VISrl_L plan vs RFA_L plan | **0.0017** |
| VISrl_R plan vs RFA_L plan | 0.027 |
| CFA_L rest vs RFA_L plan | **0** |
| CFA_R rest vs RFA_L plan | **0** |
| MOp_L rest vs RFA_L plan | **0** |
| MOp_R rest vs RFA_L plan | **0** |
| MOs_L rest vs RFA_L plan | **0** |
| MOs_R rest vs RFA_L plan | **0** |
| RFA_L rest vs RFA_L plan | **0** |
| RFA_R rest vs RFA_L plan | **0** |
| RSPagl_L rest vs RFA_L plan | **0** |
| RSPagl_R rest vs RFA_L plan | **0** |
| RSPd_L rest vs RFA_L plan | **0** |
| RSPd_R rest vs RFA_L plan | **0** |
| SSp-bfd_L rest vs RFA_L plan | **0** |
| SSp-bfd_R rest vs RFA_L plan | **0** |
| SSp-ll_L rest vs RFA_L plan | **0** |

| | |
|---|---|
| SSp-ll_R rest vs RFA_L plan | **0** |
| SSp-n_L rest vs RFA_L plan | **0** |
| SSp-n_R rest vs RFA_L plan | **8.4e-10** |
| SSp-tr_L rest vs RFA_L plan | **0** |
| SSp-tr_R rest vs RFA_L plan | **0** |
| SSp-ul_L rest vs RFA_L plan | **0** |
| SSp-ul_R rest vs RFA_L plan | **0** |
| SSp-un_L rest vs RFA_L plan | **0** |
| SSp-un_R rest vs RFA_L plan | **0** |
| VISa_L rest vs RFA_L plan | **0** |
| VISam_L rest vs RFA_L plan | **0** |
| VISam_R rest vs RFA_L plan | **0** |
| VISa_R rest vs RFA_L plan | **0** |
| VISp_L rest vs RFA_L plan | **0** |
| VISpm_L rest vs RFA_L plan | **0** |
| VISpm_R rest vs RFA_L plan | **0** |
| VISp_R rest vs RFA_L plan | **0** |
| VISrl_L rest vs RFA_L plan | **0** |
| VISrl_R rest vs RFA_L plan | **0** |
| RSPagl_L plan vs RFA_R plan | 1 |
| RSPagl_R plan vs RFA_R plan | 1 |
| RSPd_L plan vs RFA_R plan | 1 |
| RSPd_R plan vs RFA_R plan | 1 |
| SSp-bfd_L plan vs RFA_R plan | 0.16 |
| SSp-bfd_R plan vs RFA_R plan | 0.76 |
| SSp-ll_L plan vs RFA_R plan | 1 |
| SSp-ll_R plan vs RFA_R plan | 1 |
| SSp-n_L plan vs RFA_R plan | 0.5 |
| SSp-n_R plan vs RFA_R plan | 0.82 |
| SSp-tr_L plan vs RFA_R plan | 1 |
| SSp-tr_R plan vs RFA_R plan | 1 |
| SSp-ul_L plan vs RFA_R plan | 1 |
| SSp-ul_R plan vs RFA_R plan | 1 |
| SSp-un_L plan vs RFA_R plan | 1 |
| SSp-un_R plan vs RFA_R plan | 1 |
| VISa_L plan vs RFA_R plan | 1 |
| VISam_L plan vs RFA_R plan | 1 |
| VISam_R plan vs RFA_R plan | 1 |
| VISa_R plan vs RFA_R plan | 1 |
| VISp_L plan vs RFA_R plan | 0.011 |
| VISpm_L plan vs RFA_R plan | 0.22 |
| VISpm_R plan vs RFA_R plan | 0.72 |
| VISp_R plan vs RFA_R plan | 0.097 |
| VISrl_L plan vs RFA_R plan | 0.044 |
| VISrl_R plan vs RFA_R plan | 0.31 |
| CFA_L rest vs RFA_R plan | **1.9e-09** |

| | |
|---|---|
| CFA_R rest vs RFA_R plan | **2.3e-08** |
| MOp_L rest vs RFA_R plan | **0** |
| MOp_R rest vs RFA_R plan | **2e-09** |
| MOs_L rest vs RFA_R plan | **0** |
| MOs_R rest vs RFA_R plan | **0** |
| RFA_L rest vs RFA_R plan | **0** |
| RFA_R rest vs RFA_R plan | **0** |
| RSPagl_L rest vs RFA_R plan | **0** |
| RSPagl_R rest vs RFA_R plan | **0** |
| RSPd_L rest vs RFA_R plan | **0** |
| RSPd_R rest vs RFA_R plan | **0** |
| SSp-bfd_L rest vs RFA_R plan | **0** |
| SSp-bfd_R rest vs RFA_R plan | **1.2e-08** |
| SSp-ll_L rest vs RFA_R plan | **0** |
| SSp-ll_R rest vs RFA_R plan | **3.7e-09** |
| SSp-n_L rest vs RFA_R plan | **4.6e-09** |
| SSp-n_R rest vs RFA_R plan | **3e-07** |
| SSp-tr_L rest vs RFA_R plan | **0** |
| SSp-tr_R rest vs RFA_R plan | **0** |
| SSp-ul_L rest vs RFA_R plan | **3.7e-09** |
| SSp-ul_R rest vs RFA_R plan | **3.2e-08** |
| SSp-un_L rest vs RFA_R plan | **9.1e-09** |
| SSp-un_R rest vs RFA_R plan | **1.1e-07** |
| VISa_L rest vs RFA_R plan | **0** |
| VISam_L rest vs RFA_R plan | **0** |
| VISam_R rest vs RFA_R plan | **0** |
| VISa_R rest vs RFA_R plan | **0** |
| VISp_L rest vs RFA_R plan | **0** |
| VISpm_L rest vs RFA_R plan | **0** |
| VISpm_R rest vs RFA_R plan | **0** |
| VISp_R rest vs RFA_R plan | **0** |
| VISrl_L rest vs RFA_R plan | **0** |
| VISrl_R rest vs RFA_R plan | **0** |
| RSPagl_R plan vs RSPagl_L plan | 1 |
| RSPd_L plan vs RSPagl_L plan | 1 |
| RSPd_R plan vs RSPagl_L plan | 1 |
| SSp-bfd_L plan vs RSPagl_L plan | 1 |
| SSp-bfd_R plan vs RSPagl_L plan | 1 |
| SSp-ll_L plan vs RSPagl_L plan | 1 |
| SSp-ll_R plan vs RSPagl_L plan | 0.99 |
| SSp-n_L plan vs RSPagl_L plan | 1 |
| SSp-n_R plan vs RSPagl_L plan | 1 |
| SSp-tr_L plan vs RSPagl_L plan | 1 |
| SSp-tr_R plan vs RSPagl_L plan | 1 |
| SSp-ul_L plan vs RSPagl_L plan | 1 |
| SSp-ul_R plan vs RSPagl_L plan | 1 |

| | |
|---|---|
| SSp-un_L plan vs RSPagl_L plan | 1 |
| SSp-un_R plan vs RSPagl_L plan | 1 |
| VISa_L plan vs RSPagl_L plan | 1 |
| VISam_L plan vs RSPagl_L plan | 1 |
| VISam_R plan vs RSPagl_L plan | 1 |
| VISa_R plan vs RSPagl_L plan | 1 |
| VISp_L plan vs RSPagl_L plan | 0.99 |
| VISpm_L plan vs RSPagl_L plan | 1 |
| VISpm_R plan vs RSPagl_L plan | 1 |
| VISp_R plan vs RSPagl_L plan | 1 |
| VISrl_L plan vs RSPagl_L plan | 1 |
| VISrl_R plan vs RSPagl_L plan | 1 |
| CFA_L rest vs RSPagl_L plan | **0.00067** |
| CFA_R rest vs RSPagl_L plan | **0.003** |
| MOp_L rest vs RSPagl_L plan | **0.00022** |
| MOp_R rest vs RSPagl_L plan | **0.00069** |
| MOs_L rest vs RSPagl_L plan | **6e-05** |
| MOs_R rest vs RSPagl_L plan | **4.1e-05** |
| RFA_L rest vs RSPagl_L plan | **7e-06** |
| RFA_R rest vs RSPagl_L plan | **2.2e-06** |
| RSPagl_L rest vs RSPagl_L plan | **2.6e-07** |
| RSPagl_R rest vs RSPagl_L plan | **8.1e-07** |
| RSPd_L rest vs RSPagl_L plan | **4.4e-07** |
| RSPd_R rest vs RSPagl_L plan | **6.6e-07** |
| SSp-bfd_L rest vs RSPagl_L plan | **0.00017** |
| SSp-bfd_R rest vs RSPagl_L plan | **0.0019** |
| SSp-ll_L rest vs RSPagl_L plan | **3.7e-05** |
| SSp-ll_R rest vs RSPagl_L plan | **0.00095** |
| SSp-n_L rest vs RSPagl_L plan | **0.0011** |
| SSp-n_R rest vs RSPagl_L plan | 0.016 |
| SSp-tr_L rest vs RSPagl_L plan | **9.8e-06** |
| SSp-tr_R rest vs RSPagl_L plan | **0.00011** |
| SSp-ul_L rest vs RSPagl_L plan | **0.00094** |
| SSp-ul_R rest vs RSPagl_L plan | **0.0036** |
| SSp-un_L rest vs RSPagl_L plan | **0.0016** |
| SSp-un_R rest vs RSPagl_L plan | **0.0085** |
| VISa_L rest vs RSPagl_L plan | **5.6e-06** |
| VISam_L rest vs RSPagl_L plan | **7.7e-07** |
| VISam_R rest vs RSPagl_L plan | **1.8e-06** |
| VISa_R rest vs RSPagl_L plan | **1.1e-05** |
| VISp_L rest vs RSPagl_L plan | **1.8e-07** |
| VISpm_L rest vs RSPagl_L plan | **5.3e-08** |
| VISpm_R rest vs RSPagl_L plan | **1.8e-07** |
| VISp_R rest vs RSPagl_L plan | **5.6e-07** |
| VISrl_L rest vs RSPagl_L plan | **3.6e-06** |
| VISrl_R rest vs RSPagl_L plan | **1.3e-05** |

| | |
|---|---|
| RSPd_L plan vs RSPagl_R plan | 1 |
| RSPd_R plan vs RSPagl_R plan | 1 |
| SSp-bfd_L plan vs RSPagl_R plan | 1 |
| SSp-bfd_R plan vs RSPagl_R plan | 1 |
| SSp-ll_L plan vs RSPagl_R plan | 1 |
| SSp-ll_R plan vs RSPagl_R plan | 1 |
| SSp-n_L plan vs RSPagl_R plan | 1 |
| SSp-n_R plan vs RSPagl_R plan | 1 |
| SSp-tr_L plan vs RSPagl_R plan | 1 |
| SSp-tr_R plan vs RSPagl_R plan | 1 |
| SSp-ul_L plan vs RSPagl_R plan | 1 |
| SSp-ul_R plan vs RSPagl_R plan | 1 |
| SSp-un_L plan vs RSPagl_R plan | 1 |
| SSp-un_R plan vs RSPagl_R plan | 1 |
| VISa_L plan vs RSPagl_R plan | 1 |
| VISam_L plan vs RSPagl_R plan | 1 |
| VISam_R plan vs RSPagl_R plan | 1 |
| VISa_R plan vs RSPagl_R plan | 1 |
| VISp_L plan vs RSPagl_R plan | 0.91 |
| VISpm_L plan vs RSPagl_R plan | 1 |
| VISpm_R plan vs RSPagl_R plan | 1 |
| VISp_R plan vs RSPagl_R plan | 1 |
| VISrl_L plan vs RSPagl_R plan | 0.99 |
| VISrl_R plan vs RSPagl_R plan | 1 |
| CFA_L rest vs RSPagl_R plan | **0.00015** |
| CFA_R rest vs RSPagl_R plan | **0.00073** |
| MOp_L rest vs RSPagl_R plan | **4.6e-05** |
| MOp_R rest vs RSPagl_R plan | **0.00015** |
| MOs_L rest vs RSPagl_R plan | **1.2e-05** |
| MOs_R rest vs RSPagl_R plan | **7.6e-06** |
| RFA_L rest vs RSPagl_R plan | **1.2e-06** |
| RFA_R rest vs RSPagl_R plan | **3.6e-07** |
| RSPagl_L rest vs RSPagl_R plan | **3.8e-08** |
| RSPagl_R rest vs RSPagl_R plan | **1.2e-07** |
| RSPd_L rest vs RSPagl_R plan | **6.6e-08** |
| RSPd_R rest vs RSPagl_R plan | **1e-07** |
| SSp-bfd_L rest vs RSPagl_R plan | **3.5e-05** |
| SSp-bfd_R rest vs RSPagl_R plan | **0.00046** |
| SSp-ll_L rest vs RSPagl_R plan | **6.9e-06** |
| SSp-ll_R rest vs RSPagl_R plan | **0.00021** |
| SSp-n_L rest vs RSPagl_R plan | **0.00024** |
| SSp-n_R rest vs RSPagl_R plan | **0.0045** |
| SSp-tr_L rest vs RSPagl_R plan | **1.7e-06** |
| SSp-tr_R rest vs RSPagl_R plan | **2.1e-05** |
| SSp-ul_L rest vs RSPagl_R plan | **0.00021** |
| SSp-ul_R rest vs RSPagl_R plan | **0.00091** |

| | |
|---|---|
| SSp-un_L rest vs RSPagl_R plan | **0.00038** |
| SSp-un_R rest vs RSPagl_R plan | **0.0023** |
| VISa_L rest vs RSPagl_R plan | **9.5e-07** |
| VISam_L rest vs RSPagl_R plan | **1.2e-07** |
| VISam_R rest vs RSPagl_R plan | **2.9e-07** |
| VISa_R rest vs RSPagl_R plan | **2e-06** |
| VISp_L rest vs RSPagl_R plan | **2.6e-08** |
| VISpm_L rest vs RSPagl_R plan | **6.4e-09** |
| VISpm_R rest vs RSPagl_R plan | **2.6e-08** |
| VISp_R rest vs RSPagl_R plan | **8.5e-08** |
| VISrl_L rest vs RSPagl_R plan | **6e-07** |
| VISrl_R rest vs RSPagl_R plan | **2.3e-06** |
| RSPd_R plan vs RSPd_L plan | 1 |
| SSp-bfd_L plan vs RSPd_L plan | 0.77 |
| SSp-bfd_R plan vs RSPd_L plan | 1 |
| SSp-ll_L plan vs RSPd_L plan | 1 |
| SSp-ll_R plan vs RSPd_L plan | 1 |
| SSp-n_L plan vs RSPd_L plan | 0.98 |
| SSp-n_R plan vs RSPd_L plan | 1 |
| SSp-tr_L plan vs RSPd_L plan | 1 |
| SSp-tr_R plan vs RSPd_L plan | 1 |
| SSp-ul_L plan vs RSPd_L plan | 1 |
| SSp-ul_R plan vs RSPd_L plan | 1 |
| SSp-un_L plan vs RSPd_L plan | 1 |
| SSp-un_R plan vs RSPd_L plan | 1 |
| VISa_L plan vs RSPd_L plan | 1 |
| VISam_L plan vs RSPd_L plan | 1 |
| VISam_R plan vs RSPd_L plan | 1 |
| VISa_R plan vs RSPd_L plan | 1 |
| VISp_L plan vs RSPd_L plan | 0.17 |
| VISpm_L plan vs RSPd_L plan | 0.85 |
| VISpm_R plan vs RSPd_L plan | 1 |
| VISp_R plan vs RSPd_L plan | 0.63 |
| VISrl_L plan vs RSPd_L plan | 0.42 |
| VISrl_R plan vs RSPd_L plan | 0.92 |
| CFA_L rest vs RSPd_L plan | **5.1e-07** |
| CFA_R rest vs RSPd_L plan | **3.4e-06** |
| MOp_L rest vs RSPd_L plan | **1.3e-07** |
| MOp_R rest vs RSPd_L plan | **5.3e-07** |
| MOs_L rest vs RSPd_L plan | **2.5e-08** |
| MOs_R rest vs RSPd_L plan | **1.5e-08** |
| RFA_L rest vs RSPd_L plan | **8.7e-10** |
| RFA_R rest vs RSPd_L plan | **0** |
| RSPagl_L rest vs RSPd_L plan | **0** |
| RSPagl_R rest vs RSPd_L plan | **0** |
| RSPd_L rest vs RSPd_L plan | **0** |

| | |
|---|---|
| RSPd_R rest vs RSPd_L plan | **0** |
| SSp-bfd_L rest vs RSPd_L plan | **9.3e-08** |
| SSp-bfd_R rest vs RSPd_L plan | **2e-06** |
| SSp-ll_L rest vs RSPd_L plan | **1.3e-08** |
| SSp-ll_R rest vs RSPd_L plan | **7.9e-07** |
| SSp-n_L rest vs RSPd_L plan | **9.2e-07** |
| SSp-n_R rest vs RSPd_L plan | **3.2e-05** |
| SSp-tr_L rest vs RSPd_L plan | **1.8e-09** |
| SSp-tr_R rest vs RSPd_L plan | **5.2e-08** |
| SSp-ul_L rest vs RSPd_L plan | **7.8e-07** |
| SSp-ul_R rest vs RSPd_L plan | **4.4e-06** |
| SSp-un_L rest vs RSPd_L plan | **1.6e-06** |
| SSp-un_R rest vs RSPd_L plan | **1.4e-05** |
| VISa_L rest vs RSPd_L plan | **4.5e-10** |
| VISam_L rest vs RSPd_L plan | **0** |
| VISam_R rest vs RSPd_L plan | **0** |
| VISa_R rest vs RSPd_L plan | **2.4e-09** |
| VISp_L rest vs RSPd_L plan | **0** |
| VISpm_L rest vs RSPd_L plan | **0** |
| VISpm_R rest vs RSPd_L plan | **0** |
| VISp_R rest vs RSPd_L plan | **0** |
| VISrl_L rest vs RSPd_L plan | **0** |
| VISrl_R rest vs RSPd_L plan | **3e-09** |
| SSp-bfd_L plan vs RSPd_R plan | 0.85 |
| SSp-bfd_R plan vs RSPd_R plan | 1 |
| SSp-ll_L plan vs RSPd_R plan | 1 |
| SSp-ll_R plan vs RSPd_R plan | 1 |
| SSp-n_L plan vs RSPd_R plan | 0.99 |
| SSp-n_R plan vs RSPd_R plan | 1 |
| SSp-tr_L plan vs RSPd_R plan | 1 |
| SSp-tr_R plan vs RSPd_R plan | 1 |
| SSp-ul_L plan vs RSPd_R plan | 1 |
| SSp-ul_R plan vs RSPd_R plan | 1 |
| SSp-un_L plan vs RSPd_R plan | 1 |
| SSp-un_R plan vs RSPd_R plan | 1 |
| VISa_L plan vs RSPd_R plan | 1 |
| VISam_L plan vs RSPd_R plan | 1 |
| VISam_R plan vs RSPd_R plan | 1 |
| VISa_R plan vs RSPd_R plan | 1 |
| VISp_L plan vs RSPd_R plan | 0.24 |
| VISpm_L plan vs RSPd_R plan | 0.92 |
| VISpm_R plan vs RSPd_R plan | 1 |
| VISp_R plan vs RSPd_R plan | 0.74 |
| VISrl_L plan vs RSPd_R plan | 0.53 |
| VISrl_R plan vs RSPd_R plan | 0.96 |
| CFA_L rest vs RSPd_R plan | **1.1e-06** |

| | |
|---|---|
| CFA_R rest vs RSPd_R plan | **7.1e-06** |
| MOp_L rest vs RSPd_R plan | **2.9e-07** |
| MOp_R rest vs RSPd_R plan | **1.1e-06** |
| MOs_L rest vs RSPd_R plan | **5.7e-08** |
| MOs_R rest vs RSPd_R plan | **3.5e-08** |
| RFA_L rest vs RSPd_R plan | **3.4e-09** |
| RFA_R rest vs RSPd_R plan | **1.2e-10** |
| RSPagl_L rest vs RSPd_R plan | **0** |
| RSPagl_R rest vs RSPd_R plan | **0** |
| RSPd_L rest vs RSPd_R plan | **0** |
| RSPd_R rest vs RSPd_R plan | **0** |
| SSp-bfd_L rest vs RSPd_R plan | **2.1e-07** |
| SSp-bfd_R rest vs RSPd_R plan | **4.1e-06** |
| SSp-ll_L rest vs RSPd_R plan | **3.1e-08** |
| SSp-ll_R rest vs RSPd_R plan | **1.7e-06** |
| SSp-n_L rest vs RSPd_R plan | **1.9e-06** |
| SSp-n_R rest vs RSPd_R plan | **6.3e-05** |
| SSp-tr_L rest vs RSPd_R plan | **5.7e-09** |
| SSp-tr_R rest vs RSPd_R plan | **1.2e-07** |
| SSp-ul_L rest vs RSPd_R plan | **1.7e-06** |
| SSp-ul_R rest vs RSPd_R plan | **9.1e-06** |
| SSp-un_L rest vs RSPd_R plan | **3.3e-06** |
| SSp-un_R rest vs RSPd_R plan | **2.7e-05** |
| VISa_L rest vs RSPd_R plan | **2.5e-09** |
| VISam_L rest vs RSPd_R plan | **0** |
| VISam_R rest vs RSPd_R plan | **0** |
| VISa_R rest vs RSPd_R plan | **7e-09** |
| VISp_L rest vs RSPd_R plan | **0** |
| VISpm_L rest vs RSPd_R plan | **0** |
| VISpm_R rest vs RSPd_R plan | **0** |
| VISp_R rest vs RSPd_R plan | **0** |
| VISrl_L rest vs RSPd_R plan | **1e-09** |
| VISrl_R rest vs RSPd_R plan | **8.4e-09** |
| SSp-bfd_R plan vs SSp-bfd_L plan | 1 |
| SSp-ll_L plan vs SSp-bfd_L plan | 0.08 |
| SSp-ll_R plan vs SSp-bfd_L plan | 0.011 |
| SSp-n_L plan vs SSp-bfd_L plan | 1 |
| SSp-n_R plan vs SSp-bfd_L plan | 1 |
| SSp-tr_L plan vs SSp-bfd_L plan | 0.56 |
| SSp-tr_R plan vs SSp-bfd_L plan | 0.11 |
| SSp-ul_L plan vs SSp-bfd_L plan | 0.98 |
| SSp-ul_R plan vs SSp-bfd_L plan | 0.84 |
| SSp-un_L plan vs SSp-bfd_L plan | 1 |
| SSp-un_R plan vs SSp-bfd_L plan | 1 |
| VISa_L plan vs SSp-bfd_L plan | 1 |
| VISam_L plan vs SSp-bfd_L plan | 1 |

| | |
|---|---|
| VISam_R plan vs SSp-bfd_L plan | 1 |
| VISa_R plan vs SSp-bfd_L plan | 1 |
| VISp_L plan vs SSp-bfd_L plan | 1 |
| VISpm_L plan vs SSp-bfd_L plan | 1 |
| VISpm_R plan vs SSp-bfd_L plan | 1 |
| VISp_R plan vs SSp-bfd_L plan | 1 |
| VISrl_L plan vs SSp-bfd_L plan | 1 |
| VISrl_R plan vs SSp-bfd_L plan | 1 |
| CFA_L rest vs SSp-bfd_L plan | 0.76 |
| CFA_R rest vs SSp-bfd_L plan | 0.94 |
| MOp_L rest vs SSp-bfd_L plan | 0.58 |
| MOp_R rest vs SSp-bfd_L plan | 0.77 |
| MOs_L rest vs SSp-bfd_L plan | 0.36 |
| MOs_R rest vs SSp-bfd_L plan | 0.31 |
| RFA_L rest vs SSp-bfd_L plan | 0.13 |
| RFA_R rest vs SSp-bfd_L plan | 0.069 |
| RSPagl_L rest vs SSp-bfd_L plan | 0.019 |
| RSPagl_R rest vs SSp-bfd_L plan | 0.039 |
| RSPd_L rest vs SSp-bfd_L plan | 0.027 |
| RSPd_R rest vs SSp-bfd_L plan | 0.034 |
| SSp-bfd_L rest vs SSp-bfd_L plan | 0.53 |
| SSp-bfd_R rest vs SSp-bfd_L plan | 0.9 |
| SSp-ll_L rest vs SSp-bfd_L plan | 0.29 |
| SSp-ll_R rest vs SSp-bfd_L plan | 0.82 |
| SSp-n_L rest vs SSp-bfd_L plan | 0.83 |
| SSp-n_R rest vs SSp-bfd_L plan | 1 |
| SSp-tr_L rest vs SSp-bfd_L plan | 0.16 |
| SSp-tr_R rest vs SSp-bfd_L plan | 0.45 |
| SSp-ul_L rest vs SSp-bfd_L plan | 0.82 |
| SSp-ul_R rest vs SSp-bfd_L plan | 0.96 |
| SSp-un_L rest vs SSp-bfd_L plan | 0.88 |
| SSp-un_R rest vs SSp-bfd_L plan | 0.99 |
| VISa_L rest vs SSp-bfd_L plan | 0.12 |
| VISam_L rest vs SSp-bfd_L plan | 0.038 |
| VISam_R rest vs SSp-bfd_L plan | 0.062 |
| VISa_R rest vs SSp-bfd_L plan | 0.17 |
| VISp_L rest vs SSp-bfd_L plan | 0.016 |
| VISpm_L rest vs SSp-bfd_L plan | **0.0069** |
| VISpm_R rest vs SSp-bfd_L plan | 0.015 |
| VISp_R rest vs SSp-bfd_L plan | 0.031 |
| VISrl_L rest vs SSp-bfd_L plan | 0.092 |
| VISrl_R rest vs SSp-bfd_L plan | 0.18 |
| SSp-ll_L plan vs SSp-bfd_R plan | 0.57 |
| SSp-ll_R plan vs SSp-bfd_R plan | 0.17 |
| SSp-n_L plan vs SSp-bfd_R plan | 1 |
| SSp-n_R plan vs SSp-bfd_R plan | 1 |

| | |
|---|---|
| SSp-tr_L plan vs SSp-bfd_R plan | 0.99 |
| SSp-tr_R plan vs SSp-bfd_R plan | 0.67 |
| SSp-ul_L plan vs SSp-bfd_R plan | 1 |
| SSp-ul_R plan vs SSp-bfd_R plan | 1 |
| SSp-un_L plan vs SSp-bfd_R plan | 1 |
| SSp-un_R plan vs SSp-bfd_R plan | 1 |
| VISa_L plan vs SSp-bfd_R plan | 1 |
| VISam_L plan vs SSp-bfd_R plan | 1 |
| VISam_R plan vs SSp-bfd_R plan | 1 |
| VISa_R plan vs SSp-bfd_R plan | 1 |
| VISp_L plan vs SSp-bfd_R plan | 1 |
| VISpm_L plan vs SSp-bfd_R plan | 1 |
| VISpm_R plan vs SSp-bfd_R plan | 1 |
| VISp_R plan vs SSp-bfd_R plan | 1 |
| VISrl_L plan vs SSp-bfd_R plan | 1 |
| VISrl_R plan vs SSp-bfd_R plan | 1 |
| CFA_L rest vs SSp-bfd_R plan | 0.16 |
| CFA_R rest vs SSp-bfd_R plan | 0.36 |
| MOp_L rest vs SSp-bfd_R plan | 0.083 |
| MOp_R rest vs SSp-bfd_R plan | 0.17 |
| MOs_L rest vs SSp-bfd_R plan | 0.034 |
| MOs_R rest vs SSp-bfd_R plan | 0.026 |
| RFA_L rest vs SSp-bfd_R plan | **0.0073** |
| RFA_R rest vs SSp-bfd_R plan | **0.0031** |
| RSPagl_L rest vs SSp-bfd_R plan | **0.0006** |
| RSPagl_R rest vs SSp-bfd_R plan | **0.0014** |
| RSPd_L rest vs SSp-bfd_R plan | **0.00088** |
| RSPd_R rest vs SSp-bfd_R plan | **0.0012** |
| SSp-bfd_L rest vs SSp-bfd_R plan | 0.07 |
| SSp-bfd_R rest vs SSp-bfd_R plan | 0.29 |
| SSp-ll_L rest vs SSp-bfd_R plan | 0.024 |
| SSp-ll_R rest vs SSp-bfd_R plan | 0.2 |
| SSp-n_L rest vs SSp-bfd_R plan | 0.21 |
| SSp-n_R rest vs SSp-bfd_R plan | 0.71 |
| SSp-tr_L rest vs SSp-bfd_R plan | **0.0094** |
| SSp-tr_R rest vs SSp-bfd_R plan | 0.051 |
| SSp-ul_L rest vs SSp-bfd_R plan | 0.2 |
| SSp-ul_R rest vs SSp-bfd_R plan | 0.4 |
| SSp-un_L rest vs SSp-bfd_R plan | 0.27 |
| SSp-un_R rest vs SSp-bfd_R plan | 0.57 |
| VISa_L rest vs SSp-bfd_R plan | **0.0062** |
| VISam_L rest vs SSp-bfd_R plan | **0.0014** |
| VISam_R rest vs SSp-bfd_R plan | **0.0026** |
| VISa_R rest vs SSp-bfd_R plan | 0.01 |
| VISp_L rest vs SSp-bfd_R plan | **0.00045** |
| VISpm_L rest vs SSp-bfd_R plan | **0.00017** |

| | |
|---|---|
| VISpm_R rest vs SSp-bfd_R plan | **0.00045** |
| VISp_R rest vs SSp-bfd_R plan | **0.0011** |
| VISrl_L rest vs SSp-bfd_R plan | **0.0045** |
| VISrl_R rest vs SSp-bfd_R plan | 0.012 |
| SSp-ll_R plan vs SSp-ll_L plan | 1 |
| SSp-n_L plan vs SSp-ll_L plan | 0.31 |
| SSp-n_R plan vs SSp-ll_L plan | 0.64 |
| SSp-tr_L plan vs SSp-ll_L plan | 1 |
| SSp-tr_R plan vs SSp-ll_L plan | 1 |
| SSp-ul_L plan vs SSp-ll_L plan | 1 |
| SSp-ul_R plan vs SSp-ll_L plan | 1 |
| SSp-un_L plan vs SSp-ll_L plan | 1 |
| SSp-un_R plan vs SSp-ll_L plan | 1 |
| VISa_L plan vs SSp-ll_L plan | 1 |
| VISam_L plan vs SSp-ll_L plan | 0.99 |
| VISam_R plan vs SSp-ll_L plan | 1 |
| VISa_R plan vs SSp-ll_L plan | 1 |
| VISp_L plan vs SSp-ll_L plan | **0.0041** |
| VISpm_L plan vs SSp-ll_L plan | 0.12 |
| VISpm_R plan vs SSp-ll_L plan | 0.53 |
| VISp_R plan vs SSp-ll_L plan | 0.046 |
| VISrl_L plan vs SSp-ll_L plan | 0.019 |
| VISrl_R plan vs SSp-ll_L plan | 0.17 |
| CFA_L rest vs SSp-ll_L plan | **0** |
| CFA_R rest vs SSp-ll_L plan | **4.6e-09** |
| MOp_L rest vs SSp-ll_L plan | **0** |
| MOp_R rest vs SSp-ll_L plan | **0** |
| MOs_L rest vs SSp-ll_L plan | **0** |
| MOs_R rest vs SSp-ll_L plan | **0** |
| RFA_L rest vs SSp-ll_L plan | **0** |
| RFA_R rest vs SSp-ll_L plan | **0** |
| RSPagl_L rest vs SSp-ll_L plan | **0** |
| RSPagl_R rest vs SSp-ll_L plan | **0** |
| RSPd_L rest vs SSp-ll_L plan | **0** |
| RSPd_R rest vs SSp-ll_L plan | **0** |
| SSp-bfd_L rest vs SSp-ll_L plan | **0** |
| SSp-bfd_R rest vs SSp-ll_L plan | **2e-09** |
| SSp-ll_L rest vs SSp-ll_L plan | **0** |
| SSp-ll_R rest vs SSp-ll_L plan | **2e-11** |
| SSp-n_L rest vs SSp-ll_L plan | **2.2e-10** |
| SSp-n_R rest vs SSp-ll_L plan | **7.5e-08** |
| SSp-tr_L rest vs SSp-ll_L plan | **0** |
| SSp-tr_R rest vs SSp-ll_L plan | **0** |
| SSp-ul_L rest vs SSp-ll_L plan | **1e-11** |
| SSp-ul_R rest vs SSp-ll_L plan | **6.6e-09** |
| SSp-un_L rest vs SSp-ll_L plan | **1.2e-09** |

| | |
|---|---|
| SSp-un_R rest vs SSp-ll_L plan | **2.7e-08** |
| VISa_L rest vs SSp-ll_L plan | **0** |
| VISam_L rest vs SSp-ll_L plan | **0** |
| VISam_R rest vs SSp-ll_L plan | **0** |
| VISa_R rest vs SSp-ll_L plan | **0** |
| VISp_L rest vs SSp-ll_L plan | **0** |
| VISpm_L rest vs SSp-ll_L plan | **0** |
| VISpm_R rest vs SSp-ll_L plan | **0** |
| VISp_R rest vs SSp-ll_L plan | **0** |
| VISrl_L rest vs SSp-ll_L plan | **0** |
| VISrl_R rest vs SSp-ll_L plan | **0** |
| SSp-n_L plan vs SSp-ll_R plan | 0.068 |
| SSp-n_R plan vs SSp-ll_R plan | 0.22 |
| SSp-tr_L plan vs SSp-ll_R plan | 1 |
| SSp-tr_R plan vs SSp-ll_R plan | 1 |
| SSp-ul_L plan vs SSp-ll_R plan | 1 |
| SSp-ul_R plan vs SSp-ll_R plan | 1 |
| SSp-un_L plan vs SSp-ll_R plan | 0.96 |
| SSp-un_R plan vs SSp-ll_R plan | 1 |
| VISa_L plan vs SSp-ll_R plan | 1 |
| VISam_L plan vs SSp-ll_R plan | 0.84 |
| VISam_R plan vs SSp-ll_R plan | 0.98 |
| VISa_R plan vs SSp-ll_R plan | 1 |
| VISp_L plan vs SSp-ll_R plan | **0.00033** |
| VISpm_L plan vs SSp-ll_R plan | 0.018 |
| VISpm_R plan vs SSp-ll_R plan | 0.16 |
| VISp_R plan vs SSp-ll_R plan | **0.0056** |
| VISrl_L plan vs SSp-ll_R plan | **0.0019** |
| VISrl_R plan vs SSp-ll_R plan | 0.03 |
| CFA_L rest vs SSp-ll_R plan | **0** |
| CFA_R rest vs SSp-ll_R plan | **0** |
| MOp_L rest vs SSp-ll_R plan | **0** |
| MOp_R rest vs SSp-ll_R plan | **0** |
| MOs_L rest vs SSp-ll_R plan | **0** |
| MOs_R rest vs SSp-ll_R plan | **0** |
| RFA_L rest vs SSp-ll_R plan | **0** |
| RFA_R rest vs SSp-ll_R plan | **0** |
| RSPagl_L rest vs SSp-ll_R plan | **0** |
| RSPagl_R rest vs SSp-ll_R plan | **0** |
| RSPd_L rest vs SSp-ll_R plan | **0** |
| RSPd_R rest vs SSp-ll_R plan | **0** |
| SSp-bfd_L rest vs SSp-ll_R plan | **0** |
| SSp-bfd_R rest vs SSp-ll_R plan | **0** |
| SSp-ll_L rest vs SSp-ll_R plan | **0** |
| SSp-ll_R rest vs SSp-ll_R plan | **0** |
| SSp-n_L rest vs SSp-ll_R plan | **0** |

| | |
|---|---|
| SSp-n_R rest vs SSp-ll_R plan | **1.2e-09** |
| SSp-tr_L rest vs SSp-ll_R plan | **0** |
| SSp-tr_R rest vs SSp-ll_R plan | **0** |
| SSp-ul_L rest vs SSp-ll_R plan | **0** |
| SSp-ul_R rest vs SSp-ll_R plan | **0** |
| SSp-un_L rest vs SSp-ll_R plan | **0** |
| SSp-un_R rest vs SSp-ll_R plan | **0** |
| VISa_L rest vs SSp-ll_R plan | **0** |
| VISam_L rest vs SSp-ll_R plan | **0** |
| VISam_R rest vs SSp-ll_R plan | **0** |
| VISa_R rest vs SSp-ll_R plan | **0** |
| VISp_L rest vs SSp-ll_R plan | **0** |
| VISpm_L rest vs SSp-ll_R plan | **0** |
| VISpm_R rest vs SSp-ll_R plan | **0** |
| VISp_R rest vs SSp-ll_R plan | **0** |
| VISrl_L rest vs SSp-ll_R plan | **0** |
| VISrl_R rest vs SSp-ll_R plan | **0** |
| SSp-n_R plan vs SSp-n_L plan | 1 |
| SSp-tr_L plan vs SSp-n_L plan | 0.91 |
| SSp-tr_R plan vs SSp-n_L plan | 0.4 |
| SSp-ul_L plan vs SSp-n_L plan | 1 |
| SSp-ul_R plan vs SSp-n_L plan | 0.99 |
| SSp-un_L plan vs SSp-n_L plan | 1 |
| SSp-un_R plan vs SSp-n_L plan | 1 |
| VISa_L plan vs SSp-n_L plan | 1 |
| VISam_L plan vs SSp-n_L plan | 1 |
| VISam_R plan vs SSp-n_L plan | 1 |
| VISa_R plan vs SSp-n_L plan | 1 |
| VISp_L plan vs SSp-n_L plan | 1 |
| VISpm_L plan vs SSp-n_L plan | 1 |
| VISpm_R plan vs SSp-n_L plan | 1 |
| VISp_R plan vs SSp-n_L plan | 1 |
| VISrl_L plan vs SSp-n_L plan | 1 |
| VISrl_R plan vs SSp-n_L plan | 1 |
| CFA_L rest vs SSp-n_L plan | 0.35 |
| CFA_R rest vs SSp-n_L plan | 0.63 |
| MOp_L rest vs SSp-n_L plan | 0.21 |
| MOp_R rest vs SSp-n_L plan | 0.36 |
| MOs_L rest vs SSp-n_L plan | 0.098 |
| MOs_R rest vs SSp-n_L plan | 0.077 |
| RFA_L rest vs SSp-n_L plan | 0.025 |
| RFA_R rest vs SSp-n_L plan | 0.011 |
| RSPagl_L rest vs SSp-n_L plan | **0.0025** |
| RSPagl_R rest vs SSp-n_L plan | **0.0056** |
| RSPd_L rest vs SSp-n_L plan | **0.0036** |
| RSPd_R rest vs SSp-n_L plan | **0.0048** |

| | |
|---|---|
| SSp-bfd_L rest vs SSp-n_L plan | 0.18 |
| SSp-bfd_R rest vs SSp-n_L plan | 0.54 |
| SSp-ll_L rest vs SSp-n_L plan | 0.072 |
| SSp-ll_R rest vs SSp-n_L plan | 0.41 |
| SSp-n_L rest vs SSp-n_L plan | 0.43 |
| SSp-n_R rest vs SSp-n_L plan | 0.91 |
| SSp-tr_L rest vs SSp-n_L plan | 0.031 |
| SSp-tr_R rest vs SSp-n_L plan | 0.14 |
| SSp-ul_L rest vs SSp-n_L plan | 0.41 |
| SSp-ul_R rest vs SSp-n_L plan | 0.67 |
| SSp-un_L rest vs SSp-n_L plan | 0.51 |
| SSp-un_R rest vs SSp-n_L plan | 0.82 |
| VISa_L rest vs SSp-n_L plan | 0.022 |
| VISam_L rest vs SSp-n_L plan | **0.0054** |
| VISam_R rest vs SSp-n_L plan | **0.0098** |
| VISa_R rest vs SSp-n_L plan | 0.034 |
| VISp_L rest vs SSp-n_L plan | **0.0019** |
| VISpm_L rest vs SSp-n_L plan | **0.00076** |
| VISpm_R rest vs SSp-n_L plan | **0.0019** |
| VISp_R rest vs SSp-n_L plan | **0.0043** |
| VISrl_L rest vs SSp-n_L plan | 0.016 |
| VISrl_R rest vs SSp-n_L plan | 0.038 |
| SSp-tr_L plan vs SSp-n_R plan | 0.99 |
| SSp-tr_R plan vs SSp-n_R plan | 0.74 |
| SSp-ul_L plan vs SSp-n_R plan | 1 |
| SSp-ul_R plan vs SSp-n_R plan | 1 |
| SSp-un_L plan vs SSp-n_R plan | 1 |
| SSp-un_R plan vs SSp-n_R plan | 1 |
| VISa_L plan vs SSp-n_R plan | 1 |
| VISam_L plan vs SSp-n_R plan | 1 |
| VISam_R plan vs SSp-n_R plan | 1 |
| VISa_R plan vs SSp-n_R plan | 1 |
| VISp_L plan vs SSp-n_R plan | 1 |
| VISpm_L plan vs SSp-n_R plan | 1 |
| VISpm_R plan vs SSp-n_R plan | 1 |
| VISp_R plan vs SSp-n_R plan | 1 |
| VISrl_L plan vs SSp-n_R plan | 1 |
| VISrl_R plan vs SSp-n_R plan | 1 |
| CFA_L rest vs SSp-n_R plan | 0.13 |
| CFA_R rest vs SSp-n_R plan | 0.3 |
| MOp_L rest vs SSp-n_R plan | 0.062 |
| MOp_R rest vs SSp-n_R plan | 0.13 |
| MOs_L rest vs SSp-n_R plan | 0.025 |
| MOs_R rest vs SSp-n_R plan | 0.019 |
| RFA_L rest vs SSp-n_R plan | **0.005** |
| RFA_R rest vs SSp-n_R plan | **0.0021** |

| | |
|---|---|
| RSPagl_L rest vs SSp-n_R plan | **0.00039** |
| RSPagl_R rest vs SSp-n_R plan | **0.00095** |
| RSPd_L rest vs SSp-n_R plan | **0.00058** |
| RSPd_R rest vs SSp-n_R plan | **0.0008** |
| SSp-bfd_L rest vs SSp-n_R plan | 0.052 |
| SSp-bfd_R rest vs SSp-n_R plan | 0.24 |
| SSp-ll_L rest vs SSp-n_R plan | 0.017 |
| SSp-ll_R rest vs SSp-n_R plan | 0.16 |
| SSp-n_L rest vs SSp-n_R plan | 0.17 |
| SSp-n_R rest vs SSp-n_R plan | 0.63 |
| SSp-tr_L rest vs SSp-n_R plan | **0.0065** |
| SSp-tr_R rest vs SSp-n_R plan | 0.038 |
| SSp-ul_L rest vs SSp-n_R plan | 0.16 |
| SSp-ul_R rest vs SSp-n_R plan | 0.33 |
| SSp-un_L rest vs SSp-n_R plan | 0.21 |
| SSp-un_R rest vs SSp-n_R plan | 0.49 |
| VISa_L rest vs SSp-n_R plan | **0.0043** |
| VISam_L rest vs SSp-n_R plan | **0.00091** |
| VISam_R rest vs SSp-n_R plan | **0.0018** |
| VISa_R rest vs SSp-n_R plan | **0.0073** |
| VISp_L rest vs SSp-n_R plan | **0.00029** |
| VISpm_L rest vs SSp-n_R plan | **0.00011** |
| VISpm_R rest vs SSp-n_R plan | **0.00029** |
| VISp_R rest vs SSp-n_R plan | **0.00071** |
| VISrl_L rest vs SSp-n_R plan | **0.003** |
| VISrl_R rest vs SSp-n_R plan | **0.0081** |
| SSp-tr_R plan vs SSp-tr_L plan | 1 |
| SSp-ul_L plan vs SSp-tr_L plan | 1 |
| SSp-ul_R plan vs SSp-tr_L plan | 1 |
| SSp-un_L plan vs SSp-tr_L plan | 1 |
| SSp-un_R plan vs SSp-tr_L plan | 1 |
| VISa_L plan vs SSp-tr_L plan | 1 |
| VISam_L plan vs SSp-tr_L plan | 1 |
| VISam_R plan vs SSp-tr_L plan | 1 |
| VISa_R plan vs SSp-tr_L plan | 1 |
| VISp_L plan vs SSp-tr_L plan | 0.084 |
| VISpm_L plan vs SSp-tr_L plan | 0.67 |
| VISpm_R plan vs SSp-tr_L plan | 0.98 |
| VISp_R plan vs SSp-tr_L plan | 0.42 |
| VISrl_L plan vs SSp-tr_L plan | 0.24 |
| VISrl_R plan vs SSp-tr_L plan | 0.78 |
| CFA_L rest vs SSp-tr_L plan | **1.1e-07** |
| CFA_R rest vs SSp-tr_L plan | **8.2e-07** |
| MOp_L rest vs SSp-tr_L plan | **2.7e-08** |
| MOp_R rest vs SSp-tr_L plan | **1.2e-07** |
| MOs_L rest vs SSp-tr_L plan | **4.2e-09** |

| | |
|---|---|
| MOs_R rest vs SSp-tr_L plan | **2.2e-09** |
| RFA_L rest vs SSp-tr_L plan | **0** |
| RFA_R rest vs SSp-tr_L plan | **0** |
| RSPagl_L rest vs SSp-tr_L plan | **0** |
| RSPagl_R rest vs SSp-tr_L plan | **0** |
| RSPd_L rest vs SSp-tr_L plan | **0** |
| RSPd_R rest vs SSp-tr_L plan | **0** |
| SSp-bfd_L rest vs SSp-tr_L plan | **1.9e-08** |
| SSp-bfd_R rest vs SSp-tr_L plan | **4.6e-07** |
| SSp-ll_L rest vs SSp-tr_L plan | **1.8e-09** |
| SSp-ll_R rest vs SSp-tr_L plan | **1.8e-07** |
| SSp-n_L rest vs SSp-tr_L plan | **2.1e-07** |
| SSp-n_R rest vs SSp-tr_L plan | **8.4e-06** |
| SSp-tr_L rest vs SSp-tr_L plan | **0** |
| SSp-tr_R rest vs SSp-tr_L plan | **1e-08** |
| SSp-ul_L rest vs SSp-tr_L plan | **1.8e-07** |
| SSp-ul_R rest vs SSp-tr_L plan | **1.1e-06** |
| SSp-un_L rest vs SSp-tr_L plan | **3.6e-07** |
| SSp-un_R rest vs SSp-tr_L plan | **3.4e-06** |
| VISa_L rest vs SSp-tr_L plan | **0** |
| VISam_L rest vs SSp-tr_L plan | **0** |
| VISam_R rest vs SSp-tr_L plan | **0** |
| VISa_R rest vs SSp-tr_L plan | **0** |
| VISp_L rest vs SSp-tr_L plan | **0** |
| VISpm_L rest vs SSp-tr_L plan | **0** |
| VISpm_R rest vs SSp-tr_L plan | **0** |
| VISp_R rest vs SSp-tr_L plan | **0** |
| VISrl_L rest vs SSp-tr_L plan | **0** |
| VISrl_R rest vs SSp-tr_L plan | **0** |
| SSp-ul_L plan vs SSp-tr_R plan | 1 |
| SSp-ul_R plan vs SSp-tr_R plan | 1 |
| SSp-un_L plan vs SSp-tr_R plan | 1 |
| SSp-un_R plan vs SSp-tr_R plan | 1 |
| VISa_L plan vs SSp-tr_R plan | 1 |
| VISam_L plan vs SSp-tr_R plan | 1 |
| VISam_R plan vs SSp-tr_R plan | 1 |
| VISa_R plan vs SSp-tr_R plan | 1 |
| VISp_L plan vs SSp-tr_R plan | **0.0067** |
| VISpm_L plan vs SSp-tr_R plan | 0.17 |
| VISpm_R plan vs SSp-tr_R plan | 0.63 |
| VISp_R plan vs SSp-tr_R plan | 0.068 |
| VISrl_L plan vs SSp-tr_R plan | 0.029 |
| VISrl_R plan vs SSp-tr_R plan | 0.24 |
| CFA_L rest vs SSp-tr_R plan | **3.6e-10** |
| CFA_R rest vs SSp-tr_R plan | **1.1e-08** |
| MOp_L rest vs SSp-tr_R plan | **0** |

| | |
|---|---|
| MOp_R rest vs SSp-tr_R plan | **4.2e-10** |
| MOs_L rest vs SSp-tr_R plan | **0** |
| MOs_R rest vs SSp-tr_R plan | **0** |
| RFA_L rest vs SSp-tr_R plan | **0** |
| RFA_R rest vs SSp-tr_R plan | **0** |
| RSPagl_L rest vs SSp-tr_R plan | **0** |
| RSPagl_R rest vs SSp-tr_R plan | **0** |
| RSPd_L rest vs SSp-tr_R plan | **0** |
| RSPd_R rest vs SSp-tr_R plan | **0** |
| SSp-bfd_L rest vs SSp-tr_R plan | **0** |
| SSp-bfd_R rest vs SSp-tr_R plan | **5.4e-09** |
| SSp-ll_L rest vs SSp-tr_R plan | **0** |
| SSp-ll_R rest vs SSp-tr_R plan | **1.3e-09** |
| SSp-n_L rest vs SSp-tr_R plan | **1.7e-09** |
| SSp-n_R rest vs SSp-tr_R plan | **1.5e-07** |
| SSp-tr_L rest vs SSp-tr_R plan | **0** |
| SSp-tr_R rest vs SSp-tr_R plan | **0** |
| SSp-ul_L rest vs SSp-tr_R plan | **1.2e-09** |
| SSp-ul_R rest vs SSp-tr_R plan | **1.5e-08** |
| SSp-un_L rest vs SSp-tr_R plan | **3.9e-09** |
| SSp-un_R rest vs SSp-tr_R plan | **5.7e-08** |
| VISa_L rest vs SSp-tr_R plan | **0** |
| VISam_L rest vs SSp-tr_R plan | **0** |
| VISam_R rest vs SSp-tr_R plan | **0** |
| VISa_R rest vs SSp-tr_R plan | **0** |
| VISp_L rest vs SSp-tr_R plan | **0** |
| VISpm_L rest vs SSp-tr_R plan | **0** |
| VISpm_R rest vs SSp-tr_R plan | **0** |
| VISp_R rest vs SSp-tr_R plan | **0** |
| VISrl_L rest vs SSp-tr_R plan | **0** |
| VISrl_R rest vs SSp-tr_R plan | **0** |
| SSp-ul_R plan vs SSp-ul_L plan | 1 |
| SSp-un_L plan vs SSp-ul_L plan | 1 |
| SSp-un_R plan vs SSp-ul_L plan | 1 |
| VISa_L plan vs SSp-ul_L plan | 1 |
| VISam_L plan vs SSp-ul_L plan | 1 |
| VISam_R plan vs SSp-ul_L plan | 1 |
| VISa_R plan vs SSp-ul_L plan | 1 |
| VISp_L plan vs SSp-ul_L plan | 0.51 |
| VISpm_L plan vs SSp-ul_L plan | 0.99 |
| VISpm_R plan vs SSp-ul_L plan | 1 |
| VISp_R plan vs SSp-ul_L plan | 0.94 |
| VISrl_L plan vs SSp-ul_L plan | 0.82 |
| VISrl_R plan vs SSp-ul_L plan | 1 |
| CFA_L rest vs SSp-ul_L plan | **8.2e-06** |
| CFA_R rest vs SSp-ul_L plan | **4.8e-05** |

| | |
|---|---|
| MOp_L rest vs SSp-ul_L plan | **2.3e-06** |
| MOp_R rest vs SSp-ul_L plan | **8.6e-06** |
| MOs_L rest vs SSp-ul_L plan | **5.1e-07** |
| MOs_R rest vs SSp-ul_L plan | **3.3e-07** |
| RFA_L rest vs SSp-ul_L plan | **4.3e-08** |
| RFA_R rest vs SSp-ul_L plan | **1.1e-08** |
| RSPagl_L rest vs SSp-ul_L plan | **1e-10** |
| RSPagl_R rest vs SSp-ul_L plan | **2.9e-09** |
| RSPd_L rest vs SSp-ul_L plan | **9.5e-10** |
| RSPd_R rest vs SSp-ul_L plan | **2.1e-09** |
| SSp-bfd_L rest vs SSp-ul_L plan | **1.7e-06** |
| SSp-bfd_R rest vs SSp-ul_L plan | **2.9e-05** |
| SSp-ll_L rest vs SSp-ul_L plan | **2.9e-07** |
| SSp-ll_R rest vs SSp-ul_L plan | **1.2e-05** |
| SSp-n_L rest vs SSp-ul_L plan | **1.4e-05** |
| SSp-n_R rest vs SSp-ul_L plan | **0.00038** |
| SSp-tr_L rest vs SSp-ul_L plan | **6.4e-08** |
| SSp-tr_R rest vs SSp-ul_L plan | **1e-06** |
| SSp-ul_L rest vs SSp-ul_L plan | **1.2e-05** |
| SSp-ul_R rest vs SSp-ul_L plan | **6.2e-05** |
| SSp-un_L rest vs SSp-ul_L plan | **2.3e-05** |
| SSp-un_R rest vs SSp-ul_L plan | **0.00017** |
| VISa_L rest vs SSp-ul_L plan | **3.4e-08** |
| VISam_L rest vs SSp-ul_L plan | **2.7e-09** |
| VISam_R rest vs SSp-ul_L plan | **8.6e-09** |
| VISa_R rest vs SSp-ul_L plan | **7.6e-08** |
| VISp_L rest vs SSp-ul_L plan | **0** |
| VISpm_L rest vs SSp-ul_L plan | **0** |
| VISpm_R rest vs SSp-ul_L plan | **0** |
| VISp_R rest vs SSp-ul_L plan | **1.6e-09** |
| VISrl_L rest vs SSp-ul_L plan | **2e-08** |
| VISrl_R rest vs SSp-ul_L plan | **8.8e-08** |
| SSp-un_L plan vs SSp-ul_R plan | 1 |
| SSp-un_R plan vs SSp-ul_R plan | 1 |
| VISa_L plan vs SSp-ul_R plan | 1 |
| VISam_L plan vs SSp-ul_R plan | 1 |
| VISam_R plan vs SSp-ul_R plan | 1 |
| VISa_R plan vs SSp-ul_R plan | 1 |
| VISp_L plan vs SSp-ul_R plan | 0.23 |
| VISpm_L plan vs SSp-ul_R plan | 0.91 |
| VISpm_R plan vs SSp-ul_R plan | 1 |
| VISp_R plan vs SSp-ul_R plan | 0.72 |
| VISrl_L plan vs SSp-ul_R plan | 0.52 |
| VISrl_R plan vs SSp-ul_R plan | 0.96 |
| CFA_L rest vs SSp-ul_R plan | **9.9e-07** |
| CFA_R rest vs SSp-ul_R plan | **6.4e-06** |

| | |
|---|---|
| MOp_L rest vs SSp-ul_R plan | **2.6e-07** |
| MOp_R rest vs SSp-ul_R plan | **1e-06** |
| MOs_L rest vs SSp-ul_R plan | **5.1e-08** |
| MOs_R rest vs SSp-ul_R plan | **3.2e-08** |
| RFA_L rest vs SSp-ul_R plan | **3e-09** |
| RFA_R rest vs SSp-ul_R plan | **1.5e-12** |
| RSPagl_L rest vs SSp-ul_R plan | **0** |
| RSPagl_R rest vs SSp-ul_R plan | **0** |
| RSPd_L rest vs SSp-ul_R plan | **0** |
| RSPd_R rest vs SSp-ul_R plan | **0** |
| SSp-bfd_L rest vs SSp-ul_R plan | **1.9e-07** |
| SSp-bfd_R rest vs SSp-ul_R plan | **3.8e-06** |
| SSp-ll_L rest vs SSp-ul_R plan | **2.8e-08** |
| SSp-ll_R rest vs SSp-ul_R plan | **1.5e-06** |
| SSp-n_L rest vs SSp-ul_R plan | **1.8e-06** |
| SSp-n_R rest vs SSp-ul_R plan | **5.8e-05** |
| SSp-tr_L rest vs SSp-ul_R plan | **5e-09** |
| SSp-tr_R rest vs SSp-ul_R plan | **1.1e-07** |
| SSp-ul_L rest vs SSp-ul_R plan | **1.5e-06** |
| SSp-ul_R rest vs SSp-ul_R plan | **8.3e-06** |
| SSp-un_L rest vs SSp-ul_R plan | **3e-06** |
| SSp-un_R rest vs SSp-ul_R plan | **2.5e-05** |
| VISa_L rest vs SSp-ul_R plan | **2.1e-09** |
| VISam_L rest vs SSp-ul_R plan | **0** |
| VISam_R rest vs SSp-ul_R plan | **0** |
| VISa_R rest vs SSp-ul_R plan | **6.2e-09** |
| VISp_L rest vs SSp-ul_R plan | **0** |
| VISpm_L rest vs SSp-ul_R plan | **0** |
| VISpm_R rest vs SSp-ul_R plan | **0** |
| VISp_R rest vs SSp-ul_R plan | **0** |
| VISrl_L rest vs SSp-ul_R plan | **8.1e-10** |
| VISrl_R rest vs SSp-ul_R plan | **7.4e-09** |
| SSp-un_R plan vs SSp-un_L plan | 1 |
| VISa_L plan vs SSp-un_L plan | 1 |
| VISam_L plan vs SSp-un_L plan | 1 |
| VISam_R plan vs SSp-un_L plan | 1 |
| VISa_R plan vs SSp-un_L plan | 1 |
| VISp_L plan vs SSp-un_L plan | 1 |
| VISpm_L plan vs SSp-un_L plan | 1 |
| VISpm_R plan vs SSp-un_L plan | 1 |
| VISp_R plan vs SSp-un_L plan | 1 |
| VISrl_L plan vs SSp-un_L plan | 1 |
| VISrl_R plan vs SSp-un_L plan | 1 |
| CFA_L rest vs SSp-un_L plan | **0.0021** |
| CFA_R rest vs SSp-un_L plan | **0.0086** |
| MOp_L rest vs SSp-un_L plan | **0.00076** |

| | |
|---|---|
| MOp_R rest vs SSp-un_L plan | **0.0022** |
| MOs_L rest vs SSp-un_L plan | **0.00022** |
| MOs_R rest vs SSp-un_L plan | **0.00015** |
| RFA_L rest vs SSp-un_L plan | **2.7e-05** |
| RFA_R rest vs SSp-un_L plan | **9.1e-06** |
| RSPagl_L rest vs SSp-un_L plan | **1.2e-06** |
| RSPagl_R rest vs SSp-un_L plan | **3.5e-06** |
| RSPd_L rest vs SSp-un_L plan | **1.9e-06** |
| RSPd_R rest vs SSp-un_L plan | **2.8e-06** |
| SSp-bfd_L rest vs SSp-un_L plan | **0.00059** |
| SSp-bfd_R rest vs SSp-un_L plan | **0.0058** |
| SSp-ll_L rest vs SSp-un_L plan | **0.00014** |
| SSp-ll_R rest vs SSp-un_L plan | **0.003** |
| SSp-n_L rest vs SSp-un_L plan | **0.0033** |
| SSp-n_R rest vs SSp-un_L plan | 0.041 |
| SSp-tr_L rest vs SSp-un_L plan | **3.8e-05** |
| SSp-tr_R rest vs SSp-un_L plan | **0.00038** |
| SSp-ul_L rest vs SSp-un_L plan | **0.0029** |
| SSp-ul_R rest vs SSp-un_L plan | 0.01 |
| SSp-un_L rest vs SSp-un_L plan | **0.0049** |
| SSp-un_R rest vs SSp-un_L plan | 0.023 |
| VISa_L rest vs SSp-un_L plan | **2.2e-05** |
| VISam_L rest vs SSp-un_L plan | **3.3e-06** |
| VISam_R rest vs SSp-un_L plan | **7.5e-06** |
| VISa_R rest vs SSp-un_L plan | **4.4e-05** |
| VISp_L rest vs SSp-un_L plan | **8.4e-07** |
| VISpm_L rest vs SSp-un_L plan | **2.5e-07** |
| VISpm_R rest vs SSp-un_L plan | **8.3e-07** |
| VISp_R rest vs SSp-un_L plan | **2.4e-06** |
| VISrl_L rest vs SSp-un_L plan | **1.5e-05** |
| VISrl_R rest vs SSp-un_L plan | **5e-05** |
| VISa_L plan vs SSp-un_R plan | 1 |
| VISam_L plan vs SSp-un_R plan | 1 |
| VISam_R plan vs SSp-un_R plan | 1 |
| VISa_R plan vs SSp-un_R plan | 1 |
| VISp_L plan vs SSp-un_R plan | 0.93 |
| VISpm_L plan vs SSp-un_R plan | 1 |
| VISpm_R plan vs SSp-un_R plan | 1 |
| VISp_R plan vs SSp-un_R plan | 1 |
| VISrl_L plan vs SSp-un_R plan | 1 |
| VISrl_R plan vs SSp-un_R plan | 1 |
| CFA_L rest vs SSp-un_R plan | **0.0002** |
| CFA_R rest vs SSp-un_R plan | **0.00097** |
| MOp_L rest vs SSp-un_R plan | **6.4e-05** |
| MOp_R rest vs SSp-un_R plan | **0.00021** |
| MOs_L rest vs SSp-un_R plan | **1.6e-05** |

| | |
|---|---|
| MOs_R rest vs SSp-un_R plan | **1.1e-05** |
| RFA_L rest vs SSp-un_R plan | **1.7e-06** |
| RFA_R rest vs SSp-un_R plan | **5.2e-07** |
| RSPagl_L rest vs SSp-un_R plan | **5.7e-08** |
| RSPagl_R rest vs SSp-un_R plan | **1.8e-07** |
| RSPd_L rest vs SSp-un_R plan | **9.7e-08** |
| RSPd_R rest vs SSp-un_R plan | **1.5e-07** |
| SSp-bfd_L rest vs SSp-un_R plan | **4.9e-05** |
| SSp-bfd_R rest vs SSp-un_R plan | **0.00062** |
| SSp-ll_L rest vs SSp-un_R plan | **9.7e-06** |
| SSp-ll_R rest vs SSp-un_R plan | **0.00029** |
| SSp-n_L rest vs SSp-un_R plan | **0.00033** |
| SSp-n_R rest vs SSp-un_R plan | **0.0059** |
| SSp-tr_L rest vs SSp-un_R plan | **2.4e-06** |
| SSp-tr_R rest vs SSp-un_R plan | **3e-05** |
| SSp-ul_L rest vs SSp-un_R plan | **0.00029** |
| SSp-ul_R rest vs SSp-un_R plan | **0.0012** |
| SSp-un_L rest vs SSp-un_R plan | **0.00051** |
| SSp-un_R rest vs SSp-un_R plan | **0.003** |
| VISa_L rest vs SSp-un_R plan | **1.4e-06** |
| VISam_L rest vs SSp-un_R plan | **1.7e-07** |
| VISam_R rest vs SSp-un_R plan | **4.2e-07** |
| VISa_R rest vs SSp-un_R plan | **2.9e-06** |
| VISp_L rest vs SSp-un_R plan | **3.9e-08** |
| VISpm_L rest vs SSp-un_R plan | **1e-08** |
| VISpm_R rest vs SSp-un_R plan | **3.9e-08** |
| VISp_R rest vs SSp-un_R plan | **1.2e-07** |
| VISrl_L rest vs SSp-un_R plan | **8.6e-07** |
| VISrl_R rest vs SSp-un_R plan | **3.3e-06** |
| VISam_L plan vs VISa_L plan | 1 |
| VISam_R plan vs VISa_L plan | 1 |
| VISa_R plan vs VISa_L plan | 1 |
| VISp_L plan vs VISa_L plan | 0.97 |
| VISpm_L plan vs VISa_L plan | 1 |
| VISpm_R plan vs VISa_L plan | 1 |
| VISp_R plan vs VISa_L plan | 1 |
| VISrl_L plan vs VISa_L plan | 1 |
| VISrl_R plan vs VISa_L plan | 1 |
| CFA_L rest vs VISa_L plan | **0.00038** |
| CFA_R rest vs VISa_L plan | **0.0017** |
| MOp_L rest vs VISa_L plan | **0.00012** |
| MOp_R rest vs VISa_L plan | **0.00039** |
| MOs_L rest vs VISa_L plan | **3.2e-05** |
| MOs_R rest vs VISa_L plan | **2.1e-05** |
| RFA_L rest vs VISa_L plan | **3.5e-06** |
| RFA_R rest vs VISa_L plan | **1.1e-06** |

| | |
|---|---|
| RSPagl_L rest vs VISa_L plan | **1.3e-07** |
| RSPagl_R rest vs VISa_L plan | **4e-07** |
| RSPd_L rest vs VISa_L plan | **2.1e-07** |
| RSPd_R rest vs VISa_L plan | **3.2e-07** |
| SSp-bfd_L rest vs VISa_L plan | **9.4e-05** |
| SSp-bfd_R rest vs VISa_L plan | **0.0011** |
| SSp-ll_L rest vs VISa_L plan | **1.9e-05** |
| SSp-ll_R rest vs VISa_L plan | **0.00054** |
| SSp-n_L rest vs VISa_L plan | **0.00061** |
| SSp-n_R rest vs VISa_L plan | 0.01 |
| SSp-tr_L rest vs VISa_L plan | **5e-06** |
| SSp-tr_R rest vs VISa_L plan | **5.8e-05** |
| SSp-ul_L rest vs VISa_L plan | **0.00053** |
| SSp-ul_R rest vs VISa_L plan | **0.0022** |
| SSp-un_L rest vs VISa_L plan | **0.00093** |
| SSp-un_R rest vs VISa_L plan | **0.0051** |
| VISa_L rest vs VISa_L plan | **2.9e-06** |
| VISam_L rest vs VISa_L plan | **3.8e-07** |
| VISam_R rest vs VISa_L plan | **9e-07** |
| VISa_R rest vs VISa_L plan | **5.9e-06** |
| VISp_L rest vs VISa_L plan | **8.8e-08** |
| VISpm_L rest vs VISa_L plan | **2.4e-08** |
| VISpm_R rest vs VISa_L plan | **8.7e-08** |
| VISp_R rest vs VISa_L plan | **2.7e-07** |
| VISrl_L rest vs VISa_L plan | **1.8e-06** |
| VISrl_R rest vs VISa_L plan | **6.7e-06** |
| VISam_R plan vs VISam_L plan | 1 |
| VISa_R plan vs VISam_L plan | 1 |
| VISp_L plan vs VISam_L plan | 1 |
| VISpm_L plan vs VISam_L plan | 1 |
| VISpm_R plan vs VISam_L plan | 1 |
| VISp_R plan vs VISam_L plan | 1 |
| VISrl_L plan vs VISam_L plan | 1 |
| VISrl_R plan vs VISam_L plan | 1 |
| CFA_L rest vs VISam_L plan | **0.0071** |
| CFA_R rest vs VISam_L plan | 0.026 |
| MOp_L rest vs VISam_L plan | **0.0027** |
| MOp_R rest vs VISam_L plan | **0.0073** |
| MOs_L rest vs VISam_L plan | **0.00083** |
| MOs_R rest vs VISam_L plan | **0.00059** |
| RFA_L rest vs VISam_L plan | **0.00012** |
| RFA_R rest vs VISam_L plan | **4.1e-05** |
| RSPagl_L rest vs VISam_L plan | **5.8e-06** |
| RSPagl_R rest vs VISam_L plan | **1.6e-05** |
| RSPd_L rest vs VISam_L plan | **9.3e-06** |
| RSPd_R rest vs VISam_L plan | **1.4e-05** |

| | |
|---|---|
| SSp-bfd_L rest vs VISam_L plan | **0.0021** |
| SSp-bfd_R rest vs VISam_L plan | 0.018 |
| SSp-ll_L rest vs VISam_L plan | **0.00053** |
| SSp-ll_R rest vs VISam_L plan | **0.0096** |
| SSp-n_L rest vs VISam_L plan | 0.011 |
| SSp-n_R rest vs VISam_L plan | 0.1 |
| SSp-tr_L rest vs VISam_L plan | **0.00016** |
| SSp-tr_R rest vs VISam_L plan | **0.0014** |
| SSp-ul_L rest vs VISam_L plan | **0.0095** |
| SSp-ul_R rest vs VISam_L plan | 0.03 |
| SSp-un_L rest vs VISam_L plan | 0.015 |
| SSp-un_R rest vs VISam_L plan | 0.061 |
| VISa_L rest vs VISam_L plan | **9.7e-05** |
| VISam_L rest vs VISam_L plan | **1.6e-05** |
| VISam_R rest vs VISam_L plan | **3.4e-05** |
| VISa_R rest vs VISam_L plan | **0.00018** |
| VISp_L rest vs VISam_L plan | **4.2e-06** |
| VISpm_L rest vs VISam_L plan | **1.3e-06** |
| VISpm_R rest vs VISam_L plan | **4.2e-06** |
| VISp_R rest vs VISam_L plan | **1.2e-05** |
| VISrl_L rest vs VISam_L plan | **6.4e-05** |
| VISrl_R rest vs VISam_L plan | **0.00021** |
| VISa_R plan vs VISam_R plan | 1 |
| VISp_L plan vs VISam_R plan | 1 |
| VISpm_L plan vs VISam_R plan | 1 |
| VISpm_R plan vs VISam_R plan | 1 |
| VISp_R plan vs VISam_R plan | 1 |
| VISrl_L plan vs VISam_R plan | 1 |
| VISrl_R plan vs VISam_R plan | 1 |
| CFA_L rest vs VISam_R plan | **0.0013** |
| CFA_R rest vs VISam_R plan | **0.0053** |
| MOp_L rest vs VISam_R plan | **0.00044** |
| MOp_R rest vs VISam_R plan | **0.0013** |
| MOs_L rest vs VISam_R plan | **0.00012** |
| MOs_R rest vs VISam_R plan | **8.2e-05** |
| RFA_L rest vs VISam_R plan | **1.5e-05** |
| RFA_R rest vs VISam_R plan | **4.7e-06** |
| RSPagl_L rest vs VISam_R plan | **5.9e-07** |
| RSPagl_R rest vs VISam_R plan | **1.8e-06** |
| RSPd_L rest vs VISam_R plan | **9.8e-07** |
| RSPd_R rest vs VISam_R plan | **1.5e-06** |
| SSp-bfd_L rest vs VISam_R plan | **0.00034** |
| SSp-bfd_R rest vs VISam_R plan | **0.0035** |
| SSp-ll_L rest vs VISam_R plan | **7.5e-05** |
| SSp-ll_R rest vs VISam_R plan | **0.0018** |
| SSp-n_L rest vs VISam_R plan | **0.002** |

| | |
|---|---|
| SSp-n_R rest vs VISam_R plan | 0.027 |
| SSp-tr_L rest vs VISam_R plan | **2e-05** |
| SSp-tr_R rest vs VISam_R plan | **0.00021** |
| SSp-ul_L rest vs VISam_R plan | **0.0018** |
| SSp-ul_R rest vs VISam_R plan | **0.0065** |
| SSp-un_L rest vs VISam_R plan | **0.0029** |
| SSp-un_R rest vs VISam_R plan | 0.015 |
| VISa_L rest vs VISam_R plan | **1.2e-05** |
| VISam_L rest vs VISam_R plan | **1.7e-06** |
| VISam_R rest vs VISam_R plan | **3.9e-06** |
| VISa_R rest vs VISam_R plan | **2.4e-05** |
| VISp_L rest vs VISam_R plan | **4.2e-07** |
| VISpm_L rest vs VISam_R plan | **1.2e-07** |
| VISpm_R rest vs VISam_R plan | **4.1e-07** |
| VISp_R rest vs VISam_R plan | **1.2e-06** |
| VISrl_L rest vs VISam_R plan | **7.7e-06** |
| VISrl_R rest vs VISam_R plan | **2.7e-05** |
| VISp_L plan vs VISa_R plan | 0.8 |
| VISpm_L plan vs VISa_R plan | 1 |
| VISpm_R plan vs VISa_R plan | 1 |
| VISp_R plan vs VISa_R plan | 0.99 |
| VISrl_L plan vs VISa_R plan | 0.97 |
| VISrl_R plan vs VISa_R plan | 1 |
| CFA_L rest vs VISa_R plan | **5.4e-05** |
| CFA_R rest vs VISa_R plan | **0.00028** |
| MOp_L rest vs VISa_R plan | **1.6e-05** |
| MOp_R rest vs VISa_R plan | **5.5e-05** |
| MOs_L rest vs VISa_R plan | **3.8e-06** |
| MOs_R rest vs VISa_R plan | **2.5e-06** |
| RFA_L rest vs VISa_R plan | **3.7e-07** |
| RFA_R rest vs VISa_R plan | **1.1e-07** |
| RSPagl_L rest vs VISa_R plan | **1e-08** |
| RSPagl_R rest vs VISa_R plan | **3.6e-08** |
| RSPd_L rest vs VISa_R plan | **1.8e-08** |
| RSPd_R rest vs VISa_R plan | **2.8e-08** |
| SSp-bfd_L rest vs VISa_R plan | **1.2e-05** |
| SSp-bfd_R rest vs VISa_R plan | **0.00018** |
| SSp-ll_L rest vs VISa_R plan | **2.2e-06** |
| SSp-ll_R rest vs VISa_R plan | **7.9e-05** |
| SSp-n_L rest vs VISa_R plan | **9e-05** |
| SSp-n_R rest vs VISa_R plan | **0.0019** |
| SSp-tr_L rest vs VISa_R plan | **5.4e-07** |
| SSp-tr_R rest vs VISa_R plan | **7.3e-06** |
| SSp-ul_L rest vs VISa_R plan | **7.8e-05** |
| SSp-ul_R rest vs VISa_R plan | **0.00035** |
| SSp-un_L rest vs VISa_R plan | **0.00014** |

| | |
|---|---|
| SSp-un_R rest vs VISa_R plan | **0.00092** |
| VISa_L rest vs VISa_R plan | **2.9e-07** |
| VISam_L rest vs VISa_R plan | **3.4e-08** |
| VISam_R rest vs VISa_R plan | **8.6e-08** |
| VISa_R rest vs VISa_R plan | **6.3e-07** |
| VISp_L rest vs VISa_R plan | **6.6e-09** |
| VISpm_L rest vs VISa_R plan | **1e-09** |
| VISpm_R rest vs VISa_R plan | **6.5e-09** |
| VISp_R rest vs VISa_R plan | **2.4e-08** |
| VISrl_L rest vs VISa_R plan | **1.8e-07** |
| VISrl_R rest vs VISa_R plan | **7.3e-07** |
| VISpm_L plan vs VISp_L plan | 1 |
| VISpm_R plan vs VISp_L plan | 1 |
| VISp_R plan vs VISp_L plan | 1 |
| VISrl_L plan vs VISp_L plan | 1 |
| VISrl_R plan vs VISp_L plan | 1 |
| CFA_L rest vs VISp_L plan | 1 |
| CFA_R rest vs VISp_L plan | 1 |
| MOp_L rest vs VISp_L plan | 0.99 |
| MOp_R rest vs VISp_L plan | 1 |
| MOs_L rest vs VISp_L plan | 0.93 |
| MOs_R rest vs VISp_L plan | 0.9 |
| RFA_L rest vs VISp_L plan | 0.69 |
| RFA_R rest vs VISp_L plan | 0.51 |
| RSPagl_L rest vs VISp_L plan | 0.24 |
| RSPagl_R rest vs VISp_L plan | 0.37 |
| RSPd_L rest vs VISp_L plan | 0.29 |
| RSPd_R rest vs VISp_L plan | 0.34 |
| SSp-bfd_L rest vs VISp_L plan | 0.98 |
| SSp-bfd_R rest vs VISp_L plan | 1 |
| SSp-ll_L rest vs VISp_L plan | 0.89 |
| SSp-ll_R rest vs VISp_L plan | 1 |
| SSp-n_L rest vs VISp_L plan | 1 |
| SSp-n_R rest vs VISp_L plan | 1 |
| SSp-tr_L rest vs VISp_L plan | 0.74 |
| SSp-tr_R rest vs VISp_L plan | 0.96 |
| SSp-ul_L rest vs VISp_L plan | 1 |
| SSp-ul_R rest vs VISp_L plan | 1 |
| SSp-un_L rest vs VISp_L plan | 1 |
| SSp-un_R rest vs VISp_L plan | 1 |
| VISa_L rest vs VISp_L plan | 0.65 |
| VISam_L rest vs VISp_L plan | 0.36 |
| VISam_R rest vs VISp_L plan | 0.48 |
| VISa_R rest vs VISp_L plan | 0.76 |
| VISp_L rest vs VISp_L plan | 0.21 |
| VISpm_L rest vs VISp_L plan | 0.12 |

| | |
|---|---|
| VISpm_R rest vs VISp_L plan | 0.21 |
| VISp_R rest vs VISp_L plan | 0.32 |
| VISrl_L rest vs VISp_L plan | 0.59 |
| VISrl_R rest vs VISp_L plan | 0.78 |
| VISpm_R plan vs VISpm_L plan | 1 |
| VISp_R plan vs VISpm_L plan | 1 |
| VISrl_L plan vs VISpm_L plan | 1 |
| VISrl_R plan vs VISpm_L plan | 1 |
| CFA_L rest vs VISpm_L plan | 0.66 |
| CFA_R rest vs VISpm_L plan | 0.89 |
| MOp_L rest vs VISpm_L plan | 0.47 |
| MOp_R rest vs VISpm_L plan | 0.67 |
| MOs_L rest vs VISpm_L plan | 0.27 |
| MOs_R rest vs VISpm_L plan | 0.23 |
| RFA_L rest vs VISpm_L plan | 0.089 |
| RFA_R rest vs VISpm_L plan | 0.045 |
| RSPagl_L rest vs VISpm_L plan | 0.012 |
| RSPagl_R rest vs VISpm_L plan | 0.024 |
| RSPd_L rest vs VISpm_L plan | 0.016 |
| RSPd_R rest vs VISpm_L plan | 0.021 |
| SSp-bfd_L rest vs VISpm_L plan | 0.42 |
| SSp-bfd_R rest vs VISpm_L plan | 0.84 |
| SSp-ll_L rest vs VISpm_L plan | 0.21 |
| SSp-ll_R rest vs VISpm_L plan | 0.72 |
| SSp-n_L rest vs VISpm_L plan | 0.74 |
| SSp-n_R rest vs VISpm_L plan | 0.99 |
| SSp-tr_L rest vs VISpm_L plan | 0.11 |
| SSp-tr_R rest vs VISpm_L plan | 0.35 |
| SSp-ul_L rest vs VISpm_L plan | 0.72 |
| SSp-ul_R rest vs VISpm_L plan | 0.91 |
| SSp-un_L rest vs VISpm_L plan | 0.81 |
| SSp-un_R rest vs VISpm_L plan | 0.97 |
| VISa_L rest vs VISpm_L plan | 0.078 |
| VISam_L rest vs VISpm_L plan | 0.024 |
| VISam_R rest vs VISpm_L plan | 0.04 |
| VISa_R rest vs VISpm_L plan | 0.12 |
| VISp_L rest vs VISpm_L plan | **0.0093** |
| VISpm_L rest vs VISpm_L plan | **0.004** |
| VISpm_R rest vs VISpm_L plan | **0.0092** |
| VISp_R rest vs VISpm_L plan | 0.019 |
| VISrl_L rest vs VISpm_L plan | 0.061 |
| VISrl_R rest vs VISpm_L plan | 0.13 |
| VISp_R plan vs VISpm_R plan | 1 |
| VISrl_L plan vs VISpm_R plan | 1 |
| VISrl_R plan vs VISpm_R plan | 1 |
| CFA_L rest vs VISpm_R plan | 0.18 |

| | |
|---|---|
| CFA_R rest vs VISpm_R plan | 0.4 |
| MOp_L rest vs VISpm_R plan | 0.095 |
| MOp_R rest vs VISpm_R plan | 0.19 |
| MOs_L rest vs VISpm_R plan | 0.04 |
| MOs_R rest vs VISpm_R plan | 0.031 |
| RFA_L rest vs VISpm_R plan | **0.0087** |
| RFA_R rest vs VISpm_R plan | **0.0037** |
| RSPagl_L rest vs VISpm_R plan | **0.00073** |
| RSPagl_R rest vs VISpm_R plan | **0.0017** |
| RSPd_L rest vs VISpm_R plan | **0.0011** |
| RSPd_R rest vs VISpm_R plan | **0.0015** |
| SSp-bfd_L rest vs VISpm_R plan | 0.08 |
| SSp-bfd_R rest vs VISpm_R plan | 0.32 |
| SSp-ll_L rest vs VISpm_R plan | 0.028 |
| SSp-ll_R rest vs VISpm_R plan | 0.22 |
| SSp-n_L rest vs VISpm_R plan | 0.24 |
| SSp-n_R rest vs VISpm_R plan | 0.74 |
| SSp-tr_L rest vs VISpm_R plan | 0.011 |
| SSp-tr_R rest vs VISpm_R plan | 0.059 |
| SSp-ul_L rest vs VISpm_R plan | 0.22 |
| SSp-ul_R rest vs VISpm_R plan | 0.43 |
| SSp-un_L rest vs VISpm_R plan | 0.29 |
| SSp-un_R rest vs VISpm_R plan | 0.61 |
| VISa_L rest vs VISpm_R plan | **0.0074** |
| VISam_L rest vs VISpm_R plan | **0.0017** |
| VISam_R rest vs VISpm_R plan | **0.0032** |
| VISa_R rest vs VISpm_R plan | 0.012 |
| VISp_L rest vs VISpm_R plan | **0.00055** |
| VISpm_L rest vs VISpm_R plan | **0.00021** |
| VISpm_R rest vs VISpm_R plan | **0.00055** |
| VISp_R rest vs VISpm_R plan | **0.0013** |
| VISrl_L rest vs VISpm_R plan | **0.0053** |
| VISrl_R rest vs VISpm_R plan | 0.014 |
| VISrl_L plan vs VISp_R plan | 1 |
| VISrl_R plan vs VISp_R plan | 1 |
| CFA_L rest vs VISp_R plan | 0.87 |
| CFA_R rest vs VISp_R plan | 0.98 |
| MOp_L rest vs VISp_R plan | 0.72 |
| MOp_R rest vs VISp_R plan | 0.88 |
| MOs_L rest vs VISp_R plan | 0.5 |
| MOs_R rest vs VISp_R plan | 0.43 |
| RFA_L rest vs VISp_R plan | 0.21 |
| RFA_R rest vs VISp_R plan | 0.12 |
| RSPagl_L rest vs VISp_R plan | 0.036 |
| RSPagl_R rest vs VISp_R plan | 0.068 |
| RSPd_L rest vs VISp_R plan | 0.048 |

| | |
|---|---|
| RSPd_R rest vs VISp_R plan | 0.061 |
| SSp-bfd_L rest vs VISp_R plan | 0.68 |
| SSp-bfd_R rest vs VISp_R plan | 0.96 |
| SSp-ll_L rest vs VISp_R plan | 0.42 |
| SSp-ll_R rest vs VISp_R plan | 0.91 |
| SSp-n_L rest vs VISp_R plan | 0.92 |
| SSp-n_R rest vs VISp_R plan | 1 |
| SSp-tr_L rest vs VISp_R plan | 0.24 |
| SSp-tr_R rest vs VISp_R plan | 0.6 |
| SSp-ul_L rest vs VISp_R plan | 0.91 |
| SSp-ul_R rest vs VISp_R plan | 0.99 |
| SSp-un_L rest vs VISp_R plan | 0.95 |
| SSp-un_R rest vs VISp_R plan | 1 |
| VISa_L rest vs VISp_R plan | 0.19 |
| VISam_L rest vs VISp_R plan | 0.067 |
| VISam_R rest vs VISp_R plan | 0.11 |
| VISa_R rest vs VISp_R plan | 0.26 |
| VISp_L rest vs VISp_R plan | 0.029 |
| VISpm_L rest vs VISp_R plan | 0.014 |
| VISpm_R rest vs VISp_R plan | 0.029 |
| VISp_R rest vs VISp_R plan | 0.055 |
| VISrl_L rest vs VISp_R plan | 0.15 |
| VISrl_R rest vs VISp_R plan | 0.28 |
| VISrl_R plan vs VISrl_L plan | 1 |
| CFA_L rest vs VISrl_L plan | 0.96 |
| CFA_R rest vs VISrl_L plan | 1 |
| MOp_L rest vs VISrl_L plan | 0.88 |
| MOp_R rest vs VISrl_L plan | 0.96 |
| MOs_L rest vs VISrl_L plan | 0.71 |
| MOs_R rest vs VISrl_L plan | 0.64 |
| RFA_L rest vs VISrl_L plan | 0.37 |
| RFA_R rest vs VISrl_L plan | 0.23 |
| RSPagl_L rest vs VISrl_L plan | 0.081 |
| RSPagl_R rest vs VISrl_L plan | 0.14 |
| RSPd_L rest vs VISrl_L plan | 0.11 |
| RSPd_R rest vs VISrl_L plan | 0.13 |
| SSp-bfd_L rest vs VISrl_L plan | 0.85 |
| SSp-bfd_R rest vs VISrl_L plan | 0.99 |
| SSp-ll_L rest vs VISrl_L plan | 0.63 |
| SSp-ll_R rest vs VISrl_L plan | 0.98 |
| SSp-n_L rest vs VISrl_L plan | 0.98 |
| SSp-n_R rest vs VISrl_L plan | 1 |
| SSp-tr_L rest vs VISrl_L plan | 0.42 |
| SSp-tr_R rest vs VISrl_L plan | 0.79 |
| SSp-ul_L rest vs VISrl_L plan | 0.98 |
| SSp-ul_R rest vs VISrl_L plan | 1 |

| | |
|---|---|
| SSp-un_L rest vs VISrl_L plan | 0.99 |
| SSp-un_R rest vs VISrl_L plan | 1 |
| VISa_L rest vs VISrl_L plan | 0.34 |
| VISam_L rest vs VISrl_L plan | 0.14 |
| VISam_R rest vs VISrl_L plan | 0.21 |
| VISa_R rest vs VISrl_L plan | 0.44 |
| VISp_L rest vs VISrl_L plan | 0.067 |
| VISpm_L rest vs VISrl_L plan | 0.034 |
| VISpm_R rest vs VISrl_L plan | 0.067 |
| VISp_R rest vs VISrl_L plan | 0.12 |
| VISrl_L rest vs VISrl_L plan | 0.28 |
| VISrl_R rest vs VISrl_L plan | 0.46 |
| CFA_L rest vs VISrl_R plan | 0.55 |
| CFA_R rest vs VISrl_R plan | 0.81 |
| MOp_L rest vs VISrl_R plan | 0.36 |
| MOp_R rest vs VISrl_R plan | 0.55 |
| MOs_L rest vs VISrl_R plan | 0.19 |
| MOs_R rest vs VISrl_R plan | 0.16 |
| RFA_L rest vs VISrl_R plan | 0.057 |
| RFA_R rest vs VISrl_R plan | 0.028 |
| RSPagl_L rest vs VISrl_R plan | **0.0068** |
| RSPagl_R rest vs VISrl_R plan | 0.015 |
| RSPd_L rest vs VISrl_R plan | **0.0097** |
| RSPd_R rest vs VISrl_R plan | 0.013 |
| SSp-bfd_L rest vs VISrl_R plan | 0.32 |
| SSp-bfd_R rest vs VISrl_R plan | 0.74 |
| SSp-ll_L rest vs VISrl_R plan | 0.15 |
| SSp-ll_R rest vs VISrl_R plan | 0.61 |
| SSp-n_L rest vs VISrl_R plan | 0.63 |
| SSp-n_R rest vs VISrl_R plan | 0.98 |
| SSp-tr_L rest vs VISrl_R plan | 0.07 |
| SSp-tr_R rest vs VISrl_R plan | 0.26 |
| SSp-ul_L rest vs VISrl_R plan | 0.61 |
| SSp-ul_R rest vs VISrl_R plan | 0.84 |
| SSp-un_L rest vs VISrl_R plan | 0.71 |
| SSp-un_R rest vs VISrl_R plan | 0.94 |
| VISa_L rest vs VISrl_R plan | 0.05 |
| VISam_L rest vs VISrl_R plan | 0.014 |
| VISam_R rest vs VISrl_R plan | 0.025 |
| VISa_R rest vs VISrl_R plan | 0.077 |
| VISp_L rest vs VISrl_R plan | **0.0053** |
| VISpm_L rest vs VISrl_R plan | **0.0022** |
| VISpm_R rest vs VISrl_R plan | **0.0053** |
| VISp_R rest vs VISrl_R plan | 0.011 |
| VISrl_L rest vs VISrl_R plan | 0.038 |
| VISrl_R rest vs VISrl_R plan | 0.083 |

| | |
|---|---|
| CFA_R rest vs CFA_L rest | 1 |
| MOp_L rest vs CFA_L rest | 1 |
| MOp_R rest vs CFA_L rest | 1 |
| MOs_L rest vs CFA_L rest | 1 |
| MOs_R rest vs CFA_L rest | 1 |
| RFA_L rest vs CFA_L rest | 1 |
| RFA_R rest vs CFA_L rest | 1 |
| RSPagl_L rest vs CFA_L rest | 1 |
| RSPagl_R rest vs CFA_L rest | 1 |
| RSPd_L rest vs CFA_L rest | 1 |
| RSPd_R rest vs CFA_L rest | 1 |
| SSp-bfd_L rest vs CFA_L rest | 1 |
| SSp-bfd_R rest vs CFA_L rest | 1 |
| SSp-ll_L rest vs CFA_L rest | 1 |
| SSp-ll_R rest vs CFA_L rest | 1 |
| SSp-n_L rest vs CFA_L rest | 1 |
| SSp-n_R rest vs CFA_L rest | 1 |
| SSp-tr_L rest vs CFA_L rest | 1 |
| SSp-tr_R rest vs CFA_L rest | 1 |
| SSp-ul_L rest vs CFA_L rest | 1 |
| SSp-ul_R rest vs CFA_L rest | 1 |
| SSp-un_L rest vs CFA_L rest | 1 |
| SSp-un_R rest vs CFA_L rest | 1 |
| VISa_L rest vs CFA_L rest | 1 |
| VISam_L rest vs CFA_L rest | 1 |
| VISam_R rest vs CFA_L rest | 1 |
| VISa_R rest vs CFA_L rest | 1 |
| VISp_L rest vs CFA_L rest | 1 |
| VISpm_L rest vs CFA_L rest | 1 |
| VISpm_R rest vs CFA_L rest | 1 |
| VISp_R rest vs CFA_L rest | 1 |
| VISrl_L rest vs CFA_L rest | 1 |
| VISrl_R rest vs CFA_L rest | 1 |
| MOp_L rest vs CFA_R rest | 1 |
| MOp_R rest vs CFA_R rest | 1 |
| MOs_L rest vs CFA_R rest | 1 |
| MOs_R rest vs CFA_R rest | 1 |
| RFA_L rest vs CFA_R rest | 1 |
| RFA_R rest vs CFA_R rest | 1 |
| RSPagl_L rest vs CFA_R rest | 1 |
| RSPagl_R rest vs CFA_R rest | 1 |
| RSPd_L rest vs CFA_R rest | 1 |
| RSPd_R rest vs CFA_R rest | 1 |
| SSp-bfd_L rest vs CFA_R rest | 1 |
| SSp-bfd_R rest vs CFA_R rest | 1 |
| SSp-ll_L rest vs CFA_R rest | 1 |

| | |
|---|---|
| SSp-ll_R rest vs CFA_R rest | 1 |
| SSp-n_L rest vs CFA_R rest | 1 |
| SSp-n_R rest vs CFA_R rest | 1 |
| SSp-tr_L rest vs CFA_R rest | 1 |
| SSp-tr_R rest vs CFA_R rest | 1 |
| SSp-ul_L rest vs CFA_R rest | 1 |
| SSp-ul_R rest vs CFA_R rest | 1 |
| SSp-un_L rest vs CFA_R rest | 1 |
| SSp-un_R rest vs CFA_R rest | 1 |
| VISa_L rest vs CFA_R rest | 1 |
| VISam_L rest vs CFA_R rest | 1 |
| VISam_R rest vs CFA_R rest | 1 |
| VISa_R rest vs CFA_R rest | 1 |
| VISp_L rest vs CFA_R rest | 1 |
| VISpm_L rest vs CFA_R rest | 1 |
| VISpm_R rest vs CFA_R rest | 1 |
| VISp_R rest vs CFA_R rest | 1 |
| VISrl_L rest vs CFA_R rest | 1 |
| VISrl_R rest vs CFA_R rest | 1 |
| MOp_R rest vs MOp_L rest | 1 |
| MOs_L rest vs MOp_L rest | 1 |
| MOs_R rest vs MOp_L rest | 1 |
| RFA_L rest vs MOp_L rest | 1 |
| RFA_R rest vs MOp_L rest | 1 |
| RSPagl_L rest vs MOp_L rest | 1 |
| RSPagl_R rest vs MOp_L rest | 1 |
| RSPd_L rest vs MOp_L rest | 1 |
| RSPd_R rest vs MOp_L rest | 1 |
| SSp-bfd_L rest vs MOp_L rest | 1 |
| SSp-bfd_R rest vs MOp_L rest | 1 |
| SSp-ll_L rest vs MOp_L rest | 1 |
| SSp-ll_R rest vs MOp_L rest | 1 |
| SSp-n_L rest vs MOp_L rest | 1 |
| SSp-n_R rest vs MOp_L rest | 1 |
| SSp-tr_L rest vs MOp_L rest | 1 |
| SSp-tr_R rest vs MOp_L rest | 1 |
| SSp-ul_L rest vs MOp_L rest | 1 |
| SSp-ul_R rest vs MOp_L rest | 1 |
| SSp-un_L rest vs MOp_L rest | 1 |
| SSp-un_R rest vs MOp_L rest | 1 |
| VISa_L rest vs MOp_L rest | 1 |
| VISam_L rest vs MOp_L rest | 1 |
| VISam_R rest vs MOp_L rest | 1 |
| VISa_R rest vs MOp_L rest | 1 |
| VISp_L rest vs MOp_L rest | 1 |
| VISpm_L rest vs MOp_L rest | 1 |

| | |
|---|---|
| VISpm_R rest vs MOp_L rest | 1 |
| VISp_R rest vs MOp_L rest | 1 |
| VISrl_L rest vs MOp_L rest | 1 |
| VISrl_R rest vs MOp_L rest | 1 |
| MOs_L rest vs MOp_R rest | 1 |
| MOs_R rest vs MOp_R rest | 1 |
| RFA_L rest vs MOp_R rest | 1 |
| RFA_R rest vs MOp_R rest | 1 |
| RSPagl_L rest vs MOp_R rest | 1 |
| RSPagl_R rest vs MOp_R rest | 1 |
| RSPd_L rest vs MOp_R rest | 1 |
| RSPd_R rest vs MOp_R rest | 1 |
| SSp-bfd_L rest vs MOp_R rest | 1 |
| SSp-bfd_R rest vs MOp_R rest | 1 |
| SSp-ll_L rest vs MOp_R rest | 1 |
| SSp-ll_R rest vs MOp_R rest | 1 |
| SSp-n_L rest vs MOp_R rest | 1 |
| SSp-n_R rest vs MOp_R rest | 1 |
| SSp-tr_L rest vs MOp_R rest | 1 |
| SSp-tr_R rest vs MOp_R rest | 1 |
| SSp-ul_L rest vs MOp_R rest | 1 |
| SSp-ul_R rest vs MOp_R rest | 1 |
| SSp-un_L rest vs MOp_R rest | 1 |
| SSp-un_R rest vs MOp_R rest | 1 |
| VISa_L rest vs MOp_R rest | 1 |
| VISam_L rest vs MOp_R rest | 1 |
| VISam_R rest vs MOp_R rest | 1 |
| VISa_R rest vs MOp_R rest | 1 |
| VISp_L rest vs MOp_R rest | 1 |
| VISpm_L rest vs MOp_R rest | 1 |
| VISpm_R rest vs MOp_R rest | 1 |
| VISp_R rest vs MOp_R rest | 1 |
| VISrl_L rest vs MOp_R rest | 1 |
| VISrl_R rest vs MOp_R rest | 1 |
| MOs_R rest vs MOs_L rest | 1 |
| RFA_L rest vs MOs_L rest | 1 |
| RFA_R rest vs MOs_L rest | 1 |
| RSPagl_L rest vs MOs_L rest | 1 |
| RSPagl_R rest vs MOs_L rest | 1 |
| RSPd_L rest vs MOs_L rest | 1 |
| RSPd_R rest vs MOs_L rest | 1 |
| SSp-bfd_L rest vs MOs_L rest | 1 |
| SSp-bfd_R rest vs MOs_L rest | 1 |
| SSp-ll_L rest vs MOs_L rest | 1 |
| SSp-ll_R rest vs MOs_L rest | 1 |
| SSp-n_L rest vs MOs_L rest | 1 |

| | |
|---|---|
| SSp-n_R rest vs MOs_L rest | 1 |
| SSp-tr_L rest vs MOs_L rest | 1 |
| SSp-tr_R rest vs MOs_L rest | 1 |
| SSp-ul_L rest vs MOs_L rest | 1 |
| SSp-ul_R rest vs MOs_L rest | 1 |
| SSp-un_L rest vs MOs_L rest | 1 |
| SSp-un_R rest vs MOs_L rest | 1 |
| VISa_L rest vs MOs_L rest | 1 |
| VISam_L rest vs MOs_L rest | 1 |
| VISam_R rest vs MOs_L rest | 1 |
| VISa_R rest vs MOs_L rest | 1 |
| VISp_L rest vs MOs_L rest | 1 |
| VISpm_L rest vs MOs_L rest | 1 |
| VISpm_R rest vs MOs_L rest | 1 |
| VISp_R rest vs MOs_L rest | 1 |
| VISrl_L rest vs MOs_L rest | 1 |
| VISrl_R rest vs MOs_L rest | 1 |
| RFA_L rest vs MOs_R rest | 1 |
| RFA_R rest vs MOs_R rest | 1 |
| RSPagl_L rest vs MOs_R rest | 1 |
| RSPagl_R rest vs MOs_R rest | 1 |
| RSPd_L rest vs MOs_R rest | 1 |
| RSPd_R rest vs MOs_R rest | 1 |
| SSp-bfd_L rest vs MOs_R rest | 1 |
| SSp-bfd_R rest vs MOs_R rest | 1 |
| SSp-ll_L rest vs MOs_R rest | 1 |
| SSp-ll_R rest vs MOs_R rest | 1 |
| SSp-n_L rest vs MOs_R rest | 1 |
| SSp-n_R rest vs MOs_R rest | 1 |
| SSp-tr_L rest vs MOs_R rest | 1 |
| SSp-tr_R rest vs MOs_R rest | 1 |
| SSp-ul_L rest vs MOs_R rest | 1 |
| SSp-ul_R rest vs MOs_R rest | 1 |
| SSp-un_L rest vs MOs_R rest | 1 |
| SSp-un_R rest vs MOs_R rest | 1 |
| VISa_L rest vs MOs_R rest | 1 |
| VISam_L rest vs MOs_R rest | 1 |
| VISam_R rest vs MOs_R rest | 1 |
| VISa_R rest vs MOs_R rest | 1 |
| VISp_L rest vs MOs_R rest | 1 |
| VISpm_L rest vs MOs_R rest | 1 |
| VISpm_R rest vs MOs_R rest | 1 |
| VISp_R rest vs MOs_R rest | 1 |
| VISrl_L rest vs MOs_R rest | 1 |
| VISrl_R rest vs MOs_R rest | 1 |
| RFA_R rest vs RFA_L rest | 1 |

| | |
|---|---|
| RSPagl_L rest vs RFA_L rest | 1 |
| RSPagl_R rest vs RFA_L rest | 1 |
| RSPd_L rest vs RFA_L rest | 1 |
| RSPd_R rest vs RFA_L rest | 1 |
| SSp-bfd_L rest vs RFA_L rest | 1 |
| SSp-bfd_R rest vs RFA_L rest | 1 |
| SSp-ll_L rest vs RFA_L rest | 1 |
| SSp-ll_R rest vs RFA_L rest | 1 |
| SSp-n_L rest vs RFA_L rest | 1 |
| SSp-n_R rest vs RFA_L rest | 1 |
| SSp-tr_L rest vs RFA_L rest | 1 |
| SSp-tr_R rest vs RFA_L rest | 1 |
| SSp-ul_L rest vs RFA_L rest | 1 |
| SSp-ul_R rest vs RFA_L rest | 1 |
| SSp-un_L rest vs RFA_L rest | 1 |
| SSp-un_R rest vs RFA_L rest | 1 |
| VISa_L rest vs RFA_L rest | 1 |
| VISam_L rest vs RFA_L rest | 1 |
| VISam_R rest vs RFA_L rest | 1 |
| VISa_R rest vs RFA_L rest | 1 |
| VISp_L rest vs RFA_L rest | 1 |
| VISpm_L rest vs RFA_L rest | 1 |
| VISpm_R rest vs RFA_L rest | 1 |
| VISp_R rest vs RFA_L rest | 1 |
| VISrl_L rest vs RFA_L rest | 1 |
| VISrl_R rest vs RFA_L rest | 1 |
| RSPagl_L rest vs RFA_R rest | 1 |
| RSPagl_R rest vs RFA_R rest | 1 |
| RSPd_L rest vs RFA_R rest | 1 |
| RSPd_R rest vs RFA_R rest | 1 |
| SSp-bfd_L rest vs RFA_R rest | 1 |
| SSp-bfd_R rest vs RFA_R rest | 1 |
| SSp-ll_L rest vs RFA_R rest | 1 |
| SSp-ll_R rest vs RFA_R rest | 1 |
| SSp-n_L rest vs RFA_R rest | 1 |
| SSp-n_R rest vs RFA_R rest | 1 |
| SSp-tr_L rest vs RFA_R rest | 1 |
| SSp-tr_R rest vs RFA_R rest | 1 |
| SSp-ul_L rest vs RFA_R rest | 1 |
| SSp-ul_R rest vs RFA_R rest | 1 |
| SSp-un_L rest vs RFA_R rest | 1 |
| SSp-un_R rest vs RFA_R rest | 1 |
| VISa_L rest vs RFA_R rest | 1 |
| VISam_L rest vs RFA_R rest | 1 |
| VISam_R rest vs RFA_R rest | 1 |
| VISa_R rest vs RFA_R rest | 1 |

| | |
|---|---|
| VISp_L rest vs RFA_R rest | 1 |
| VISpm_L rest vs RFA_R rest | 1 |
| VISpm_R rest vs RFA_R rest | 1 |
| VISp_R rest vs RFA_R rest | 1 |
| VISrl_L rest vs RFA_R rest | 1 |
| VISrl_R rest vs RFA_R rest | 1 |
| RSPagl_R rest vs RSPagl_L rest | 1 |
| RSPd_L rest vs RSPagl_L rest | 1 |
| RSPd_R rest vs RSPagl_L rest | 1 |
| SSp-bfd_L rest vs RSPagl_L rest | 1 |
| SSp-bfd_R rest vs RSPagl_L rest | 1 |
| SSp-ll_L rest vs RSPagl_L rest | 1 |
| SSp-ll_R rest vs RSPagl_L rest | 1 |
| SSp-n_L rest vs RSPagl_L rest | 1 |
| SSp-n_R rest vs RSPagl_L rest | 1 |
| SSp-tr_L rest vs RSPagl_L rest | 1 |
| SSp-tr_R rest vs RSPagl_L rest | 1 |
| SSp-ul_L rest vs RSPagl_L rest | 1 |
| SSp-ul_R rest vs RSPagl_L rest | 1 |
| SSp-un_L rest vs RSPagl_L rest | 1 |
| SSp-un_R rest vs RSPagl_L rest | 1 |
| VISa_L rest vs RSPagl_L rest | 1 |
| VISam_L rest vs RSPagl_L rest | 1 |
| VISam_R rest vs RSPagl_L rest | 1 |
| VISa_R rest vs RSPagl_L rest | 1 |
| VISp_L rest vs RSPagl_L rest | 1 |
| VISpm_L rest vs RSPagl_L rest | 1 |
| VISpm_R rest vs RSPagl_L rest | 1 |
| VISp_R rest vs RSPagl_L rest | 1 |
| VISrl_L rest vs RSPagl_L rest | 1 |
| VISrl_R rest vs RSPagl_L rest | 1 |
| RSPd_L rest vs RSPagl_R rest | 1 |
| RSPd_R rest vs RSPagl_R rest | 1 |
| SSp-bfd_L rest vs RSPagl_R rest | 1 |
| SSp-bfd_R rest vs RSPagl_R rest | 1 |
| SSp-ll_L rest vs RSPagl_R rest | 1 |
| SSp-ll_R rest vs RSPagl_R rest | 1 |
| SSp-n_L rest vs RSPagl_R rest | 1 |
| SSp-n_R rest vs RSPagl_R rest | 1 |
| SSp-tr_L rest vs RSPagl_R rest | 1 |
| SSp-tr_R rest vs RSPagl_R rest | 1 |
| SSp-ul_L rest vs RSPagl_R rest | 1 |
| SSp-ul_R rest vs RSPagl_R rest | 1 |
| SSp-un_L rest vs RSPagl_R rest | 1 |
| SSp-un_R rest vs RSPagl_R rest | 1 |
| VISa_L rest vs RSPagl_R rest | 1 |

| | |
|---|---|
| VISam_L rest vs RSPagl_R rest | 1 |
| VISam_R rest vs RSPagl_R rest | 1 |
| VISa_R rest vs RSPagl_R rest | 1 |
| VISp_L rest vs RSPagl_R rest | 1 |
| VISpm_L rest vs RSPagl_R rest | 1 |
| VISpm_R rest vs RSPagl_R rest | 1 |
| VISp_R rest vs RSPagl_R rest | 1 |
| VISrl_L rest vs RSPagl_R rest | 1 |
| VISrl_R rest vs RSPagl_R rest | 1 |
| RSPd_R rest vs RSPd_L rest | 1 |
| SSp-bfd_L rest vs RSPd_L rest | 1 |
| SSp-bfd_R rest vs RSPd_L rest | 1 |
| SSp-ll_L rest vs RSPd_L rest | 1 |
| SSp-ll_R rest vs RSPd_L rest | 1 |
| SSp-n_L rest vs RSPd_L rest | 1 |
| SSp-n_R rest vs RSPd_L rest | 1 |
| SSp-tr_L rest vs RSPd_L rest | 1 |
| SSp-tr_R rest vs RSPd_L rest | 1 |
| SSp-ul_L rest vs RSPd_L rest | 1 |
| SSp-ul_R rest vs RSPd_L rest | 1 |
| SSp-un_L rest vs RSPd_L rest | 1 |
| SSp-un_R rest vs RSPd_L rest | 1 |
| VISa_L rest vs RSPd_L rest | 1 |
| VISam_L rest vs RSPd_L rest | 1 |
| VISam_R rest vs RSPd_L rest | 1 |
| VISa_R rest vs RSPd_L rest | 1 |
| VISp_L rest vs RSPd_L rest | 1 |
| VISpm_L rest vs RSPd_L rest | 1 |
| VISpm_R rest vs RSPd_L rest | 1 |
| VISp_R rest vs RSPd_L rest | 1 |
| VISrl_L rest vs RSPd_L rest | 1 |
| VISrl_R rest vs RSPd_L rest | 1 |
| SSp-bfd_L rest vs RSPd_R rest | 1 |
| SSp-bfd_R rest vs RSPd_R rest | 1 |
| SSp-ll_L rest vs RSPd_R rest | 1 |
| SSp-ll_R rest vs RSPd_R rest | 1 |
| SSp-n_L rest vs RSPd_R rest | 1 |
| SSp-n_R rest vs RSPd_R rest | 1 |
| SSp-tr_L rest vs RSPd_R rest | 1 |
| SSp-tr_R rest vs RSPd_R rest | 1 |
| SSp-ul_L rest vs RSPd_R rest | 1 |
| SSp-ul_R rest vs RSPd_R rest | 1 |
| SSp-un_L rest vs RSPd_R rest | 1 |
| SSp-un_R rest vs RSPd_R rest | 1 |
| VISa_L rest vs RSPd_R rest | 1 |
| VISam_L rest vs RSPd_R rest | 1 |

| | |
|---|---|
| VISam_R rest vs RSPd_R rest | 1 |
| VISa_R rest vs RSPd_R rest | 1 |
| VISp_L rest vs RSPd_R rest | 1 |
| VISpm_L rest vs RSPd_R rest | 1 |
| VISpm_R rest vs RSPd_R rest | 1 |
| VISp_R rest vs RSPd_R rest | 1 |
| VISrl_L rest vs RSPd_R rest | 1 |
| VISrl_R rest vs RSPd_R rest | 1 |
| SSp-bfd_R rest vs SSp-bfd_L rest | 1 |
| SSp-ll_L rest vs SSp-bfd_L rest | 1 |
| SSp-ll_R rest vs SSp-bfd_L rest | 1 |
| SSp-n_L rest vs SSp-bfd_L rest | 1 |
| SSp-n_R rest vs SSp-bfd_L rest | 1 |
| SSp-tr_L rest vs SSp-bfd_L rest | 1 |
| SSp-tr_R rest vs SSp-bfd_L rest | 1 |
| SSp-ul_L rest vs SSp-bfd_L rest | 1 |
| SSp-ul_R rest vs SSp-bfd_L rest | 1 |
| SSp-un_L rest vs SSp-bfd_L rest | 1 |
| SSp-un_R rest vs SSp-bfd_L rest | 1 |
| VISa_L rest vs SSp-bfd_L rest | 1 |
| VISam_L rest vs SSp-bfd_L rest | 1 |
| VISam_R rest vs SSp-bfd_L rest | 1 |
| VISa_R rest vs SSp-bfd_L rest | 1 |
| VISp_L rest vs SSp-bfd_L rest | 1 |
| VISpm_L rest vs SSp-bfd_L rest | 1 |
| VISpm_R rest vs SSp-bfd_L rest | 1 |
| VISp_R rest vs SSp-bfd_L rest | 1 |
| VISrl_L rest vs SSp-bfd_L rest | 1 |
| VISrl_R rest vs SSp-bfd_L rest | 1 |
| SSp-ll_L rest vs SSp-bfd_R rest | 1 |
| SSp-ll_R rest vs SSp-bfd_R rest | 1 |
| SSp-n_L rest vs SSp-bfd_R rest | 1 |
| SSp-n_R rest vs SSp-bfd_R rest | 1 |
| SSp-tr_L rest vs SSp-bfd_R rest | 1 |
| SSp-tr_R rest vs SSp-bfd_R rest | 1 |
| SSp-ul_L rest vs SSp-bfd_R rest | 1 |
| SSp-ul_R rest vs SSp-bfd_R rest | 1 |
| SSp-un_L rest vs SSp-bfd_R rest | 1 |
| SSp-un_R rest vs SSp-bfd_R rest | 1 |
| VISa_L rest vs SSp-bfd_R rest | 1 |
| VISam_L rest vs SSp-bfd_R rest | 1 |
| VISam_R rest vs SSp-bfd_R rest | 1 |
| VISa_R rest vs SSp-bfd_R rest | 1 |
| VISp_L rest vs SSp-bfd_R rest | 1 |
| VISpm_L rest vs SSp-bfd_R rest | 1 |
| VISpm_R rest vs SSp-bfd_R rest | 1 |

| | |
|---|---|
| VISp_R rest vs SSp-bfd_R rest | 1 |
| VISrl_L rest vs SSp-bfd_R rest | 1 |
| VISrl_R rest vs SSp-bfd_R rest | 1 |
| SSp-ll_R rest vs SSp-ll_L rest | 1 |
| SSp-n_L rest vs SSp-ll_L rest | 1 |
| SSp-n_R rest vs SSp-ll_L rest | 1 |
| SSp-tr_L rest vs SSp-ll_L rest | 1 |
| SSp-tr_R rest vs SSp-ll_L rest | 1 |
| SSp-ul_L rest vs SSp-ll_L rest | 1 |
| SSp-ul_R rest vs SSp-ll_L rest | 1 |
| SSp-un_L rest vs SSp-ll_L rest | 1 |
| SSp-un_R rest vs SSp-ll_L rest | 1 |
| VISa_L rest vs SSp-ll_L rest | 1 |
| VISam_L rest vs SSp-ll_L rest | 1 |
| VISam_R rest vs SSp-ll_L rest | 1 |
| VISa_R rest vs SSp-ll_L rest | 1 |
| VISp_L rest vs SSp-ll_L rest | 1 |
| VISpm_L rest vs SSp-ll_L rest | 1 |
| VISpm_R rest vs SSp-ll_L rest | 1 |
| VISp_R rest vs SSp-ll_L rest | 1 |
| VISrl_L rest vs SSp-ll_L rest | 1 |
| VISrl_R rest vs SSp-ll_L rest | 1 |
| SSp-n_L rest vs SSp-ll_R rest | 1 |
| SSp-n_R rest vs SSp-ll_R rest | 1 |
| SSp-tr_L rest vs SSp-ll_R rest | 1 |
| SSp-tr_R rest vs SSp-ll_R rest | 1 |
| SSp-ul_L rest vs SSp-ll_R rest | 1 |
| SSp-ul_R rest vs SSp-ll_R rest | 1 |
| SSp-un_L rest vs SSp-ll_R rest | 1 |
| SSp-un_R rest vs SSp-ll_R rest | 1 |
| VISa_L rest vs SSp-ll_R rest | 1 |
| VISam_L rest vs SSp-ll_R rest | 1 |
| VISam_R rest vs SSp-ll_R rest | 1 |
| VISa_R rest vs SSp-ll_R rest | 1 |
| VISp_L rest vs SSp-ll_R rest | 1 |
| VISpm_L rest vs SSp-ll_R rest | 1 |
| VISpm_R rest vs SSp-ll_R rest | 1 |
| VISp_R rest vs SSp-ll_R rest | 1 |
| VISrl_L rest vs SSp-ll_R rest | 1 |
| VISrl_R rest vs SSp-ll_R rest | 1 |
| SSp-n_R rest vs SSp-n_L rest | 1 |
| SSp-tr_L rest vs SSp-n_L rest | 1 |
| SSp-tr_R rest vs SSp-n_L rest | 1 |
| SSp-ul_L rest vs SSp-n_L rest | 1 |
| SSp-ul_R rest vs SSp-n_L rest | 1 |
| SSp-un_L rest vs SSp-n_L rest | 1 |

| | |
|---|---|
| SSp-un_R rest vs SSp-n_L rest | 1 |
| VISa_L rest vs SSp-n_L rest | 1 |
| VISam_L rest vs SSp-n_L rest | 1 |
| VISam_R rest vs SSp-n_L rest | 1 |
| VISa_R rest vs SSp-n_L rest | 1 |
| VISp_L rest vs SSp-n_L rest | 1 |
| VISpm_L rest vs SSp-n_L rest | 1 |
| VISpm_R rest vs SSp-n_L rest | 1 |
| VISp_R rest vs SSp-n_L rest | 1 |
| VISrl_L rest vs SSp-n_L rest | 1 |
| VISrl_R rest vs SSp-n_L rest | 1 |
| SSp-tr_L rest vs SSp-n_R rest | 1 |
| SSp-tr_R rest vs SSp-n_R rest | 1 |
| SSp-ul_L rest vs SSp-n_R rest | 1 |
| SSp-ul_R rest vs SSp-n_R rest | 1 |
| SSp-un_L rest vs SSp-n_R rest | 1 |
| SSp-un_R rest vs SSp-n_R rest | 1 |
| VISa_L rest vs SSp-n_R rest | 1 |
| VISam_L rest vs SSp-n_R rest | 1 |
| VISam_R rest vs SSp-n_R rest | 1 |
| VISa_R rest vs SSp-n_R rest | 1 |
| VISp_L rest vs SSp-n_R rest | 1 |
| VISpm_L rest vs SSp-n_R rest | 1 |
| VISpm_R rest vs SSp-n_R rest | 1 |
| VISp_R rest vs SSp-n_R rest | 1 |
| VISrl_L rest vs SSp-n_R rest | 1 |
| VISrl_R rest vs SSp-n_R rest | 1 |
| SSp-tr_R rest vs SSp-tr_L rest | 1 |
| SSp-ul_L rest vs SSp-tr_L rest | 1 |
| SSp-ul_R rest vs SSp-tr_L rest | 1 |
| SSp-un_L rest vs SSp-tr_L rest | 1 |
| SSp-un_R rest vs SSp-tr_L rest | 1 |
| VISa_L rest vs SSp-tr_L rest | 1 |
| VISam_L rest vs SSp-tr_L rest | 1 |
| VISam_R rest vs SSp-tr_L rest | 1 |
| VISa_R rest vs SSp-tr_L rest | 1 |
| VISp_L rest vs SSp-tr_L rest | 1 |
| VISpm_L rest vs SSp-tr_L rest | 1 |
| VISpm_R rest vs SSp-tr_L rest | 1 |
| VISp_R rest vs SSp-tr_L rest | 1 |
| VISrl_L rest vs SSp-tr_L rest | 1 |
| VISrl_R rest vs SSp-tr_L rest | 1 |
| SSp-ul_L rest vs SSp-tr_R rest | 1 |
| SSp-ul_R rest vs SSp-tr_R rest | 1 |
| SSp-un_L rest vs SSp-tr_R rest | 1 |
| SSp-un_R rest vs SSp-tr_R rest | 1 |

| | |
|---|---|
| VISa_L rest vs SSp-tr_R rest | 1 |
| VISam_L rest vs SSp-tr_R rest | 1 |
| VISam_R rest vs SSp-tr_R rest | 1 |
| VISa_R rest vs SSp-tr_R rest | 1 |
| VISp_L rest vs SSp-tr_R rest | 1 |
| VISpm_L rest vs SSp-tr_R rest | 1 |
| VISpm_R rest vs SSp-tr_R rest | 1 |
| VISp_R rest vs SSp-tr_R rest | 1 |
| VISrl_L rest vs SSp-tr_R rest | 1 |
| VISrl_R rest vs SSp-tr_R rest | 1 |
| SSp-ul_R rest vs SSp-ul_L rest | 1 |
| SSp-un_L rest vs SSp-ul_L rest | 1 |
| SSp-un_R rest vs SSp-ul_L rest | 1 |
| VISa_L rest vs SSp-ul_L rest | 1 |
| VISam_L rest vs SSp-ul_L rest | 1 |
| VISam_R rest vs SSp-ul_L rest | 1 |
| VISa_R rest vs SSp-ul_L rest | 1 |
| VISp_L rest vs SSp-ul_L rest | 1 |
| VISpm_L rest vs SSp-ul_L rest | 1 |
| VISpm_R rest vs SSp-ul_L rest | 1 |
| VISp_R rest vs SSp-ul_L rest | 1 |
| VISrl_L rest vs SSp-ul_L rest | 1 |
| VISrl_R rest vs SSp-ul_L rest | 1 |
| SSp-un_L rest vs SSp-ul_R rest | 1 |
| SSp-un_R rest vs SSp-ul_R rest | 1 |
| VISa_L rest vs SSp-ul_R rest | 1 |
| VISam_L rest vs SSp-ul_R rest | 1 |
| VISam_R rest vs SSp-ul_R rest | 1 |
| VISa_R rest vs SSp-ul_R rest | 1 |
| VISp_L rest vs SSp-ul_R rest | 1 |
| VISpm_L rest vs SSp-ul_R rest | 1 |
| VISpm_R rest vs SSp-ul_R rest | 1 |
| VISp_R rest vs SSp-ul_R rest | 1 |
| VISrl_L rest vs SSp-ul_R rest | 1 |
| VISrl_R rest vs SSp-ul_R rest | 1 |
| SSp-un_R rest vs SSp-un_L rest | 1 |
| VISa_L rest vs SSp-un_L rest | 1 |
| VISam_L rest vs SSp-un_L rest | 1 |
| VISam_R rest vs SSp-un_L rest | 1 |
| VISa_R rest vs SSp-un_L rest | 1 |
| VISp_L rest vs SSp-un_L rest | 1 |
| VISpm_L rest vs SSp-un_L rest | 1 |
| VISpm_R rest vs SSp-un_L rest | 1 |
| VISp_R rest vs SSp-un_L rest | 1 |
| VISrl_L rest vs SSp-un_L rest | 1 |
| VISrl_R rest vs SSp-un_L rest | 1 |

| | |
|---|---|
| VISa_L rest vs SSp-un_R rest | 1 |
| VISam_L rest vs SSp-un_R rest | 1 |
| VISam_R rest vs SSp-un_R rest | 1 |
| VISa_R rest vs SSp-un_R rest | 1 |
| VISp_L rest vs SSp-un_R rest | 1 |
| VISpm_L rest vs SSp-un_R rest | 1 |
| VISpm_R rest vs SSp-un_R rest | 1 |
| VISp_R rest vs SSp-un_R rest | 1 |
| VISrl_L rest vs SSp-un_R rest | 1 |
| VISrl_R rest vs SSp-un_R rest | 1 |
| VISam_L rest vs VISa_L rest | 1 |
| VISam_R rest vs VISa_L rest | 1 |
| VISa_R rest vs VISa_L rest | 1 |
| VISp_L rest vs VISa_L rest | 1 |
| VISpm_L rest vs VISa_L rest | 1 |
| VISpm_R rest vs VISa_L rest | 1 |
| VISp_R rest vs VISa_L rest | 1 |
| VISrl_L rest vs VISa_L rest | 1 |
| VISrl_R rest vs VISa_L rest | 1 |
| VISam_R rest vs VISam_L rest | 1 |
| VISa_R rest vs VISam_L rest | 1 |
| VISp_L rest vs VISam_L rest | 1 |
| VISpm_L rest vs VISam_L rest | 1 |
| VISpm_R rest vs VISam_L rest | 1 |
| VISp_R rest vs VISam_L rest | 1 |
| VISrl_L rest vs VISam_L rest | 1 |
| VISrl_R rest vs VISam_L rest | 1 |
| VISa_R rest vs VISam_R rest | 1 |
| VISp_L rest vs VISam_R rest | 1 |
| VISpm_L rest vs VISam_R rest | 1 |
| VISpm_R rest vs VISam_R rest | 1 |
| VISp_R rest vs VISam_R rest | 1 |
| VISrl_L rest vs VISam_R rest | 1 |
| VISrl_R rest vs VISam_R rest | 1 |
| VISp_L rest vs VISa_R rest | 1 |
| VISpm_L rest vs VISa_R rest | 1 |
| VISpm_R rest vs VISa_R rest | 1 |
| VISp_R rest vs VISa_R rest | 1 |
| VISrl_L rest vs VISa_R rest | 1 |
| VISrl_R rest vs VISa_R rest | 1 |
| VISpm_L rest vs VISp_L rest | 1 |
| VISpm_R rest vs VISp_L rest | 1 |
| VISp_R rest vs VISp_L rest | 1 |
| VISrl_L rest vs VISp_L rest | 1 |
| VISrl_R rest vs VISp_L rest | 1 |
| VISpm_R rest vs VISpm_L rest | 1 |

| | |
|---|---|
| VISp_R rest vs VISpm_L rest | 1 |
| VISrl_L rest vs VISpm_L rest | 1 |
| VISrl_R rest vs VISpm_L rest | 1 |
| VISp_R rest vs VISpm_R rest | 1 |
| VISrl_L rest vs VISpm_R rest | 1 |
| VISrl_R rest vs VISpm_R rest | 1 |
| VISrl_L rest vs VISp_R rest | 1 |
| VISrl_R rest vs VISp_R rest | 1 |
| VISrl_R rest vs VISrl_L rest | 1 |
